# The evolutionary path of the epithelial sodium channel δ-subunit in Cetartiodactyla points to a role in sodium sensing

**DOI:** 10.1101/2024.11.18.623996

**Authors:** Fynn Zahnow, Chiara Jäger, Yassmin Mohamed, Gianluca Vogelhuber, Fabian May, Alexandra Maria Ciocan, Arianna Manieri, Stephan Maxeiner, Gabriela Krasteva-Christ, Matthew R. D. Cobain, Lars Podsiadlowski, José Luis Crespo-Picazo, Daniel García-Párraga, Mike Althaus

## Abstract

The epithelial sodium channel (ENaC) plays a key role in osmoregulation in tetrapod vertebrates and is a candidate receptor for salt taste sensation. There are four ENaC subunits (α, β, γ, δ) which form αβγ- or δβγ ENaCs. While αβγ-ENaC is a ‘maintenance protein’ controlling sodium and potassium homeostasis, δβγ-ENaC might represent a ‘stress protein’ monitoring high sodium concentrations. The δ-subunit emerged with water-to-land transition of tetrapod vertebrate ancestors. We investigated the evolutionary path of ENaC-coding genes in Cetartiodactyla, a group comprising even-toed ungulates and the cetaceans (whales/dolphins) which transitioned from terrestrial to marine environments in the Eocene. The genes *SCNN1A* (α-ENaC), *SCNN1B* (β-ENaC) and *SCNN1G* (γ-ENaC) are intact in all 22 investigated cetartiodactylan families. While *SCNN1D* (δ-ENaC) is intact in terrestrial Artiodactyla, it is a pseudogene in 12 cetacean families. A fusion of *SCNN1D* exons 11 and 12 under preservation of the open reading frame was observed in the Antilopinae, representing a new feature of this clade. Transcripts of *SCNN1A*, *SCNN1B* and *SCNN1G* were present in kidney and lung tissues of Bottlenose dolphins, highlighting αβγ-ENaC’s role as a maintenance protein. Consistent with *SCNN1D* loss, Bottlenose dolphins and Beluga whales did not show behavioural differences to stimuli with or without sodium in seawater-equivalent concentrations. These data suggest a function of δ-ENaC as a sodium sensing protein which might have become obsolete in cetaceans after the migration to high-salinity marine environments. Consistently, there is reduced selection pressure or pseudogenisation of *SCNN1D* in other marine mammals, including sirenians, pinnipeds and sea otter.

## Introduction

Transitions between terrestrial and aquatic environments were key events in mammalian evolution. Extant cetaceans (whales and dolphins) evolved from terrestrial artiodactyls (even-toed ungulates) which transitioned into the oceans in the Eocene (Thewissen et al. 2007). This transition required phenotypic adaptations to marine environments and the loss or pseudogenisation of protein-coding genes is a frequent genomic change observed in cetaceans (Sharma et al. 2018). For example, gene-loss has been described for hair- and epidermis-specific genes (Sharma et al. 2018), enamel-specific genes (Randall et al. 2024), genes involved in the control of haemoglobin oxygen affinity (Sharma et al. 2018), lipid metabolism and/or oxidative stress response (Meyer et al. 2018) or taste receptor genes (Feng et al. 2014; Zhu et al. 2014). While some genes may simply become less relevant when species transition into new environments, gene loss or pseudogenisation has also been suggested to trigger broader phenotypic changes (Graham et al. 2023).

The transition of cetacean ancestors to marine habitats was accompanied by major dietary changes from herbivory/omnivory to carnivory (Thewissen et al. 2007) as well as the evolution of mechanisms to maintain electrolyte homeostasis in a hypersaline marine environment (Thewissen et al. 1996). While terrestrial herbivorous artiodactyls are challenged with maintaining stable sodium concentrations in their extracellular fluids in a sodium-scarce environment, marine cetaceans are faced with environmental sodium concentrations >3 times greater than their plasma sodium levels (Ewing et al. 2017) and little to no access to freshwater.

A key protein in the control of sodium homeostasis in mammals is the epithelial sodium channel (ENaC) (Rotin and Staub 2021). Canonical ENaC is a heterotrimeric sodium-selective ion channel which is composed of three subunits encoded by the genes *SCNN1A* (α-ENaC subunit), *SCNN1B* (β-ENaC subunit) and *SCNN1G* (γ-ENaC subunit). αβγ-ENaC is located in the apical membranes of epithelial cells in the distal nephrons of the kidneys, lungs and colon (Rotin and Staub 2021). In the distal nephron, aldosterone enhances ENaC expression and thereby matches dietary sodium intake with its excretion rate. In rodents, αβγ-ENaC is also expressed in magnocellular neurons of the paraventricular nucleus and supraoptic nucleus of the hypothalamus and might serve as a sodium sensor in the control of vasopressin release (Teruyama et al. 2012). Consistent with sensory functions of this ion channel, αβγ-ENaC has been shown to mediate appetitive sodium taste in mice (Chandrashekar et al. 2010; Nomura et al. 2020).

An additional *SCNN1D* gene, encoding the δ-ENaC subunit, emerged with the water-to-land transition in tetrapod vertebrates (Wichmann and Althaus 2020; Wang et al. 2022). This subunit can assemble with β- and γ-ENaC, yielding δβγ-ENaCs with distinct biophysical properties and mechanisms regulating its function. δβγ-ENaCs consistently generate larger transmembrane ion currents in heterologous expression systems than αβγ-ENaCs (Haerteis et al. 2009; Wichmann et al. 2018; Gettings et al. 2021). ENaC activity is rapidly controlled by an auto-regulatory mechanism termed sodium self-inhibition (SSI). SSI is triggered by binding of sodium ions to a cation binding site formed by amino acid residues in the extracellular β6-β7-loop of the α-ENaC or δ-ENaC subunits (Kashlan et al. 2015; Wichmann et al. 2019; Noreng et al. 2020), which subsequently reduces channel open probability. It is hypothesised that SSI rapidly adjusts ENaC open probability to fluctuating extracellular sodium concentrations, particularly in the distal nephron where this mechanism might prevent excess sodium absorption when urinary sodium concentrations are high (Kleyman et al. 2018). However, SSI of δβγ-ENaC is species-dependent, with mammalian δβγ-ENaCs showing a markedly reduced SSI due to lack of key structural motifs forming the sodium binding site in the extracellular domain of the δ-ENaC subunit (Wichmann et al. 2019; Gettings et al. 2021). Due to SSI, activity of mammalian αβγ-ENaC does not increase under extracellular sodium concentrations higher than plasma sodium concentrations, whereas mammalian δβγ-ENaC operates over broader sodium concentration ranges in heterologous expression systems (Gettings et al. 2021). Consistently, αβγ-ENaC mediates attractive taste to low sodium concentrations in mice (Chandrashekar et al. 2010; Nomura et al. 2020). In humans, it has been hypothesized that δβγ-ENaC in taste buds could sense high sodium concentrations that exceed plasma levels (Bigiani 2020). While αβγ-ENaC is a key ‘maintenance protein’ controlling overall body sodium homeostasis, δβγ-ENaC likely represents a ‘stress protein’ monitoring high sodium concentrations in tetrapod vertebrates. We, therefore, questioned how ENaC isoforms might operate in hypersaline marine environments and investigated the evolutionary path of ENaC subunit coding genes in cetaceans and terrestrial Artiodactyla.

## Results and Discussion

Using genomic sequences available in the National Center for Biotechnology Information (NCBI) database, we analysed all four *SCNN1* genes in 45 representative species of 22 cetartiodactylan families. To facilitate inter-species comparisons, exon nomenclature was followed as previously suggested (Gettings et al. 2021), with exons 2 and 13 encoding the first and second transmembrane domain of each ENaC subunit, respectively. In most species, exon 2 contains the translation start codon (with exception of some *SCNN1D* genes in terrestrial Artiodactyla where the start codon is located on an upstream exon 1, **Supplemental Data 1**, **Source Data 1**), while the STOP codon is located on exon 13. Thus, exons 2-13 encode the highly conserved core region of each ENaC subunit. Additional exons upstream of exon 2 are more variable and might be subject to alternative splicing. In this study, we, therefore, concentrated on the core region of each ENaC subunit and analysed the overall gene structure, exon sizes, splice donor/acceptor sites and continuous open reading frames (ORFs), in comparison with the corresponding sequences of the Alpaca (*Vicugna pacos*) as a reference. Because multiple NCBI mRNA sequences were predicted by algorithms and contain corrections (including removal of insertions/deletions and STOP codons), we constructed all sequences using genomic DNA of each respective species as template (**Source Data 1**).

The structures of the *SCNN1A*, *SCNN1B* and *SCNN1G* genes were highly conserved in the analysed cetacean and terrestrial artiodactylan species (**Figure 1, Supplemental Data 1, Source Data 1**). However, we observed clade-specific gene alterations. For example, in *SCNN1A* we detected a micro-deletion of 3 bases under preservation of the reading frame in exon 4 of the Odontoceti (toothed whales), with exception of the Indian river dolphins (Platanistidae), and sperm whales (Kogiidae and Physeteridae). The micro-deletion of the 3 bases in the river dolphin families Pontoporiidae, Iniidae and Ziphiidae, but not the Platanistidae, is consistent with the hypothesis that Platanistidae are an early diverging group of the Odontoceti, distant from the other river dolphin families (de Muizon et al. 2018; Benites-Palomino et al. 2024) (**Figure 2**). By contrast, exon 4 carried an insertion of 3 bases in the Ruminantia (**Figure 1, Supplemental Data 1**). All analysed Cetartiodactyla contained a rare non-canonical GC splice donor motif (Burset et al. 2001) flanking exon 6 (**Figure 1, Supplemental Data 1**). Nevertheless, the analysed *SCNN1A*, *SCNN1B* and *SCNN1G* sequences yielded continuous open reading frames (ORFs) (**Supplemental Data 2**). An alignment of the obtained amino acid sequences to human orthologues revealed high similarities, including: (i) the presence of key structural and regulatory sequence motifs such as an N-terminally located HG-motif impacting channel open probability, (ii) a conserved cation binding site initiating SSI in the α-ENaC subunit (*SCNN1A*), (iii) furin cleavage sites in the α-ENaC (*SCNN1A*) and γ-ENaC (*SCNN1G*) subunits, (iv) key cysteines involved in providing tertiary structure, (v) a GSS/A motif playing a role in ion size discrimination, as well as (vi) C-terminally located PPPxY motifs controlling ENaC membrane abundance (Rotin and Staub 2021; Althaus et al. 2023). We confirmed high degrees of conservation in all core structural elements of each subunit (**Supplemental Data 3**) by projecting the degree of sequence conservation based on the amino acid alignments (**Supplemental Data 2**) onto the Cryo-EM-derived structure for human ENaC subunits (Noreng et al. 2018; Noreng et al. 2020). These data clearly demonstrate the presence of functional genes encoding canonical αβγ-ENaC in the analysed Cetartiodactyla (**Figure 2**).

**Figure 1.**
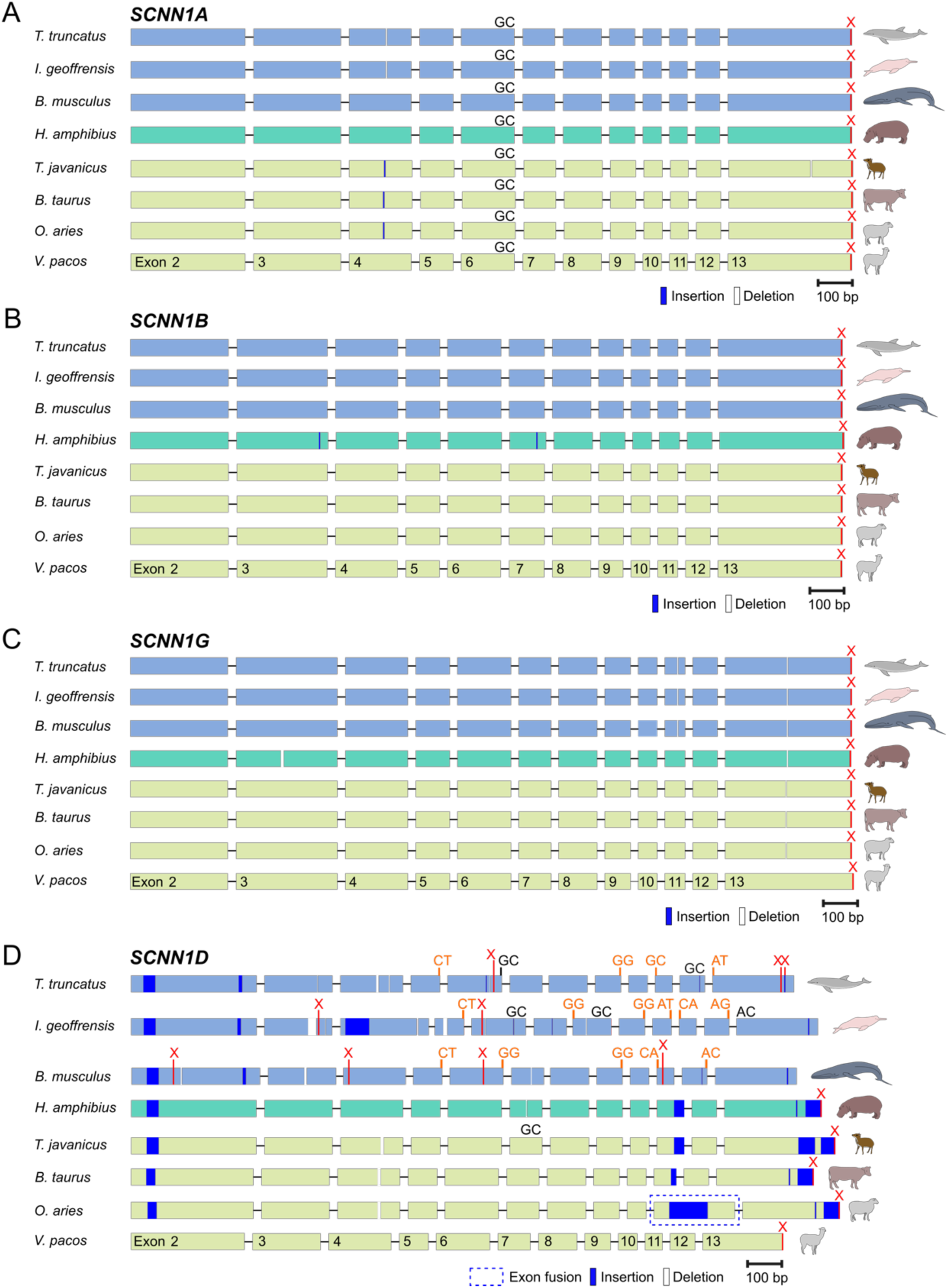
Structure of *SCNN1* genes in select Cetartiodactyla. Panels A-D show the genes *SCNN1A*, *SCNN1B*, *SCNN1G* and *SCNN1D*, respectively. Exons 2-13 are numbered according to the nomenclature by (Gettings et al. 2021) with exons 2 and 13 encoding the first and second transmembrane domain of each ENaC subunit, respectively. Only exons are drawn to scale. Exon sizes are outlined with respect to the Alpaca (*Vicugna pacos*) as a reference and insertions and deletions are highlighted. The size of exon 13 is shown up to the corresponding STOP codon in *V. pacos*. Non-canonical splice donor/acceptor motifs (AC/GC) flanking the exons are indicated in black font, whereas mutated splice sites are indicated in orange. Stop codons are marked with a red X. The dashed box indicates a fusion of exons 11 and 12 of the *SCNN1D* gene due to mutation of the splice donor following exon 11 in *Ovis aries*.

**Figure 2.**
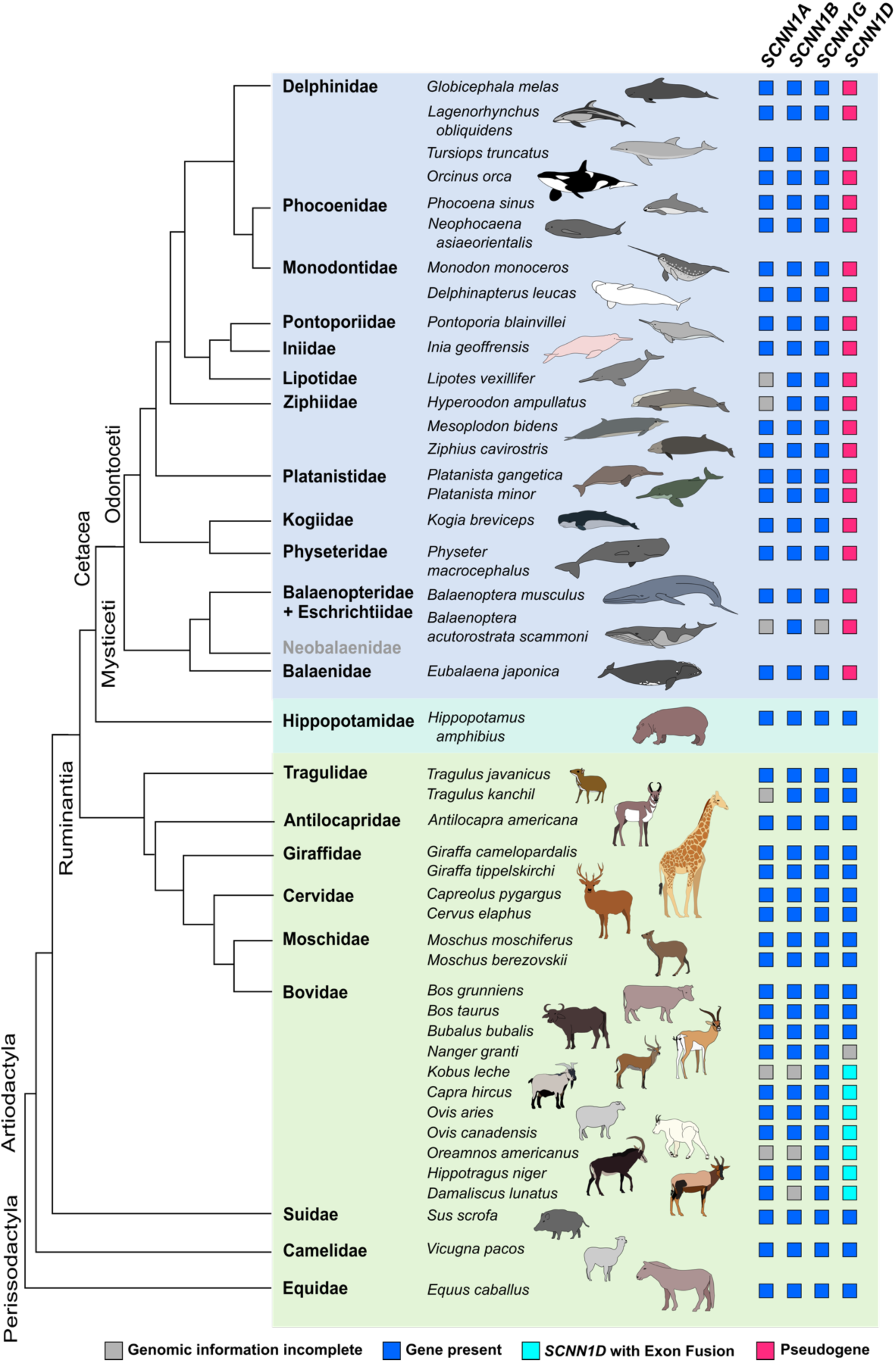
Presence of ENaC-coding *SCNN1* genes in Cetartiodactyla. Presence or pseudogenisation of *SCNN1* genes was investigated in the highlighted species using the National Center for Biotechnology Information (NCBI) database. Blue squares indicate intact genes yielding continuous open reading frames (ORFs), whereas magenta squares indicate pseudogenes containing multiple insertions/deletions and splice donor/acceptor site mutations causing preliminary STOP codons. Turquoise squares highlight species containing a fusion of exons 11 and 12 due to a mutation of the splice donor following exon 11, which did not disrupt the ORF. Gray squares highlight incomplete genomic information. There is currently no available genomic information of species representing the family Neobalaenidae. The phylogenetic tree was created based on (Cabrera et al. 2021).

In contrast to the *SCNN1A*, *SCNN1B* and *SCNN1G* genes, the *SCNN1D* gene encoding the δ-ENaC subunit displayed significant differences and clade-specific structural changes (**Figure 1, Supplemental Data 1, Source Data 1**). Consistent with our previous analysis of the *SCNN1D* gene in rodents (Gettings et al. 2021), the gene is condensed due to smaller intron sizes, in comparison to the *SCNN1A*, *SCNN1B* and *SCNN1G* genes (**Supplemental Data 1, Source Data 1**). The *SCNN1D* gene contains multiple insertions/deletions and mutations of splice donor/acceptor sites in all analysed cetacean species, causing frame-shifts and premature STOP codons, thereby disrupting the ORFs. A comprehensive map of these alterations is provided in **Supplemental Data 1**. By contrast, the *SCNN1D* genes were intact and yielded continuous ORFs in the investigated terrestrial Artiodactyla (**Figure 1, Supplemental Data 2**). The presence of a presumably functional *SCNN1D* gene in the Hippopotamus (*H. amphibius*) suggests that *SCNN1D* pseudogenised early in the evolution of Cetacea (**Figure 1**, **Figure 2, Supplemental Data 1, Source Data 1**). It is a pseudogene in both the toothed whales (Odontoceti) and baleen whales (Mysticeti), suggesting that loss of a functional *SCNN1D* gene is a general feature of cetaceans since more than 34.5 million years (Fordyce 2003; Lambert et al. 2017; Tsai et al. 2024). All investigated cetaceans share an identical mutated splice donor following exon 5 and share mutations in the splice donor following exon 9, as well as the splice acceptor ahead of exon 11 (**Supplemental Data 1**). This suggests that a single pseudogenisation event may have happened at the base of cetacean origin and additional inactivating mutations and insertions/deletions accumulated later in the different cetacean families.

We observed a fusion of exons 11 and 12 of the *SCNN1D* gene due to mutation of splice donor sites in the Bovidae (**Figure 1**, **Figure 2**, **Figure 3 A, B, Supplemental Data 1, Source Data 1**). This exon fusion causes incorporation of the former intron separating exons 11 and 12 into the coding mRNA sequences but does not disrupt the ORFs. *SCNN1D* transcripts have previously been detected in the digestive tract of sheep (*Ovis aries*) (Chen et al. 2019). We, therefore, isolated total mRNA of *O. aries* digestive tract tissues and confirmed the presence of the *SCNN1D* exon fusion using RT-PCR and sequencing of the PCR products (**Figure 3 A, B**). Transcripts of *SCNN1D* were also detected in *O. aries* tongue tissue (**Figure 3 C**). Interestingly, we previously observed the same exon 11/12 fusion in *SCNN1D* in the rodent infraorder Hystricognathi (Gettings et al. 2021), suggesting that this genetic alteration evolved at least twice and independently in mammals. To further investigate the evolutionary origin of the exon 11/12 fusion in the Bovidae, we analysed 19 representative species of 12 families (**Figure 3 D**): The Boselaphini (*B. tragocamelus*), Tragelaphini (*T. eurycerus*) and Bovini (*B. grunniens, B. taurus, B. bubalis*), grouped as the Bovinae, and the Aepycerotini (*A. melampus*), Neotragini (*N. pygmaeus*), Antilopini (*N. dama, M. kirkii*), Reduncini (*K. leche*), Oreotragini (*O. oreotragus*), Cephalophini (*C. harveyi*), Alcelaphini (*D. lunatus, C. taurinus*), Hippotragini (*H. niger*) and Caprini (*C. hircus, O. aries, O. canadensis, O. americanus*), grouped as the Antilopinae (Hassanin et al. 2012). The exon fusion was present in all investigated Antilopinae but absent in the Bovinae. The exon fusion can therefore be considered a new autapomorphic feature of the Antilopinae and is a feature of this clade since at least 16 million years (Timetree, accessed 12/2023).

**Figure 3.**
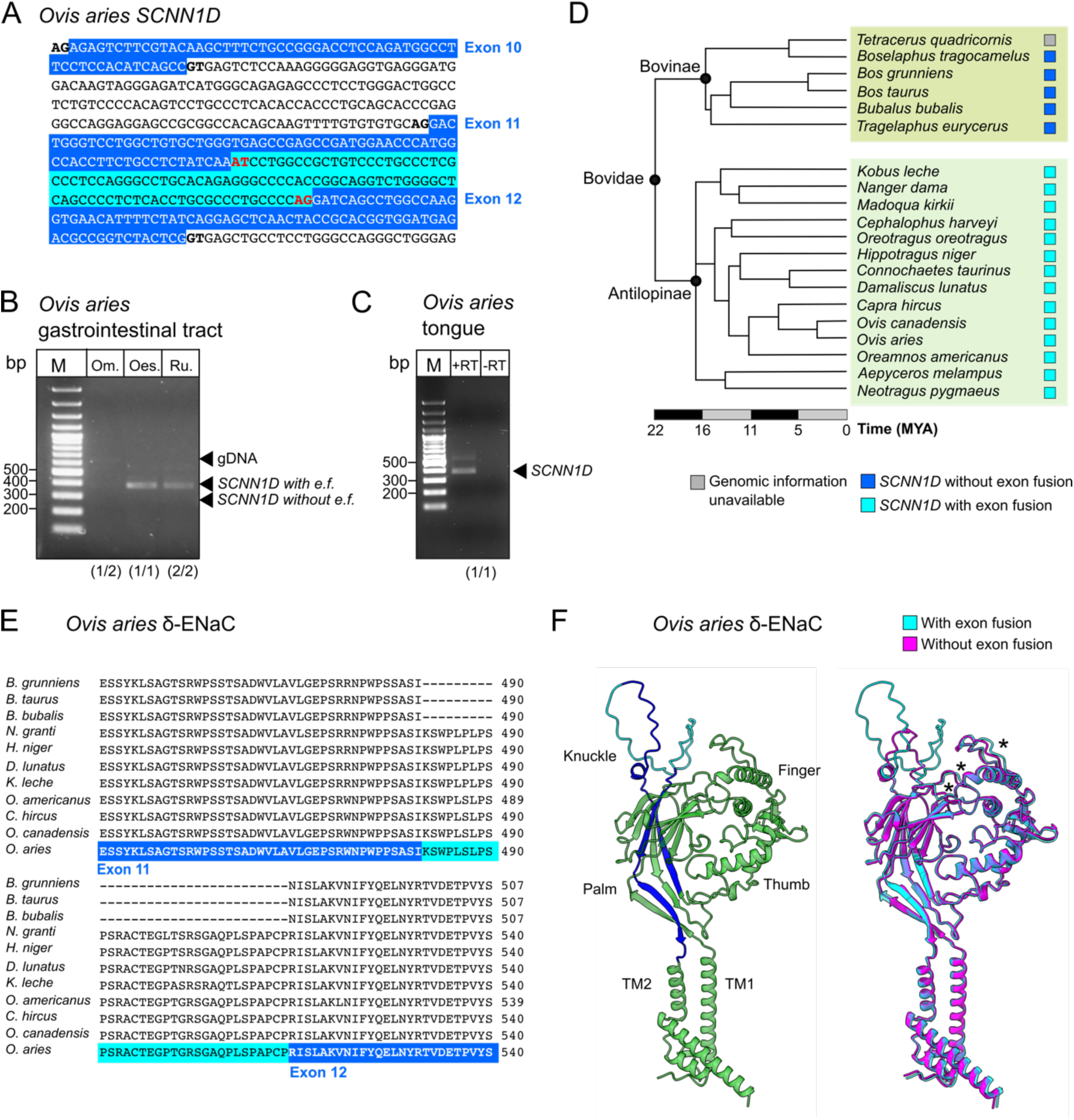
Altered *SCNN1D* genes in Antilopinae. A) Mutation of the splice donor following exon 11 to ‘AT’ in *Ovis aries* (representative of the Antilopinae) causes fusion of exons 11 and 12 to a ‘super exon’. B) Reverse transcriptase (RT)-PCR confirmed the exon fusion in cDNA derived from mRNA isolated from O. *aries* gastrointestinal tissues. Om. = Omasum; Oes. = Oesophagus; Ru. = Rumen. Arrows highlight expected amplicon sizes based on absence or presence of the exon fusion (e.f.) or contamination with genomic DNA (gDNA). bp = base pairs. M = DNA size marker. C) Reverse transcriptase (RT)-PCR confirmed expression of *SCNN1D* in sheep tongue tissue. bp = base pairs. M = DNA size marker. D) Presence of the *SCNN1D* exon fusion in Bovidae. *SCNN1D* with exon fusion (turquoise squares) is exclusively found in representative species of the Antilopinae, but not in the Bovinae. MYA = million years ago. E) Partial δ-ENaC amino acid sequence alignment of representative species of the Bovidae. Sequences highlighted in blue indicate amino acids encoded by exons 11 and 12, sequences highlighted in turquoise indicate additional amino acids due to incorporation of the former intron. F) *O. aries* δ-ENaC structures predicted by ColabFold suggest location of the additional amino acids in the ‘knuckle’ domain. For clarity, predictions of the intracellular N-/C-termini are not displayed. Left: The predicted structures follow the canonical ENaC subunit architecture. Dark blue parts are derived from former exons 11 and 12, turquoise parts highlight additional ‘intron-derived’ amino acids. Right: Alignment of the *O. aries* δ-ENaC structure containing the exon fusion (turquoise) to a hypothetical *O. aries* δ-ENaC lacking the exon fusion (magenta). The exon fusion does not impair the general channel architectures but might alter the position of loops (highlighted by asterisks) between the ‘knuckle’ and ‘finger’ domains.

The fusion of exons 11/12 and inclusion of the former intron, incorporates additional amino acids into the δ-ENaC protein. Amino acid alignment of select Bovidae indicates that the incorporated 33 amino acids are highly conserved (**Figure 3 E**). ColabFold predictions of the sheep (*O. aries*) δ-ENaC (**Source Data 2**) yielded a protein structure that is consistent with the Cryo-EM derived ‘clenched-hand holding a ball of β-sheets’ structures of human ENaC subunits (Noreng et al. 2018) (**Figure 3 F**). The additional 33 amino acids are placed in a flexible loop at the top surface of the protein, termed the ‘knuckle’ domain. Removal of the additional amino acids (**Source Data 2**) does not cause major structural rearrangements in the core protein, but subtle changes are predicted in the ‘finger’ domain and the region connecting the ‘finger’ and ‘knuckle’ domains (**Figure 3 F**). These regions contain important ENaC regulatory motifs, such as the ‘gating relief of inhibition by proteolysis’ (GRIP) domains and sodium binding sites (Noreng et al. 2018; Noreng et al. 2020). The incorporation of the additional amino acids in the ‘knuckle’ domain might have therefore altered regulatory mechanisms in δ-ENaC of the Antilopinae, but this remains to be confirmed in electrophysiological experiments. The previously observed *SCNN1D* exon 11/12 fusion in the rodent infraorder Hystricognathi (Gettings et al. 2021) affects the same extracellular ‘knuckle’ region of the ion channel. However, the incorporated intron in the Hystricognathi is considerably smaller than that of the Antilopinae, illustrating the structural flexibility of the extracellular ‘knuckle’ region of the ion channel.

Overall, our bioinformatic analyses clearly demonstrate pseudogenisation of *SCNN1D* in cetaceans, while the gene is maintained, including clade-specific alterations, in terrestrial Artiodactyla. What drove the loss of δ-ENaC in whales and dolphins, and what is the reason for its maintenance in their terrestrial sister groups? With respect to sodium homeostasis, two major physiological changes appeared with the transition from a terrestrial to a marine environment: a change from herbivory/omnivory to carnivory (Thewissen et al. 2007) and the independence from freshwater sources (Thewissen et al. 1996). Carnivores generally ingest food with similar biochemical compositions to their own bodies, thereby making specific chemical senses less necessary, while herbivores ingest a larger variety of potentially harmful plant compounds (Demi et al. 2021). Consistently, pseudogenisation of taste receptor genes has been described in several cetacean species but not in terrestrial Artiodactyla (Feng et al. 2014; Zhu et al. 2014). While extinct cetacean ancestors such as *Ambulocetus* likely relied on freshwater (Thewissen et al. 1996), modern marine cetaceans do not rely on freshwater sources, and they are well adapted to maintain electrolyte homeostasis in a hypersaline marine environment (Ortiz 2001). Hence, a necessity to discriminate seawater from freshwater (based on sodium content), or a ‘stress response’ warning of sodium concentrations greater than plasma concentrations appears less necessary in cetaceans, while it might be vital in terrestrial Artiodactyla. Furthermore, herbivorous diet generally demands more stringed sodium conserving mechanisms: sodium concentrations in plant tissue are lower than in animal tissue and herbivores often look for sites for sodium supplementation in their environments (mineral licks) (Kaspari 2020). Digestion of plant material also requires significant amounts of saliva (with sodium as the main cation) ensuring an optimal environment for microbial digestion, and sodium reabsorption in the kidneys and gastrointestinal tract is pivotal to avoid a sodium deficit in ruminants (Leonhard-Marek et al. 2010). Renal and gastrointestinal adaptations as well as adaptations with respect to chemical sodium senses might therefore explain the pseudogenisation of *SCNN1D* in cetaceans and its preservation in terrestrial Artiodactyla.

Consistent with our bioinformatics data, previous studies demonstrated the presence of *SCNN1A*, *SCNN1B* and *SCNN1G* genes encoding αβγ-ENaC in some cetacean species, while the genes encoding sweet and bitter receptors pseudogenised (Feng et al. 2014; Zhu et al. 2014). These studies led to the conclusion that ‘salty’ is the only taste modality cetaceans can sense. However, αβγ-ENaC is not only key for maintaining vertebrate sodium homeostasis, but also enhances the electrochemical gradients for potassium excretion via ‘Renal Outer Medullary Potassium’ (ROMK) channels in the principal cells of the distal nephrons and it controls the volume of liquid lining the lung epithelia (Wichmann and Althaus 2020). We isolated mRNA from kidney and lung tissues of three Bottlenose dolphins (*Tursiops truncatus*) and confirmed expression of αβγ-ENaC in these tissues by RT-PCR (**Figure 4, Source Data 3**). Although αβγ-ENaC has been shown to mediate appetitive sodium taste to low sodium concentrations in mice (Chandrashekar et al. 2010; Nomura et al. 2020), the presence of functional *SCNN1A*, *SCNN1B* and *SCNN1G* genes in cetaceans does not necessarily associate with the ability to taste sodium, given the importance of this ion channel in dolphin kidneys and lungs.

**Figure 4.**
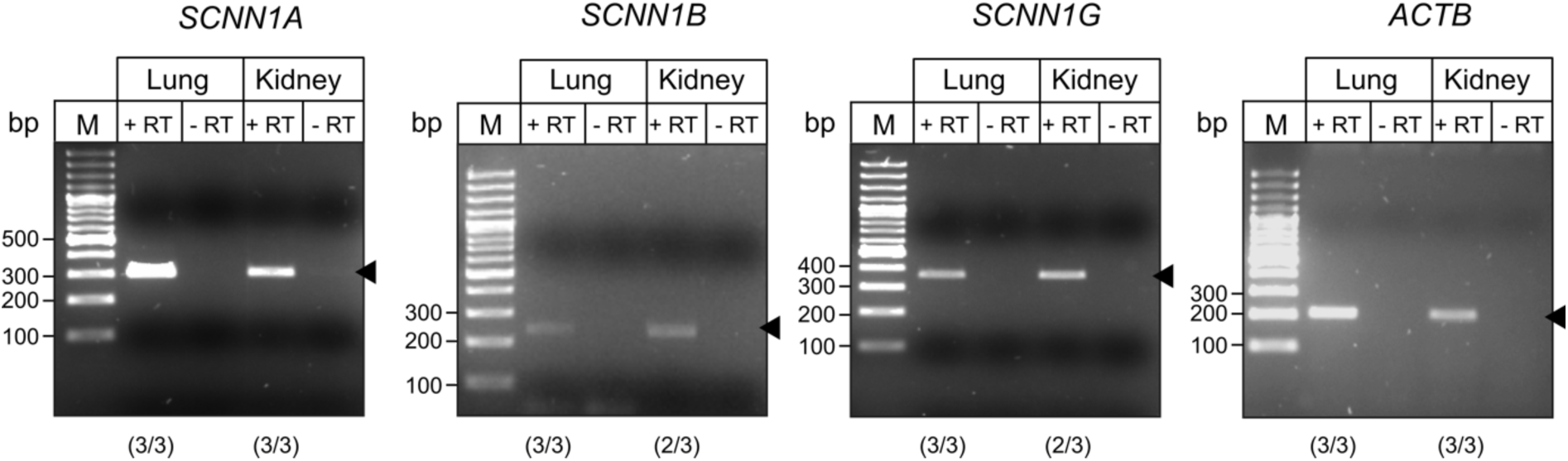
Expression of ENaC-coding genes in Bottlenose dolphin (*Tursiops truncatus*) lung and kidney tissues. RT-PCRs were performed with cDNA synthesised from tissues derived from three animals. Arrow heads point towards the expected amplicon sizes. Numbers in parentheses indicate positive amplicons out of the total number of animals. PCR results from all animals are provided as **Source Data 3**. β-actin (*ACTB*) was used as controls. RT = Reverse Transcriptase; M = DNA size marker; bp = base pairs.

Whether or not cetaceans have a functional gustatory system and can taste sodium remains controversial (Bouchard et al. 2017; Werth and Crompton 2023). Behavioural studies demonstrated sodium tasting abilities in Bottlenose dolphins (Friedl et al. 1990; Kuznetzov 1990), however, these studies were performed on very few animals, were not blinded and have been criticized due to their Go/No-Go experimental design (Kremers et al. 2016). We, therefore, performed a series of blinded behavioural experiments on Bottlenose dolphins (*T. truncatus*) and Beluga whales (*Delphinapterus leucas*) under human care to investigate their sodium tasting abilities.

We modified an experimental approach that has been employed to study spontaneous responses and preferences to chemical stimuli in cetaceans (Kremers et al. 2016). Cetaceans under human care are routinely offered gelatine blocks to provide hydration and nutritional enrichment. We provided gelatine blocks prepared with or without 500 mM NaCl (similar to seawater concentrations, (Bœuf and Payan 2001)) to six Bottlenose dolphins (three males and three females) and two Beluga whales (one male and one female) and determined the latency of begging for further gelatine blocks as a measure of discrimination between gelatine with or without added NaCl (**Figure 5 A, B**). We considered the gelatine without added NaCl the atypical exposure for marine cetaceans. Experiments were blinded and the order of gelatine block application was randomised. In total, there were 230 latency observations for Bottlenose dolphins and 96 latency observations for Beluga whales. Latency times varied between individual animals but were highly similar between gelatine with or without NaCl (**Figure 5 A, B, Supplemental Data 4, Source Data 4**). Latency data were logarithmically transformed and modelled with a mixed effects structure in a Bayesian framework, which demonstrated that the latency times did not statistically vary with the presence or absence of NaCl in the gelatine blocks in either Bottlenose dolphins or Beluga whales (**Supplemental Data 4, Source Data 4**). However, in some experiments, the cetaceans submerged their heads after providing the gelatine blocks and thereby took up seawater in their oral cavity before swallowing the gelatine. The Bayesian mixed effects model revealed a significant increase in latency time when seawater is present in the oral cavity of approximately 30% for both species (**Supplemental Data 4, Source Data 4**). It might take longer for the animals to process the blocks while swallowing the seawater, however, it highlights the possibility that seawater might mask the absence or presence of NaCl in the gelatine blocks in this experimental setting. We, therefore, took advantage of a specific sampling and interaction behaviour that was displayed by three Bottlenose dolphins in response to installation of additional water jet sources as environmental enrichment devices, (**Figure 5 C**) and analysed the spontaneous interaction of the animals with the provided fresh- and seawater sources in 15 trials lasting 15 min each. There were no significant differences in the number of interactions (**Figure 5 D**) as well as the interaction times (**Figure 5 E**), indicating that there was no observable preference for either water source. In summary, these experiments did not reveal measurable behavioural differences to stimuli with or without sodium in seawater-equivalent concentrations in cetaceans. However, we would like to emphasize that a limitation of these experiments is the underlying assumption that the animals would display measurable behavioural differences (shift in latency time or interaction with the water sources) if they were able to distinguish between stimuli with or without NaCl. While previous studies with Bottlenose dolphins detected changes in latency of begging behaviour (Kremers et al. 2016) and sampling/interaction behaviour (Bruck et al. 2022) in response to chemical stimuli, the absence of equivalent behavioural changes in response to NaCl cannot be excluded.

**Figure 5.**
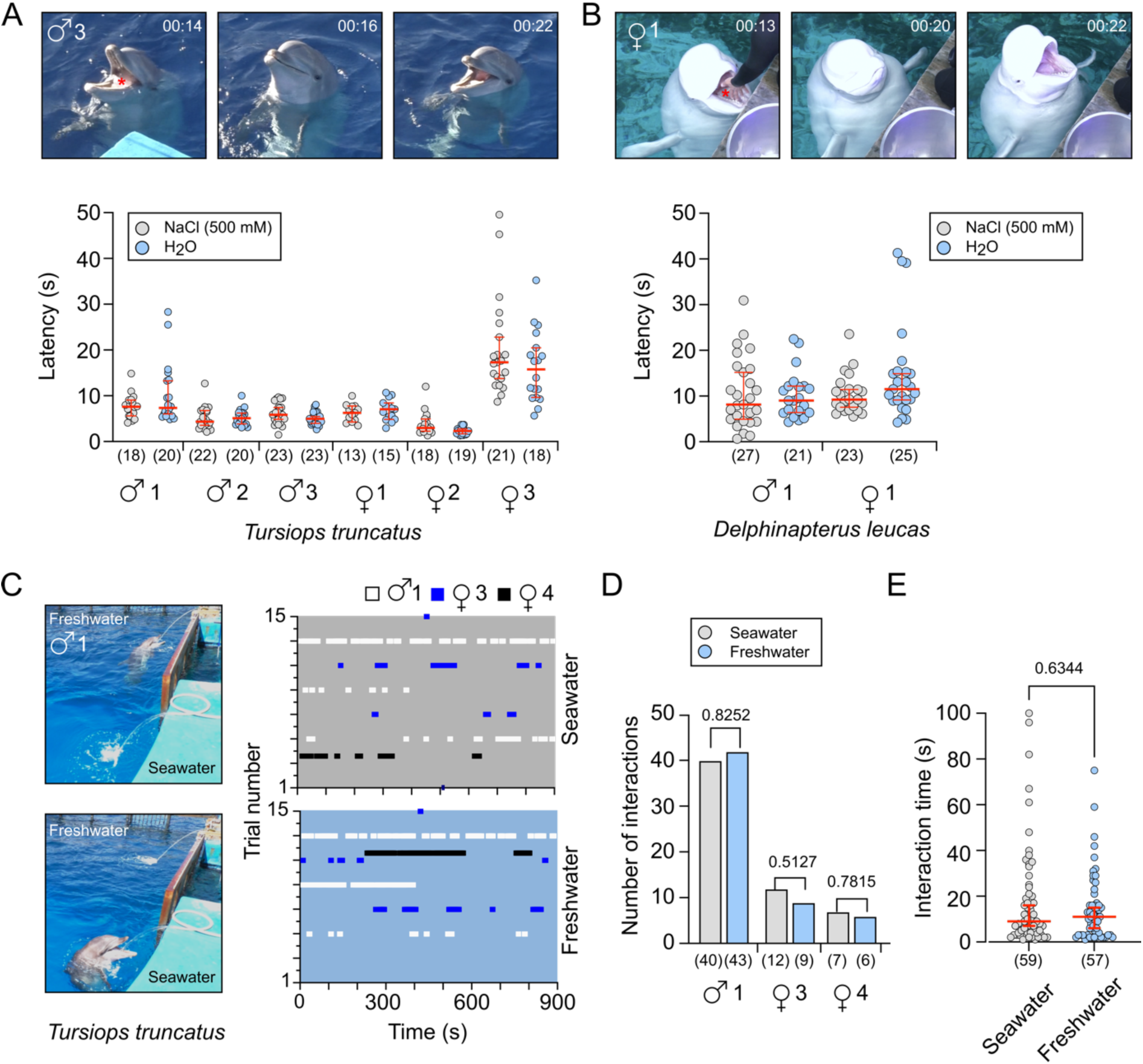
Behavioural assessment of salt tasting abilities in cetaceans. A, B) The latency of begging behaviour (s) in response to feeding of 3 gelatine blocks with or without 500 mM NaCl was determined for 6 Bottlenose dolphins (A) and 2 Beluga whales (B). The images are video stills from the recorded sessions, indicating the time of gelatine block feeding until the display of begging behaviour. Gelatine blocks are marked with a red asterisk. Numbers in parentheses indicate “n”. Red lines indicate medians ± 95 % confidence intervals. For statistical analyses, the dataset was modelled with a mixed effects structure in a Bayesian framework. Statistical data are provided in **Supplemental data 4** and **Source Data 4**. C) A freshwater and a seawater source were installed and the interaction of 3 Bottlenose dolphins with the water sources were measured. There were 15 trials lasting 15 min each and the interaction times of each animal with the respective water source is displayed. D) There were no differences in the number of interactions (chi-square test) between the two water sources. E) There was no significant difference in the overall interaction time between freshwater and seawater sources. Numbers in parentheses indicate “n”. Red lines indicate medians ± 95 % confidence intervals (Mann-Whitney U-Test).

A lack of sodium taste in cetaceans would be consistent with the pseudogenisation of *SCNN1D* and functional δ-ENaC in cetaceans. The emergence of this ENaC subunit with the transition from aquatic to terrestrial environments in the Devonian and its pseudogenisation with the transition from terrestrial to marine environments in the Eocene generally suggests a role for δ-ENaC in sodium homeostasis in low-salinity and terrestrial habitats. A counterargument to this hypothesis is illustrated by extant river dolphin species which live in low-salinity environments and also lack a functional *SCNN1D* gene (**Figure 1 and 2**). However, these species evolved from marine ancestors (de Muizon et al. 2018), indicating that they did not regain δ-ENaC after migration from marine to freshwater environments. Furthermore, ENaC-independent physiological mechanisms for the detection of salinity might have evolved in cetaceans. For example, mechanical effects on odontoblastic processes in the Narwhal tusk have been hypothesized to mediate sensory responses to salinity gradients in these cetaceans (Nweeia et al. 2014).

Secondary transitions to marine environments have occurred multiple times and independently in mammals. We therefore investigated whether *SCNN1D* pseudogenised in other marine lineages including Sirenia (manatees and dugongs), Pinnipedia (seals) and the sea otter. Consistent with the analysed Cetartiodactyla, *SCNN1A*, *SCNN1B* and *SCNN1G* remain intact in all investigated species (**Figure 6, Supplemental Data 5, Source Data 5**). *SCNN1D* is a pseudogene in the aquatic Sirenia, but also in all other investigated Afrotheria (**Figure 6**). The gene is intact in the two analysed Xenarthra (**Figure 6**). Interestingly, the ancestors of the Paenungulata, a clade comprising the Sirenia, Proboscidea (elephants) and Hyracoidea (hyraxes), were likely amphibious (Springer 2022; Liu et al. 2024) and fossils of early sirenians and proboscideans are found in marine sediments (Seiffert 2007), suggesting that *SCNN1D* might have been lost before the transition back to a terrestrial environment. The gene also pseudogenised in the investigated species belonging to the afrotherian clade Afroinsectiphilia (aardvark, elephant shrews, tenrecs) but their evolutionary origin relative to the Paenungulata is less clear (Liu et al. 2024). The analysed Afrotheria share a dysfunctional splice acceptor flanking exon 3 and splice donor flanking exon 12 (**Supplemental Data 5**). However, the splice site alteration is not identical between species and it is therefore difficult to conclude whether *SCNN1D* was lost by a single gene inactivating event at the base of Afrotherian evolution or the gene pseudogenised independently in different Afrotherian lineages. Within the Mustelidae, a clade of the Carnivora including otters, badgers, martens and weasels, *SCNN1D* pseudogenised only in the sea otter (*Enhydra lutris kenyoni*), consistent with the loss of *SCNN1D* in species adopting a marine lifestyle. In the Pinnipedia, we detected pseudogenisation of *SCNN1D* in three species belonging to the Phocidae, two elephant seal species (*Mirounga leonina* and *Mirounga angustirostris*) as well as the Weddel seal (*Leptonychotes weddellii*), while it remained intact in the Otariidae (eared seals). There are no shared inactivating mutations in the elephant seals and Weddel seal, suggesting that *SCNN1D* pseudogenised independently. In the Ursidae (bears) *SCNN1D* remains intact in all investigated species, including the polar bear (*Ursus maritimus*).

**Figure 6.**
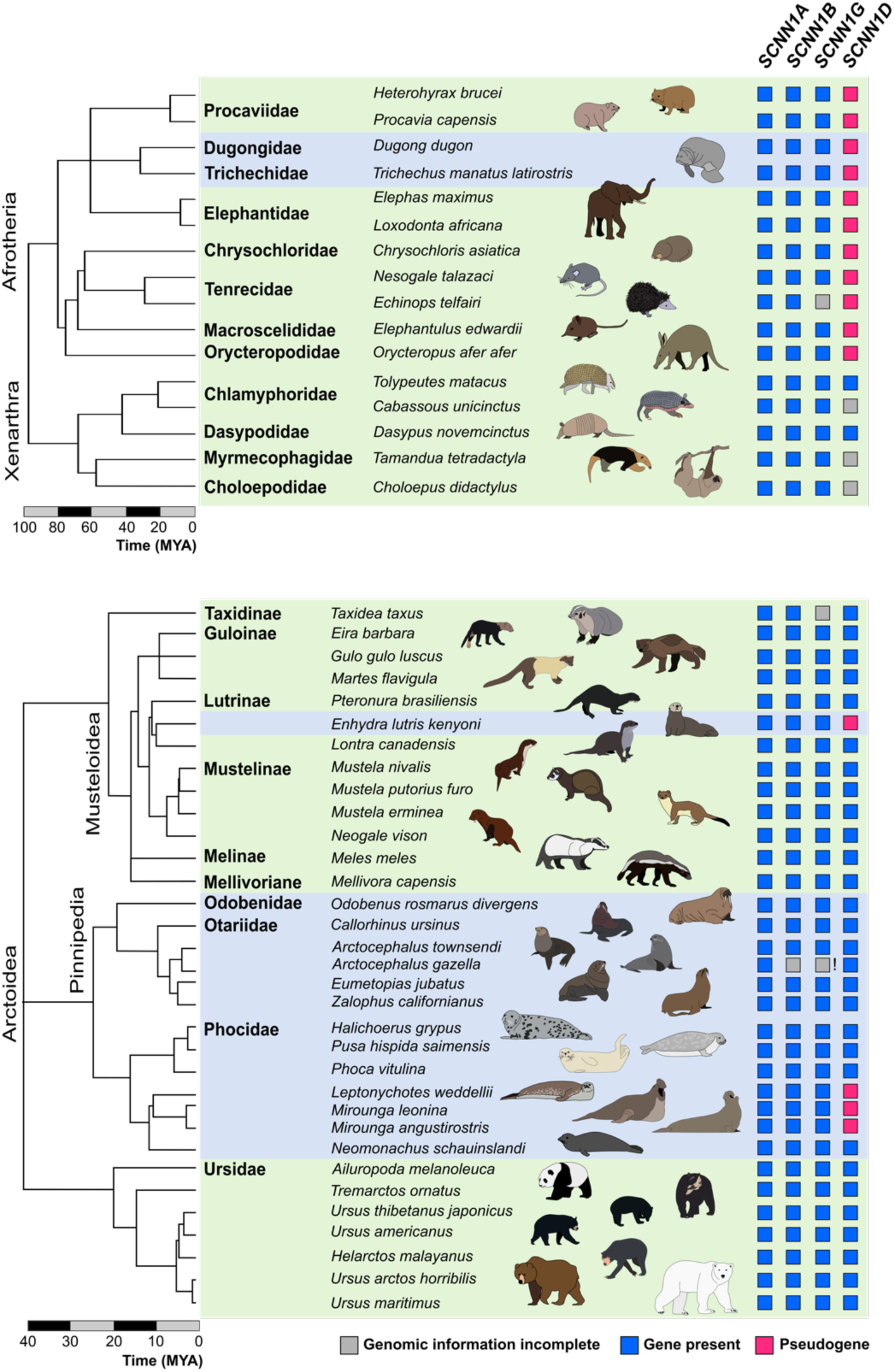
Presence of ENaC-coding *SCNN1* genes in Afrotheria and Carnivora. Presence or pseudogenisation of *SCNN1* genes was investigated in the highlighted species using the National Center for Biotechnology Information (NCBI) database. Blue squares indicate intact genes yielding continuous open reading frames (ORFs), whereas magenta squares indicate pseudogenes containing multiple insertions/deletions and splice donor/acceptor site mutations causing preliminary STOP codons. Gray squares highlight incomplete genomic information. ! = Likely sequencing error in exon 6. The phylogenetic trees and time indices were obtained from Timetree (http://timetree.org).

In order to analyse selection pressure on the *SCNN1* genes, we used codon alignments of the four genes of all investigated species, in combination with rooted trees of the four *SCNN1* genes. We applied tests for site-specific and branch-specific effects (**Source Data 6**). The result of the BUSTED (branch-site unrestricted statistical test for episodic diversification) test for episodic diversification was significant for the genes *SCNN1A* and *SCNN1D* (**Table 1**). FEL (fixed effects likelihood) and SLAC (single-likelihood ancestor counting) tests flagged a small number of sites (less than ten) to be under positive selection in all the four genes under study. The number of sites under negative (= stabilising) selection comprised about 50% (FEL) or 40% (SLAC) of the codons in genes *SCNN1A*, *SCNN1B* and *SCNN1G*, while it was substantially less in *SCNN1D* (36% in FEL and 23% in SLAC test). This suggests that *SCNN1D* is evolving under relatively relaxed conditions, at least for a few branches. Testing with SLAC explicitly for single branches it became apparent that pinnipeds (just 2 sites under negative selection) are strikingly different from terrestrial branches of Artiodactyla (54 sites under negative selection) or Carnivora (49 sites under negative selection) (**Source Data 6**). Thus, the loss of this gene in some pinnipeds makes sense as it seems to evolve under much more relaxed conditions in this branch.

**Table 1.**
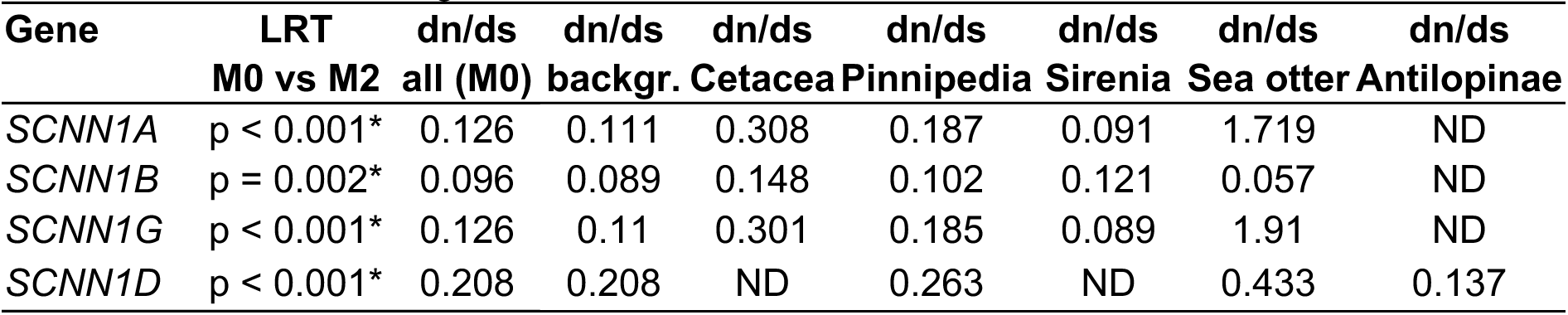
Statistical tests on site-wise selection (busted: p-value for any sites under positive selection, FEL / SLAC: number of sites under pos./neg. selection)

Likelihood-ratio tests for all four genes show, that dn/ds ratio is significantly better modelled when a non-uniform ratio is assumed comparing different branches (**Table 2**). We especially checked the aquatic branches independently from the terrestrial ones. Although dn/ds ratios are generally below 1 for the genes in total (meaning that in general negative selection is predominant over all sites), pinnipeds and cetaceans have slightly higher dn/ds ratios in comparison to the background (= the terrestrial taxa). The sea otter has values greater than 1 in genes *SCNN1A* and *SCNN1G* suggesting strong positive selection on these genes. Although below 1, the pseudogene *SCNN1D* of the sea otter (which could be reconstructed due to minimal alterations and absence of deletions) has the highest value in comparison to the other branches. As described above, the Antilopinae underwent an extra insertion in *SCNN1D* (fusion of exons 11/12). Interestingly, here *SCNN1D* shows a smaller dn/ds ratio compared to the background, implying a stronger negative selection.

**Table 2.**
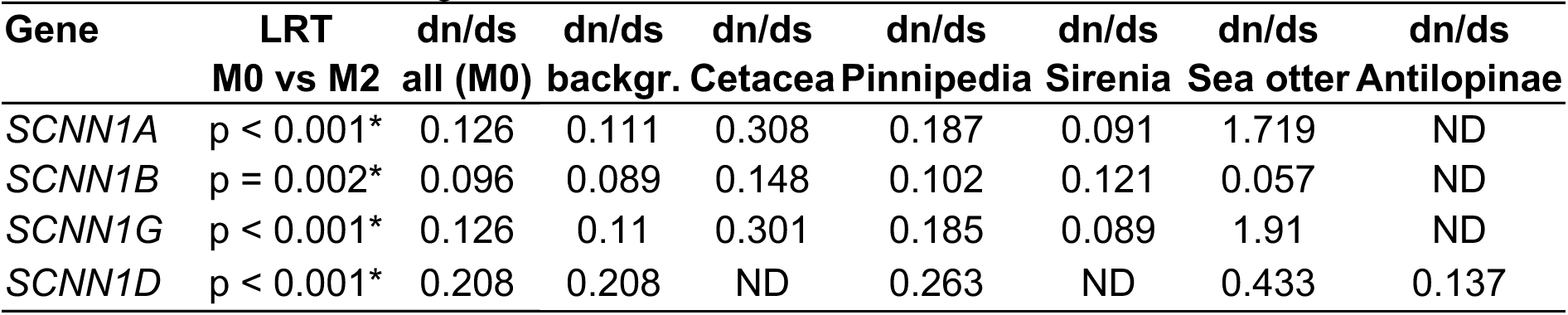
Likelihood ratio test (LRT) of uniform (M0) versus branch-specific dn/ds ratios (M2) and dn/ds values for background and selected branches.

In summary, the analysis of *SCNN1D* in marine mammals reveals the following patterns: (1) *SCNN1D* tends to pseudogenise in lineages that transitioned to marine environments, including cetaceans, sirenians, pinnipeds and the sea otter. (2) Pseudogenisation or preservation of *SCNN1D* is not consistent with herbivorous or carnivorous diet, thereby suggesting that dietary/gastrointestinal adaptations are not sufficient to explain loss of this gene. (3) *SCNN1D* is also a pseudogene in some terrestrial mammals, including the Afroinsectiphilia (**Figure 6**) as well as five rodent lineages (Gettings et al. 2021). It has been hypothesised that δβγ-ENaC represents a ‘stress protein’ monitoring high sodium concentrations. Selection pressure for such a sodium sensing mechanism may not only decrease upon transition into a high-salinity marine environment, it may, for example, also be reduced when renal sodium handling abilities of terrestrial species renders high salt intake less problematic. Such scenarios might explain the loss of *SCNN1D* in terrestrial species.

Overall, αβγ-ENaC is a key ‘maintenance protein’ controlling the overall body sodium homeostasis in mammals. Furthermore, it is important for renal potassium handling as well as control of the volume and composition of lung lining liquids (Wichmann and Althaus 2020). These physiological functions are required in terrestrial and marine mammals and explain the presence of αβγ-ENaC transcripts in kidney and lung tissues from marine species, e.g. Bottlenose dolphins (**Figure 4**). By contrast, functional electrophysiological studies on heterologously expressed δβγ-ENaCs suggest that this isoform is primed for operation under high extracellular sodium concentrations due to reduced regulatory constraints such as SSI. In humans, δ-ENaC is not expressed in the kidney, but transcripts are found in taste buds (Huque et al. 2009; Xu et al. 2017; Bigiani 2020) as well as immune cells which may have an ENaC-mediated mechanism sensing high extracellular sodium concentrations (Pitzer et al. 2022). These lines of evidence point towards a function as a ‘stress protein’ monitoring high sodium concentrations in tetrapod vertebrates. This function might have become obsolete in animals which migrated to high-salinity marine environments.

## Materials and Methods

### Bioinformatic analyses of Cetartiodactyla genomes

*SCNN1* gene sequences were analysed as previously described (Gettings et al. 2021; Maxeiner et al. 2023). All employed databases and resources are listed in **Table 3**. The exon sequences of the *SCNN1A*, *SCNN1B*, *SCNN1G* and *SCNN1D* genes of the Alpaca (*Vicugna pacos*) were used as a template to blast the whole genome contig database of the National Center for Biotechnology Information (NCBI BLAST) with default settings for blast and megablast searches. Exons 2-13 of all *SCNN1* genes were aligned to the Alpaca template sequences using Multalin (Corpet 1988). Details on exon and intron sizes, splice donor/acceptor sites as well as insertions/deletions with respect to the Alpaca templates are provided in **Supplemental Data 1**. Coding sequences were translated using the Expasy Translate Tool to retrieve open reading frames. Amino acid sequences were aligned using Clustal Omega. Structure predictions (Supplemental Data 4) of δ-ENaC subunits from sheep (*O. aries*) displayed in **Figure 3** were performed with ColabFold v1.5.5 (Mirdita et al. 2022) based on amino acid sequences provided in **Source Data 1**. Molecular graphics and analyses were performed with UCSF ChimeraX (Meng et al. 2023), developed by the Resource for Biocomputing, Visualization, and Informatics at the University of California, San Francisco, with support from National Institutes of Health R01-GM129325 and the Office of Cyber Infrastructure and Computational Biology, National Institute of Allergy and Infectious Diseases.

**Table 3.** Employed online tools and resources in this study.

| Resource | Weblink | Access period<br>(month/year) |
| --- | --- | --- |
| NCBI BLAST | https://blast.ncbi.nlm.nih.gov/Blast.cgi | 01/2022-08/2024 |
| Multalin | http://multalin.toulouse.inra.fr/multalin/ | 01/2022-08/2024 |
| ExPASy Translate Tool | https://web.expasy.org/translate/ | 01/2022-08/2024 |
| Clustal Omega v1.2.4 | https://www.ebi.ac.uk/Tools/msa/clustalo/ | 01/2022-04/2024 |
| Timetree | http://timetree.org | 11/2023-08/2024 |
| ColabFold v1.5.5 | https://colab.research.google.com/github/sokrypton/ColabFold/blob/main/Colabfold.ipynb | 01/2024-04/2024 |

### Phylogenetic analyses of SCNN1 genes

Time-calibrated phylogenetic trees were generated using Timetree (accessed November 2023) (Kumar et al. 2022). For the analysis of selection pressure, we used codon alignments of the four genes in combination with rooted trees of the four *SCNN1* genes (**Source Data 6**). We applied tests for site-specific and branch-specific effects. BUSTED, FEL and SLAC were conducted with the HyPhy package (Kosakovsky Pond et al. 2005). With BUSTED (branch-site unrestricted statistical test for episodic diversification) we tested if there is at least one site under positive selection in the whole phylogeny (Murrell et al. 2015). FEL (fixed effects likelihood; (Kosakovsky Pond and Frost 2005)) tests for deviation from an even distribution of dn/ds ratio across the phylogeny. SLAC (single-likelihood ancestor counting; (Kosakovsky Pond and Frost 2005)) analyses dn/ds ratio on a per site basis, suggesting sites under positive or negative selection pressure. General dn/ds ratio for whole genes was analysed with codeml from the PAML package(Yang 2007). Hypothesis testing for different settings (uniform dn/ds versus branch-wise variation) was then done using the likelihood ratio test (LRT) as implemented in EasyCodeML (Gao et al. 2019).

### Tissue mRNA isolation and Reverse Transcriptase (RT) - PCR

Sheep (*O. aries*) gastrointestinal tissue samples were kindly provided by Dr. Franziska Liebe, Institute for Veterinary Anatomy, Free University Berlin, Germany. Tissue isolation was under governance by the Berlin Veterinary Health Inspectorate (Landesamt für Gesundheit und Soziales Berlin) and approved under the animal experimentation number G0098/21. Sheep tongue tissue samples were kindly provided by Prof. Franziska Dengler, Department of Biochemical Sciences, Institute of Physiology, University of Veterinary Medicine, Vienna, Austria. Tissue isolation was under governance by the Federal Ministry Republic of Austria, Education, Science and Research, and approved under the animal experimentation number BMBWF-68.205/0100-V/3b/2018. Bottlenose dolphin (*T. truncatus*) tissue samples were kindly provided by the tissue bank of Fundación Oceanogràfic, Valencia, Spain and Mundomar Benidorm, Alicante, Spain. Tissue samples were stored in DNA/RNA stabilizer solution at -80°C (New England Biolabs, Frankfurt, Germany) according to manufacturer’s instructions.

Bottlenose dolphin tissues were homogenised with the Precellys Evolution Touch homogeniser (Bertin Technologies), using the parameters for hard tissue samples (3 x 20 s at 6800 rpm, 30 s pause). Homogenisation was repeated 4 times, with a 5 min incubation on ice after the 2nd cycle. RNA was isolated with the Monarch® Total RNA Miniprep Kit (New England Biolabs) according to manufacturer’s protocol. Prior isolation, Proteinase K was added to the homogenates and the mixtures were incubated for 5 min at 55 °C. RNA was eluted in nuclease free water and stored until further use at -80 °C. To further remove residual genomic DNA, an in-tube treatment with DNase I was performed using Monarch® DNase I and RNA was purified using the Monarch® RNA Cleanup Kit according to manufacturer’s instructions (New England Biolabs). Isolated RNA (0.5 - 1 μg) was reverse transcribed into cDNA using the LunaScript RT Master Mix Kit and d(T)23VN primers according to manufacturer’s instructions (New England Biolabs). Negative controls lacking the reverse transcriptase (RT) enzyme were performed using the No-RT ControlMix provided by the same manufacturer. 1 μl of the cDNA reactions were used as templates for PCR reactions using primers listed in **Table 4**. PCRs of cDNA derived from Bottlenose dolphin tissues were performed with the One Taq 2X Master Mix with Standard Buffer (New England Biolabs) according to manufacturer’s instructions. PCRs started with an initial denaturation at 94 °C for 30 s, followed by 34 cycles of denaturation for 30 s at 94 °C, annealing at 57 °C for 30 s and extension for 30 s at 68 °C. After a final extension at 68 °C for 5 min, PCR products were mixed with TriTrack loading dye (ThermoFisher, Darmstadt, Germany) and loaded on 2 % agarose gels containing 2 µg/100 ml Midori Green (Nippon Genetics, Düren, Germany). PCR amplicons were visualised using an ChemiDoc XRS+ Molecular Imager (Biorad, Feldkirchen, Germany).

Sheep tissues were homogenised with the Precellys Evolution Touch homogeniser, using the parameters for soft tissue samples (15 s at 5600 rpm, 30 s pause, 15 s at 5600 rpm). Homogenisation was repeated 4 times, with a brief incubation on ice after each cycle. Proteinase K was added to the homogenates and the mixtures were incubated for 20 min at 55 °C. Afterwards, RNA extraction was performed exactly as described above. PCR was performed using Q5® High-Fidelity DNA Polymerase and reactions started with an initial denaturation at 98 °C for 30 s, followed by 30 (gastrointestinal tissues) or 34 (tongue tissue) cycles of denaturation for 10 s at 98 °C, annealing at 69 °C for 30 s and extension at 72 °C for 30 s. After a final extension at 72 °C for 2 min, PCR products were mixed with TriTrack loading loaded on 2 % agarose gels containing 2 µg /100 ml Midori Green and PCR amplicons were visualised using a ChemiDoc XRS+ Molecular Imager (Biorad).

**Table 4.**
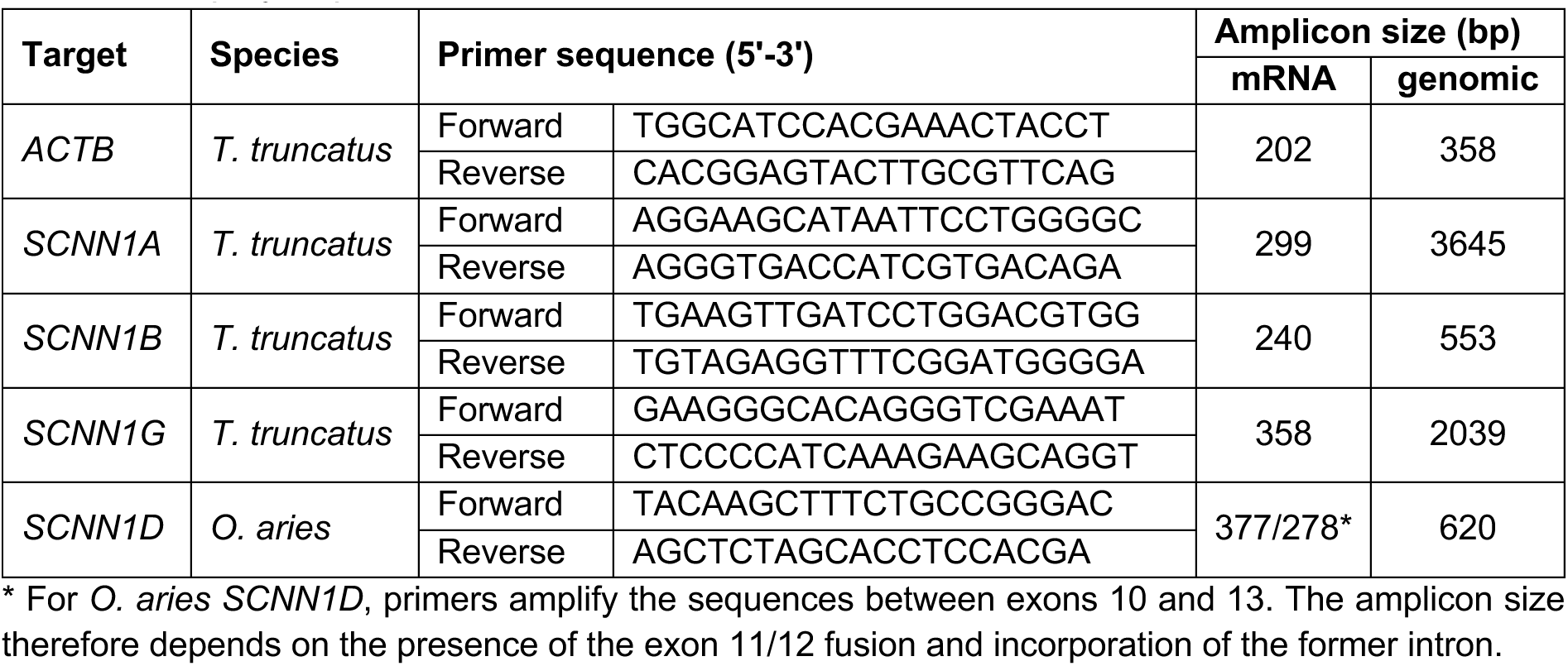
Employed primers for RT-PCR reactions.

All PCR amplicons of appropriate sizes were isolated with the Monarch DNA Extraction Kit (New England Biolabs) and sequenced (Eurofins Genomics, Ebersberg, Germany).

### Behavioural assessment of salt tasting abilities in cetaceans

Behavioural experiments aiming to investigate salt tasting abilities in cetaceans under human care were performed between March and May 2023 at the Oceanogràfic, Ciudad de las Artes y las Ciencias, Valencia, Spain. Experiments were approved by the Oceanogràfic Animal Care & Welfare Committee under the project reference OCE-11-23. Experiments were performed with 6 adult Bottlenose dolphins (*Tursiops truncatus*), 3 males and 3 females, and 2 Beluga whales (*Delphinapterus leucas*), 1 male and 1 female. Two experimental approaches were employed: (i) determination of the latency of begging behaviour in response to gelatine blocks without or with 500 mM NaCl; and (ii) analysis of interactions with provided freshwater and seawater sources.

*(i) Determination of latency of begging behaviour:* Gelatine blocks were prepared by dissolving 100 g gelatine (Promolac, Cornellá del Terri, Spain; Lot. No. 22V2382) with or without 87.7 g of NaCl (Scharlab, Barcelona, Spain) in 1 L of hot deionized water. Afterwards, 2 L of deionized water were added, and the mixture was allowed to solidify over night at 3 °C. This yielded either gelatine with or without 500 mM NaCl. The solidified gelatine was cut into 36 equally sized blocks of 75 g. Each experimental trial started by feeding 3 gelatine blocks at once to one animal. Three blocks were fed simultaneously to enhance the contact time with the oral mucosa. Afterwards the trainers assumed a neutral position and waited for the animals to display begging behaviour, after which another 3 blocks of gelatine were fed. This procedure was repeated so that each trial consisted of 4 stimuli (2 x 3 gelatine blocks with NaCl; 2 x 3 gelatine blocks without NaCl). The order of stimuli was randomised, and the trainers were blinded. For Bottlenose dolphins, begging behaviour was defined as a clear behavioural response towards the trainer, while the animals had both eyes above the water and the empty mouth was opened (**Figure 3 A, B**). The same criteria were applied to beluga whales, with the exception that begging behaviour was sometimes displayed with the eyes under water due to the larger size of the animals. The experimental trials were recorded on video and latency times were determined from the time of contact of the gelatine blocks with the oral cavity and the earliest moment begging behaviour was initiated.

To analyse the Bottlenose dolphin data, latency values were logarithmic transformed, and the dataset was modelled with a mixed effects structure in a Bayesian framework, with the following expectation:

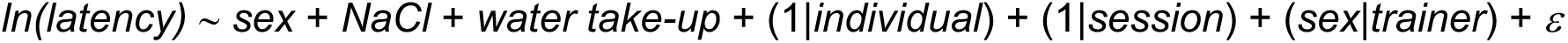

where *sex* is a separate intercept value for females and males; *NaCl* is the binary factor indicating whether the gelatine block contains *NaCl* or not (absence of NaCl is taken as the reference) with the expectation that it is not statistically distinguishable; *water take-up* is a binary factor indicating whether there is water in the mouth of the cetacean while being fed the block, as this may increase the processing time of the block and therefore increase latency (absence of water is taken as the reference); (1|*individual*) is the random effect of the individual dolphin; (1|*session*) is the random effect of the experiment session, which should account for any bias between days including potential differences in temperature; (*sex*|*trainer*) is the random effect of the trainer, conditional on the sex of the dolphin (i.e. dolphins of different sexes may respond differently to trainers due to social behaviours); and *ɛ* is the residual error that is Gaussian distributed about mean zero. The Bayesian models were run as a single chain with vague priors, for 1000000 iterations with a burn-in of 500000 and a thinning interval of 100, giving a posterior sample size of 5000. Model convergence was checked using the geweke diagnostic which tests for the equality of means between the initial 10 % of the final 50 % of samples. For the Bottlenose dolphin model, all 28 variables had converged (standard z-scores < 2). The models were run in R 4.3.1 (R Core Team 2023) using the “MCMCglmm” (v. 2.35) (Hadfield 2010).

The Beluga whale data was modelled in a similar fashion, but as there were only two animals, one of each sex, differences due to sex-based or individual differences are confounded with no replication. Therefore, only individual differences were modelled:

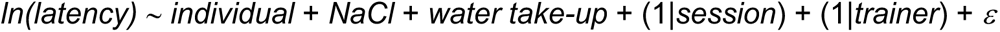

where *individual* is the intercept value separated for the male and female Beluga whale. For the Beluga whale model, all 25 variables had converged (standard z-scores < 2), with model setup parameterised the same as the bottlenose dolphin model. Data and statistical models are provided in Supplementary Data 5.

*(ii) Interaction with freshwater and seawater sources:* Three of the six Bottlenose dolphins (1 male and 2 females) voluntarily played with water that was fed into their pool via hoses. To study their interaction with seawater and freshwater stream, two water hoses were installed, one was feeding seawater from the neighbouring pool, the other one was connected to freshwater (**Figure 3 C**). In both cases, the water pressure was kept at 26 ± 2.5 L/min and temperature ranged between 22 and 25 °C depending on the day of the trial. Once the water jets were installed, the animals were left alone for 15 min and their interaction with the water streams was recorded on video. On some days, the animals did not interact with the water jets and those trials were excluded from the analysis. Overall, 15 trials were successful. The number of interactions with either freshwater or seawater source was determined for each animal and analysed with a chi-square test. In addition, the total interaction times (s) were determined and analysed with a Mann-Whitney test. Statistical analyses were performed with GraphPad Prism (v. 10) (GraphPad Software, San Diego, USA).

## Acknowledgments

We would like to thank Adrian González Quintero and the cetacean team of trainers at the dolphins and belugas at Oceanogràfic for their work and dedication with the animals and their outstanding support with cetacean experiments. We thank Dr. Julia Holtel (IFGA, Hochschule Bonn-Rhein-Sieg) for help with genetic analyses and Dr. Faysal Bibi (Leibniz Institute for Evolution and Biodiversity Science, Berlin, Germany) for expert advice on bovine evolution. We thank Dr. Franziska Liebe (Institute for Veterinary Anatomy, Free University Berlin, Germany) and Prof. Franziska Dengler (Department of Biochemical Sciences, Institute of Physiology, Pathophysiology and Biophysics, University of Veterinary Medicine, Vienna, Austria) for providing sheep tissue samples and Mundomar Benidorm (Benidorm, Alicante, Spain) for providing Bottlenose dolphin tissue samples.

## Availability of data and materials

Data are made available in the manuscript, as Supplemental Data or as Source Data deposited at the Zenodo data depository upon final publication of the manuscript.

