## Supplemental Data 2 for "The evolutionary path of the epithelial sodium channel δ-subunit in Cetartiodactyla points to a role in sodium sensing"

### ENaC subunit sequence alignments

**α-ENaC (*SCNN1A*) amino acid sequence alignment** created with Clustal Omega (v.1.2.4, accessed 24.03.2024). Key structural motifs are highlighted in the human ENaC subunit based on the Cryo-EM derived structure (Noreng et al. 2018). Transmembrane domains (TM1/TM2) are highlighted in yellow. An N-Terminal HG-motif affecting ENaC open probability is marked in magenta. Protease cleavage sites are highlighted in blue. Cysteines involved in tertiary structure formation are indicated in red. Amino acids contributing to sodium ion binding in the cation binding pocket (Noreng et al. 2020) are shown in dark blue. Residues putatively forming the selectivity filter within TM2 are shown in red font. The C-terminal PPPxY motif regulating membrane abundance is shown in **gray**.

```
alpha-ENaC_Homo_sapiens                     MEGNKLEEQDSSPPQSTPGLMKGNKREEQGLGPEPAAPQQPTAEEEALIEFHRSYRELFE  60
alpha-ENaC_Globicephala_melas               --------------------MKGDKHEEPEPGPEPAAPSPSTDEEEPLLEFHHSYRELFQ  40
alpha-ENaC_Lagenorhynchus_obliquidens       --------------------MKGDKHEEPEPGPEPAAPPPSTDEEEPLLEFHHSYRELFQ  40
alpha-ENaC_Tursiops_truncatus               --------------------MKGDKHEEPEPGPEPAAPPPSTDEEEPLLEFHHSYRELFQ  40
alpha-ENaC_Orcinus_orca                     --------------------MKGDKHEEPEPGPEPAAPPPSTDEEEPLLEFHHSYRELFQ  40
alpha-ENaC_Phocoena_sinus                   --------------------MKGDKHEEPEPGPEPAAPPPSTDEEEPLLEFHHSYRELFQ  40
alpha-ENaC_Neophocaena_asiaeorientalis      --------------------MKGDKHEEPEPGPEPAAPPPSTDEEEPLLEFHHSYRELFQ  40
alpha-ENaC_Monodon_monocerons               --------------------MKGDKHEEPEPGPEPAAPPPSTDEEEPLLEFHHSYRELFQ  40
alpha-ENaC_Delphinapterus_leucas            --------------------MKGDKHEEPEPGPEPAAPPPSTDEEEPLLEFHHSYRELFQ  40
alpha-ENaC_Pontoporia_blainvillei           --------------------MKGDKREEPEPGPEPAAPPPSTDEEEALLEFHDSYRELFQ  40
alpha-ENaC_Inia_geoffrensis                 --------------------MKGDKREEPEPGPEPAAPPPSTDEEEALLEFHHSYRELFQ  40
alpha-ENaC_Mesoplodon_bidens                --------------------MKGDKREEPEPGPEPASPPPSTDEEEALLEFHHSYRELFQ  40
alpha-ENaC_Ziphius_cavirostris              --------------------MKGDKREEPEPGPEPAAPPPSTDEEEALLEFHHSYRELFQ  40
alpha-ENaC_Platanista_gangetica             --------------------MKGDKREEPEPGPEPAAPPPPTDEEEALLEFHHSYRELFQ  40
alpha-ENaC_Platanista_minor                 --------------------MKGDKREEPEPGPEPAAPPPPTDEEEALLEFHHSYRELFQ  40
alpha-ENaC_Kogia_breviceps                  --------------------MKGDKREEPEAGPEPAAPAPPMDEEEALLEFHHSYRELFQ  40
alpha-ENaC_Physeter_catodon                 --------------------MKGDEREEPEAGPEPAAPPPPADEQEALLEFHHSYRELFQ  40
alpha-ENaC_Balaenoptera_musculus            --------------------MKGDKREEPEPGPEPAAPPPPTEEEEALLEFHHSYRELFQ  40
alpha-ENaC_Eubalaena_japonica               --------------------MKGDKREEPEPGPEPAAAPPPTEEEEALLEFHHSYRELFQ  40
alpha-ENaC_Hippopotamus_amphibius           --------------------MKEDKREEPGPGPEPEAPPPPTEEEEALLEFHRSYRELFE  40
alpha-ENaC_Tragulus_javanicus               --------------------MKGDKREEPGPGPEPAAPPRPTEDEEALLEFHRSYRELFE  40
alpha-ENaC_Antilocapra_americana            --------------------MKGDKPEEPGPGPEPAGPPPRTEEEEALLEFHRSYRELFE  40
alpha-ENaC_Giraffa_camelopardalis           --------------------MKGDKPEEPGLGPEPSGPLPRTEE-EALLEFHRSYRELFE  39
alpha-ENaC_Giraffa_tippelskirchi            --------------------MKGDKPEEPGLGPEPSGPLPRTEE-EALLEFHRSYRELFE  39
alpha-ENaC_Capreolus_pygargus               --------------------MKGDKPEEPGPGPEPSGPPPPTEEEEALLEFHRSYRELFE  40
alpha-ENaC_Cervus_elaphus                   --------------------MKGDKPEEPGQGPEPSGPPPPTEEEEALLEFHRSYRELFE  40
alpha-ENaC_Moschus_moschiferus              --------------------MKGDKPEELGPGPEPSGPPPPTEEEEALLEFHRSYRELFE  40
alpha-ENaC_Moschus_berezovskii              --------------------MKGDKPEELGPGPEPSGPPPPTEEEEALLEFHRSYRELFE  40
alpha-ENaC_Bos_grunniens                    --------------------MKGDKPEEPGPGPEPSGPPPPTEEEEALLEFHRSYRELFE  40
alpha-ENaC_Bos_taurus                       --------------------MKGDKPEEPGPGPEPSGPPPPTEEEEALLEFHRSYRELFE  40
alpha-ENaC_Bubalus_bubalis                  --------------------MKGDKPEEPGPGPEPSGPLPPTEEEEALLEFHRSYRELFE  40
alpha-ENaC_Nanger_granti                    --------------------MKGDKPEEPGPGPETSGPPPPTEEEEALLEFHRSYRELFE  40
alpha-ENaC_Capra_hircus                     --------------------MKGDKREELGPGPEPSGPLPPTEEEEALLEFHRSYRELFE  40
alpha-ENaC_Ovis_aries                       --------------------MKGDKREELGPGPEPSAPPLPTEEEEALLEFHRSYRELFE  40
alpha-ENaC_Ovis_canadensis                  --------------------MKGDKREELGPGPEPSAPPPPTEEEEALLEFHRSYRELFE  40
alpha-ENaC_Hippotragus_niger                --------------------MKGDKPEEPGPGSEPSGPPQPTEEEEALLEFHRSYRELFE  40
alpha-ENaC_Damaliscus_lunatus               --------------------MKGDKPEEPGPGPEPSGPPPPTEEEEALLEFHRSYRELFE  40
alpha-ENaC_Sus_scrofa                       --------------------MKGDKREEPGPGPEPTAPPLPTEEEEALLEFHRSYRELFQ  40
alpha-ENaC_Vicugna_pacos                    --------------------MKGDKREEPGPDPEPAAPQQPTEEEEALIEFHRSYRELFE  40
alpha-ENaC_Equus_callabus                   --------------------MKGDKHEEQELGSEPTAPQQPTEEEEALIEFHRSYRELFE  40
                                                                ** :: **   . *  .      : * *:*** ******:

                                                                          TM1
alpha-ENaC_Homo_sapiens                     FFCNNTTIHGAIRLVCSQHNRMKTAFWAVLWLCTFGMMYWQFGLLFGEYFSYPVSLNINL  120
alpha-ENaC_Globicephala_melas               FFCNNTTIHGAIRLVCSQHNRMKTAFWAVLWLCTFGMMYWQFGLLFGEYFSYPVSLNINL  100
alpha-ENaC_Lagenorhynchus_obliquidens       FFCNNTTIHGAIRLVCSQHNRMKTAFWAVLWLCTFGMMYWQFGLLFGEYFSYPVSLNINL  100
alpha-ENaC_Tursiops_truncatus               FFCNNTTIHGAIRLVCSQHNRMKTAFWAVLWLCTFGMMYWQFGLLFGEYFSYPVSLNINL  100
alpha-ENaC_Orcinus_orca                     FFCNNTTIHGAIRLVCSQHNRMKTAFWAVLWLCTFGMMYWQFGLLFGEYFSYPVSLNINL  100
alpha-ENaC_Phocoena_sinus                   FFCNNTTIHGAIRLVCSQHNRMKTAFWAVLWLCTFGMMYWQFGLLFGDYFSYPVSLNINL  100
alpha-ENaC_Neophocaena_asiaeorientalis      FFCNNTTIHGAIRLVCSQHNRMKTAFWAVLWLCTFGMMYWQFGLLFGDYFSYPVSLNINL  100
alpha-ENaC_Monodon_monocerons               FFCNNTTIHGAIRLVCSQHNRMKTAFWAVLWLCTFGMMYWQFGLLFGEYFSYPVSLNINL  100
alpha-ENaC_Delphinapterus_leucas            FFCNNTTIHGAIRLVCSQHNRMKTAFWAVLWLCTFGMMYWQFGLLFGEYFSYPVSLNINL  100
alpha-ENaC_Pontoporia_blainvillei           FFCNNTTIHGAIRLVCSQHNRMKTAFWAVLWLCTFGMMYWQFGLLFGEYFSYPVSLNINL  100
alpha-ENaC_Inia_geoffrensis                 FFCSNTTIHGAIRLVCSQHNRMKTAFWAVLWLCTFGMMYWQFGLLFGEYFSYPVSLNINL  100
alpha-ENaC_Mesoplodon_bidens                FFCNNTTIHGAIRLVCSQHNRMKTAFWAVLWLCTFGMMYWQFGLLFGEYFSYPVSLNINL  100
alpha-ENaC_Ziphius_cavirostris              FFCNNTTIHGAIRLVCSQHNRMKTAFWAVLWLCTFGMMYWQFGLLFGEYFSYPVSLNINL  100
alpha-ENaC_Platanista_gangetica             FFCSNTTIHGAIRLVCSQHNRMKTAFWAVLWLCTFGMMYWQFGLLFGEYFSYPVSLNINL  100
alpha-ENaC_Platanista_minor                 FFCSNTTIHGAIRLVCSQHNRMKTAFWAVLWLCTFGMMYWQFGLLFGEYFSYPVSLNINL  100
alpha-ENaC_Kogia_breviceps                  FFCNNTTIHGAIRLVCSQHNRMKTAFWAVLWLCAFSMMYWQFGLLFGEYFSYPVSLNINL  100
```

```
alpha-ENaC_Physeter_catodon                 FFCSNTTIHGAIRLVCSRHNRMKTAFWAALWLCAFGMMYWQFGLLFGEYFSYPVSLNINL  100
alpha-ENaC_Balaenoptera_musculus            FFCNNTTIHGAIRLVCSQHNRMKTAFWAVLWLCTFGMMYWQFGLLFGEYFSYPVSLNINL  100
alpha-ENaC_Eubalaena_japonica               FFCNNTTIHGAIRLVCSQHNRMKTAFWAVLWLCTFGMMYWQFGLLFGEYFSYPVSLNINL  100
alpha-ENaC_Hippopotamus_amphibius           FFCNNTTIHGAIRLVCSQHNRMKTAFWAVLWLCTFGMMYWQFGLLFGEYFSYPVSLNINL  100
alpha-ENaC_Tragulus_javanicus               FFCNNTTIHGAIRLVCSRHNRMKTAFWAVLWLCTFGMMYWQFGQLFGEYFSYPVSLNINL  100
alpha-ENaC_Antilocapra_americana            FFCNNTTIHGAIRLVCSQHNRMKTAFWAVLWLCTFGMMYWQFGQLFGEYFSYPVSLNINL  100
alpha-ENaC_Giraffa_camelopardalis           FFCNNTTIHGAIRLVCSQHNRMKTVFWAVLWLCTFGMMYWQFGQLFGEYFSYPVSLNINL  99
alpha-ENaC_Giraffa_tippelskirchi            FFCNNTTIHGAIRLVCSQHNRMKTVFWAVLWLCTFGMMYWQFGQLFGEYFSYPVSLNINL  99
alpha-ENaC_Capreolus_pygargus               FFCNNTTIHGAIRLVCSQHNRMKTVFWAVLWLCTFGMMYWQFGQLFGEYFSYPVNLNINL  100
alpha-ENaC_Cervus_elaphus                   FFCNNTTIHGAIRLVCSQHNRMKTVFWAVLWLCTFGMMYWQFGQLFGEYFSYPVSLNINL  100
alpha-ENaC_Moschus_moschiferus              FFCNNTTIHGAIRLVCSQHNRMKTVFWAVLWLCTFGMMYWQFGQLFGEYFSYPVSLNINL  100
alpha-ENaC_Moschus_berezovskii              FFCNNTTIHGAIRLVCSQHNRMKTVFWAVLWLCTFGMMYWQFGQLFGEYFSYPVSLNINL  100
alpha-ENaC_Bos_grunniens                    FFCNNTTIHGAIRLVCSQHNRMKTVFWAVLWLCTFGMMYWQFGQLFGEYFSYPVSLNINL  100
alpha-ENaC_Bos_taurus                       FFCNNTTIHGAIRLVCSQHNRMKTVFWAVLWLCTFGMMYWQFGQLFGEYFSYPVSLNINL  100
alpha-ENaC_Bubalus_bubalis                  FFCNNTTIHGAIRLVCSQHNRMKTVFWAVLWLCTFGMMYWQFGQLFGEYFSYPVSLNINL  100
alpha-ENaC_Nanger_granti                    FFCNNTTIHGAIRLVCSQHNRMKTVFWAVLWLCTFGMMYWQFGQLFGEYFSYPVSLNINL  100
alpha-ENaC_Capra_hircus                     FFCNNTTIHGAIRLVCSQHNRMKTVFWAVLWLCTFGMMYWQFGQLFGEYFSYPVSLNINL  100
alpha-ENaC_Ovis_aries                       FFCNNTTIHGAIRLVCSQHNRMKTVFWAVLWLCTFGMMYWQFGQLFGEYFSYPVSLNINL  100
alpha-ENaC_Ovis_canadensis                  FFCNNTTIHGAIRLVCSQHNRMKTVFWAVLWLCTFGMMYWQFGQLFGEYFSYPVSLNINL  100
alpha-ENaC_Hippotragus_niger                FFCNNTTIHGAIRLVCSQHNRMKTVFWAVLWLCTFGMMYWQFGQLFGEYFSYPVSLNINL  100
alpha-ENaC_Damaliscus_lunatus               FFCNNTTIHGAIRLVCSQHNRMKTVFWAVLWLCTFGMMYWQFGQLFGEYFSYPVSLNINL  100
alpha-ENaC_Sus_scrofa                       FFCNNTTIHGAIRLVCSQHNRMKTAFWAVLWLCTFGMMYWQFGLLFEEYFSYPVSLNINL  100
alpha-ENaC_Vicugna_pacos                    FFCNNTTIHGAIRLVCSQHNRMKTAFWAVLWLCTFGMMYWQFGLLFQEYFSYPVSLNINL  100
alpha-ENaC_Equus_callabus                   FFCNHTTIHGAIRLVCSQHNRMKTAFWAVLWLCTFGMMYWQFGLLFGEYFSYPVSLNINL  100
                                            ***.:************:******.***.****:*.******* ** :******.*****

alpha-ENaC_Homo_sapiens                     NSDKLVFPAVTICTLNPYRYPEIKEELEELDRITEQTLFDLYKYSSFTTLVAGSRSRRDL  180
alpha-ENaC_Globicephala_melas               NSEKLVFPAVTVCTLNPYRYTEMKKDLEELDRTTEQTLFDLYKYNSSNSLVAHARGRRDL  160
alpha-ENaC_Lagenorhynchus_obliquidens       NSEKLVFPAVTVCTLNPYRYTEMKKDLEELDRTTEQTLFDLYKYNSSNSLVAHARGRRDL  160
alpha-ENaC_Tursiops_truncatus               NSEKLVFPAVTVCTLNPYRYTEMKKDLEELDRTTEQTLFDLYKYNSSNSLVAHARGRRDL  160
alpha-ENaC_Orcinus_orca                     NSEKLVFPAVTVCTLNPYRYTEMKKDLEELDRTTEQTLFDLYKYNSSNSLVAHARGRRDL  160
alpha-ENaC_Phocoena_sinus                   NSEKLVFPAVTVCTLNPYRYTEMKKDLEELDRITEQTLFDLYKYNSSNTLVAHARGRRDL  160
alpha-ENaC_Neophocaena_asiaeorientalis      NSEKLVFPAVTVCTLNPYRYTEMKKDLEELDRITEQTLFDLYKYNSSNTLVAHARGRRDL  160
alpha-ENaC_Monodon_monocerons               NSEKLVFPAVTVCTLNPYRYTEMKKDLEELDRITEQTLFDLYKYNSSNTLVAHARGRRDL  160
alpha-ENaC_Delphinapterus_leucas            NSEKLVFPAVTVCTLNPYRYTEMKKDLEELDRITEQTLFDLYKYNSSNTLVAHARGRRDL  160
alpha-ENaC_Pontoporia_blainvillei           NSEKLVFPAVTVCTLNPYRYTEMKKDLEELDRITEQTLFGLYKYNSSNTLVAHARGRRDL  160
alpha-ENaC_Inia_geoffrensis                 NSEKLVFPAVTVCTLNPYRYTEMKKDLEELDRITQQTLFDLYEYNSSNTLVAHARGRRDL  160
alpha-ENaC_Mesoplodon_bidens                NSEKLVFPAVTVCTLNPYRYTEIKGVLEELDQITEQTLFDLYKYNSSNTLVAHARGRRDL  160
alpha-ENaC_Ziphius_cavirostris              NSEKLVFPAVTVCTLNPYRYTEIKGELEELDQVTEQTLFDLYKYNSSNTLVAHARGRRDL  160
alpha-ENaC_Platanista_gangetica             NSEKLVFPAVTVCTLNPYRYTEIKEELEELDRITEQTLFDLYKYNSSNTLVAHARGRRDL  160
alpha-ENaC_Platanista_minor                 NSEKLVFPAVTVCTLNPYRYTEIKEELEELDRITEQTLFDLYKYNSSNTLVAHARGRRDL  160
alpha-ENaC_Kogia_breviceps                  NSEKLVFPAVTVCTLNPYRYTEIKEELEELDRITEQTLFDLYKYNSSNTLVAHARGRRDL  160
alpha-ENaC_Physeter_catodon                 NSEKLVFPAVTVCTLNPYRYTEIKEELEELDRITEQTLFDLYKYNSSNTLVAHARGRRDL  160
alpha-ENaC_Balaenoptera_musculus            NSEKLVFPAVTVCTLNPYRYTEIKEELEELDRITEQTLFDLYKYNSSNTLVAHARGRRDL  160
alpha-ENaC_Eubalaena_japonica               NSDKLVFPAVTVCTLNPYRYTEIKEELEELDRITEQTLFDLYKYNSSNTLVAHARGRRDL  160
alpha-ENaC_Hippopotamus_amphibius           NSDKLVFPAVTVCTLNPYRYKEIKEELEELDRITEQTLFDLYKYNSSNTLVAHARGRRDL  160
alpha-ENaC_Tragulus_javanicus               NSDKLVFPAVSICTLNPYRYKEIQDELEELDRITEQTLFDLYKYNSSNTLTAHARPRRDL  160
alpha-ENaC_Antilocapra_americana            NSDKLVFPAVSICTLNPYRYKEIQEELEELDRITEQTLFDLYKYNSSNTLVASARSRRDL  160
alpha-ENaC_Giraffa_camelopardalis           NSDKLIFPAVSICTLNPYRYKEIQEELEELDRITEQTLFDLYKYNSSNTLVA--RSRRDL  157
alpha-ENaC_Giraffa_tippelskirchi            NSDKLIFPAVSICTLNPYRYKEIQEELEELDRITEQTLFDLYKYNSSNTLVA--RSRRDL  157
alpha-ENaC_Capreolus_pygargus               NSDKLIFPAVSICTLNPYRYKEIQEELEELDRITEQTLFDLYEYNSSSSLVARARARRAL  160
alpha-ENaC_Cervus_elaphus                   NSDKLIFPAVSICTLNPYRYKEIQEELEELDRITEQTLFDLYEYNSSNTLVAHARSRRDL  160
alpha-ENaC_Moschus_moschiferus              NSDKLIFPAVSICTLNPYRYKDIQEELEELDRITEQTLFDLYKYNSSNSLVAHARSRREL  160
alpha-ENaC_Moschus_berezovskii              NSDKLIFPAVSICTLNPYRYKDIQEELEELDRITEQTLFDLYKYNSSNSLVAHARSRREL  160
alpha-ENaC_Bos_grunniens                    NSDKLVFPAVSICTLNPYRYKEIQEELEELDRITEQTLFDLYKYNSSKTLVAHARSRRDL  160
alpha-ENaC_Bos_taurus                       NSDKLVFPAVSICTLNPYRYKEIQEELEELDRITEQTLFDLYKYNSSKTLVAHARSRRDL  160
alpha-ENaC_Bubalus_bubalis                  NSDKLVFPAVSICTLNPYRYKEIQEELEELDRITEQTLFDLYKYNSSNTLVAHARSRRDL  160
alpha-ENaC_Nanger_granti                    NSDKLIFPAVSICTLNPYRYKEIQEELEDLDRITEQTLFDLYKYNASHTLVAHARSRRDL  160
alpha-ENaC_Capra_hircus                     NSDKLIFPAVSICTLNPYRYKEIQEELEDLDRITEQTLFDLYKYNASHTLVAHARSRRDL  160
alpha-ENaC_Ovis_aries                       NSDKLIFPAVSICTLNPYRYKEIQEELEDLDRITEQTLFDLYKYNASHTLVAHARSRRDL  160
alpha-ENaC_Ovis_canadensis                  NSDKLIFPAVSICTLNPYRYKEIQEELEDLDRITEQTLFDLYKYNASHTLVAHARSRRDL  160
alpha-ENaC_Hippotragus_niger                NSDKLIFPAVSICTLNPYRYKEIQEELEDLDRITEQTLFDLYKYNSSHTLVAHARSRRDL  160
alpha-ENaC_Damaliscus_lunatus               NSDKLIFPAVSICTLNPYRYKEIQEELEDLDRITEQTLFDLYKYNSSHTLVAHARSRRDL  160
alpha-ENaC_Sus_scrofa                       NSDKLVFPAVTICTLNPYRYKEIKEELEELDRITEQTLFDLYKYNSSNTLVAHARLRRDL  160
alpha-ENaC_Vicugna_pacos                    NSDKLVFPAVSICTLNPYRYTEIKEELEELDRITEQTLFDLYKYNSSNTLVARARGRRDL  160
alpha-ENaC_Equus_callabus                   NSDKLVFPAVTICTLNPYRYAKIKEELEELDRITEQTLYDLYKYNFSNTLVAHPRGRRDL  160
                                            **:**:****::******** .::  **:**: *:***:.**:*.   :*.*  * ** *

alpha-ENaC_Homo_sapiens                     RGTLPHPLQRLRVPPPPHGARRARSVA-SSLRDNNPQVDWKDWKIGFQLCNQNKSDCFYQ  239
alpha-ENaC_Globicephala_melas               RQSLPHPLQRLPVPAPPHAASRVRRSD-SSLSHSNPKVNRKDWKIGFQLCNQNNSDCFYR  219
alpha-ENaC_Lagenorhynchus_obliquidens       RESLPHPLQRLPVPAPPHAASRVRRSD-SSLSHSNPKVNRKDWKIGFQLCNQNNSDCFYR  219
alpha-ENaC_Tursiops_truncatus               RESLPHPLQRLPVPAPPHAASRVRRSD-SSLSHSNPKVNRKDWMIGFQLCNQNNSDCFYR  219
alpha-ENaC_Orcinus_orca                     RESLPHPLQRLPVPAPPHAASRVRRSD-SSLSHSNPKVNRKDWKIGFQLCNQNNSDCFYR  219
alpha-ENaC_Phocoena_sinus                   RESLPHPLQRLQVPAPPHASSRVRRSN-SSLSDNNPQVNRKDWKIGFQLCNQNNSDCFYR  219
alpha-ENaC_Neophocaena_asiaeorientalis      RESLPHPLQRLQVPAPPHAASRVRRSN-SSLSDNNPQVNRKDWKIGFQLCNQNNSDCFYR  219
alpha-ENaC_Monodon_monocerons               RESLPHPLQRLPVPAPPHAASRVRRSD-SSLRDNNPQVNRKDWKIGFQLCNQNNSDCFYR  219
alpha-ENaC_Delphinapterus_leucas            RESLPHPLQRLPVPAPPHAASRVRRSD-SSLRDNNPQVNRKDWKIGFQLCNQNNSDCFYR  219
alpha-ENaC_Pontoporia_blainvillei           RESLPHPLQRLPVPAPPHAASRVRRSG-SSLSDNNPPVKRKDWNIGFKLCNQNNSDCFYQ  219
alpha-ENaC_Inia_geoffrensis                 RESLPHPLQRLPVPAPPHAASRVRRSG-SSLSDNNPQVNRKDWKIGFQLCNQNNSDCFYQ  219
alpha-ENaC_Mesoplodon_bidens                RESLPHPLQRLPVPAPPHAASRVRRSG-SSLEENNPPVNRKDWNVGFRLCNQNKSDCFYQ  219
alpha-ENaC_Ziphius_cavirostris              RESLPHPLQRLPVPAPPHAASRVRRSG-SAVEENNPQVNRKDWKVGFQLCSQNASECFYQ  219
alpha-ENaC_Platanista_gangetica             LESLPHPLQRLPVPAPPHAASRVRRSG-SSLRDNNPQVNRKDWKIGFQLCNQNKSDCFYQ  219
alpha-ENaC_Platanista_minor                 LESLPHPLQRLPVPAPPHAASRVRRSG-SSLRDNNPQVNRKDWKIGFQLCNQNKSDCFYQ  219
alpha-ENaC_Kogia_breviceps                  RESLPHPLQLLPVPAPPRAASRVRRSG-SSLRDNNPQVNRKDWKIGFQLCNQNKSDCFYQ  219
```

```
alpha-ENaC_Physeter_catodon                 RESLPHPLQRLPVPAPPRAASRVRRSG-SSLRDNNPQVNRKDWKIGFQLCNQNKSDCFYQ  219
alpha-ENaC_Balaenoptera_musculus            RESLPHPLQRLPVPAPPHAASRVRRSG-SSLRDNNPQVNRKDWKIGFQLCNQNKSDCFYQ  219
alpha-ENaC_Eubalaena_japonica               RESLPHPLQRVPVPAPPHAASRVRRSG-SSLRDNNPQVNRKDWKIGFQLCNQNKSDCFYQ  219
alpha-ENaC_Hippopotamus_amphibius           REPLPHPLQRLPVAAPPHAARRVRRAG-SSVRDNNPQVNRKDWKIGFQLCNQNKSDCFYQ  219
alpha-ENaC_Tragulus_javanicus               REPLPHPLQRLPVPAPPHAASRVRRAG-SSMRDNNPQVNREDWKIGFQLCNQNKSDCFYK  219
alpha-ENaC_Antilocapra_americana            REPLPHPLQRLPVPAPPHAAGRVRRAG-SSVLDNNPQVNRKDWKIGFQLCNENKSDCFYQ  219
alpha-ENaC_Giraffa_camelopardalis           REPLPHPLQRLPVPAPPHEARRVRHTG-SSVRDNNPQVNRKDWKIGFQLCNQNKSDCFYQ  216
alpha-ENaC_Giraffa_tippelskirchi            REPLPHPLQRLPVPAPPHEARRVRHTG-SSVRDNNPQVNRKDWKIGFQLCNQNKSDCFYQ  216
alpha-ENaC_Capreolus_pygargus               RRPLPHPLRRLPVPAPPHAARRARRAG-SSVQDNNPQVNRKDWKIGFQLCNQNKSDCFYQ  219
alpha-ENaC_Cervus_elaphus                   RKPLPHPLQRLPVPAPPHEARKVRRAG-SSVRDNNPQVNRKDWKIGFQLCNQNKSDCFYQ  219
alpha-ENaC_Moschus_moschiferus              REPLPHPLQRLPVRTSPHAARRVRRPG-SSVRDNNPQVNRKDWKIGFQLCNQNKSDCFYQ  219
alpha-ENaC_Moschus_berezovskii              REPLPHPLQRLPVPTSPHAARRVRRPG-SSVRDNNPQVNRKDWKIGFQLCNQNKSDCFYQ  219
alpha-ENaC_Bos_grunniens                    REPLPHPLQRLPVPAPSHAARGVRRAG-SSMRDNNPQVNRKDWKIGFQLCNQNKSDCFYQ  219
alpha-ENaC_Bos_taurus                       REPLPHPLQRLPVPAPPHAARGVRRAG-SSMRDNNPQVNRKDWKIGFQLCNQNKSDCFYQ  219
alpha-ENaC_Bubalus_bubalis                  REPLPHPLQRLPVPAPPHAARGVRRAG-SSVRDNNPQVNRKDWKIGFQLCNQNKSDCFYQ  219
alpha-ENaC_Nanger_granti                    REPLPHPLQRLPIPAPPNAARRARRAG-SSVRDNNPQVNRKDWKIGFQLCNQNKSDCFYQ  219
alpha-ENaC_Capra_hircus                     REPLPHPLQRLPIPAPPHAARRVRRAG-SSVRDNNPQVNRKDWKIGFQLCNQNKSDCFYQ  219
alpha-ENaC_Ovis_aries                       REPLPHPLQRLPIPAPPHAARRVRRAG-SSVRDNNPQVNRKDWKIGFQLCNQNKSDCFYQ  219
alpha-ENaC_Ovis_canadensis                  REPLPHPLQRLPIPAPPHAARRVRRAG-SSVRDNNPQVNRKDWKIGFQLCNQNKSDCFYQ  219
alpha-ENaC_Hippotragus_niger                REPLPHPLQRLPIPAPPHAARRVHRTG-SSVRDNNPQVNRKDWKIGFQLCNQNKSDCFYQ  219
alpha-ENaC_Damaliscus_lunatus               REPLPHPLQRLPIPAPPHAARRVHRAG-SSVRDNNPQVNRKDWKIGFQLCNQNKSDCFYQ  219
alpha-ENaC_Sus_scrofa                       REPLPHPLQRLTVPAPPHSARRVRSATSSSVRDNNPQVNRKDWKIGFQLCNQNKSDCFYQ  220
alpha-ENaC_Vicugna_pacos                    REALPHPLQRLPVPAPPHAARSARSA-TSSVRDNNPKVNRKDWKIGFQLCNRNKSDCFYQ  219
alpha-ENaC_Equus_callabus                   GETLPHPLQRLPGPAPPHEARRARMA--SSVRDNNPQVNRKDWKIGFQLCNQNKSDCFYQ  218
                                               *****: :      . :  .:    *:: ..** *. :** :**:**..* *:***:

alpha-ENaC_Homo_sapiens                     TYSSGVDAVREWYRFHYINILSRLP-ETLPSLEEDTLGNFIFACRFNQVSCNQANYSHFH  298
alpha-ENaC_Globicephala_melas               TYSSGVDAVREWYRFHYINILSRRQ-D-TPMLEEEALGKFIFACRFNQVSCNEANYSHFH  277
alpha-ENaC_Lagenorhynchus_obliquidens       TYSSGVDAVREWYRFHYINILSRRQ-D-TPMLEEEALGKFIFACRFNQVSCNEANYSHFH  277
alpha-ENaC_Tursiops_truncatus               TYSSGVDAVREWYRFHYINILSRRQ-D-TPMLEEEALGKFIFACRFNQVSCNEANYSHFH  277
alpha-ENaC_Orcinus_orca                     TYSSGVDAVREWYRFHYINILSRRQ-D-TPMLEEEALGKFIFACRFNQVSCNEANYSHFH  277
alpha-ENaC_Phocoena_sinus                   TYSSGVDAVREWYRFHYINILSRRQ-D-TPLLEEEALGKFIFACRFNQVSCNEANYSHFH  277
alpha-ENaC_Neophocaena_asiaeorientalis      TYSSGVDAVREWYRFHYINILSRRQ-D-TPLLEEEALGKFIFACRFNQVSCNEANYSHFH  277
alpha-ENaC_Monodon_monocerons               TYSSGVDAVREWYRFHYINILSRRQ-D-TPLLEEEALGKFIFACRFNQVSCNEANYSHFH  277
alpha-ENaC_Delphinapterus_leucas            TYSSGVDAVREWYRFHYINILSRRQ-D-TPLLEEEALGKFIFACRFNQVSCNEANYSHFH  277
alpha-ENaC_Pontoporia_blainvillei           TYSSGVDAVREWYRFHYINILSRRQ-D-TPLLEEEALGDFIFACRFNQVSCDEANYSRFH  277
alpha-ENaC_Inia_geoffrensis                 TYSSGVDAVREWYRFHYINILSRRQ-D-TPLLEEEALGDFIFACRFNQVSCNEANYSRFH  277
alpha-ENaC_Mesoplodon_bidens                RYSSGVDAVREWYRFHYINILSRRR-D-SPLLEEEALGKFIFACRFNQFSCNEANYSRFH  277
alpha-ENaC_Ziphius_cavirostris              RYSSGVDAVREWYRFHYINILSRRR-D-SPLLEEEALGKFIFACRFNQFSCNEANYSRFH  277
alpha-ENaC_Platanista_gangetica             TYSSGVDAVREWYRFHYINILSRRQ-DTSPSLEEDALGKFIFACRFNQFSCNEANYSHFH  278
alpha-ENaC_Platanista_minor                 TYSSGVDAVREWYRFHYINILSRRQ-DTSPSLEEDALGKFIFACRFNQFSCNEANYSHFH  278
alpha-ENaC_Kogia_breviceps                  TYSSGVDAVREWYRFHYINILSRRR-DPSPVLEEDALGKFIFACRFNQASCNEANYSHFH  278
alpha-ENaC_Physeter_catodon                 TYSSGVDAVREWYRFHYINILSRRR-DPSPPLEEDALGKFIFACRFNQASCNEANYSHFH  278
alpha-ENaC_Balaenoptera_musculus            TYSSGVDAVREWYRFHYINILSRRQ-DTSPSLEEDALGKFIFACRFNQASCNEANYSHFH  278
alpha-ENaC_Eubalaena_japonica               TYSSGVDAVREWYRFHYINILSRRQ-DTSPSLEEDALGKFIFACRFNQASCNEANYSHFH  278
alpha-ENaC_Hippopotamus_amphibius           TYSSGVDAVREWYRFHYINILSRRK-DTSSSLEEDVLGKFIFTCRFNQVSCNEANYSHFH  278
alpha-ENaC_Tragulus_javanicus               TYSSGVDAVREWYRFHYINILARRQQDTSPLLEEDVLGKFIFTCRFNQNSCNEANYSHFH  279
alpha-ENaC_Antilocapra_americana            KYSSGVDAVREWYRFHYINILSRRRQDTSPSLEEDVLGKFIFTCRFNQDSCNEANYSHFH  279
alpha-ENaC_Giraffa_camelopardalis           TYSSGVDAVREWYRFHYINILSRRRQDTSPSLEEDVLGKFIFTCRFNQDSCNEANYSHFH  276
alpha-ENaC_Giraffa_tippelskirchi            TYSSGVDAVREWYRFHYINILSRRRQDTSPSLEEDVLGKFIFTCRFNQDSCNEANYSHFH  276
alpha-ENaC_Capreolus_pygargus               TYSSGVDAVREWYRFHYINILSRRKQDTSPSLEEDVLGKFIFTCRFNQDSCNEANYSHFH  279
alpha-ENaC_Cervus_elaphus                   TYSSGVDAVREWYRFHYINILSRRRQDTSPSLEEDVLGKFIFTCRFNQDSCNEANYSHFH  279
alpha-ENaC_Moschus_moschiferus              TYSSGVDAVREWYRFHYINILSRRRQDTSPSLEEDVLGKFIFSCRFNQDSCNEANYSHFH  279
alpha-ENaC_Moschus_berezovskii              TYSSGVDAVREWYRFHYINILSRRRQDTSPSLEEDVLGKFIFSCRFNQDSCNEANYSHFH  279
alpha-ENaC_Bos_grunniens                    TYSSGVDAVREWYRFHYINILSRRRQDTSPSLEDDVLGKFIFTCRFNQDSCSEANYSHFH  279
alpha-ENaC_Bos_taurus                       TYSSGVDAVREWYRFHYINILSRRRQDTSPSLEEDVLGKFIFTCRFNQDSCNEANYSHFH  279
alpha-ENaC_Bubalus_bubalis                  TYSSGVDAVTEWYRFHYINILSRRRQDTSPSLEEDVLGKFIFTCRFNQDSCNEANYSHFH  279
alpha-ENaC_Nanger_granti                    TYSSGVDAVREWYRFHYINILSRRRQDTSPSLEEDVLGKFIFTCRFNQDSCNEANYSHFH  279
alpha-ENaC_Capra_hircus                     TYSSGVDAVREWYRFHYINILSRRRQDTSPSLEEDVLGKFIFTCRFNQDSCNEANYSHFH  279
alpha-ENaC_Ovis_aries                       TYSSGVDAVREWYRFHYINILSRRRQDTSPSLEEDVLGKFIFTCRFNQDSCNEANYSHFH  279
alpha-ENaC_Ovis_canadensis                  TYSSGVDAVREWYRFHYINILSRRRRDTSPSLEEDVLGKFIFTCRFNQDSCNEANYSHFH  279
alpha-ENaC_Hippotragus_niger                TYSSGVDAVREWYRFHYINILSRRRQDTSPSLEEDVLGKFIFTCRFNQDSCNEANYSHFH  279
alpha-ENaC_Damaliscus_lunatus               TYSSGVDAVREWYRFHYINILSRRRQDTSPSLEEDVLGKFIFTCRFNQDSCNEANYSHFH  279
alpha-ENaC_Sus_scrofa                       TYSSGVDAVREWYRFHYINILSRRR-DTSPSLEEDALGKFIFACRFNQVSCNEANYSHFH  279
alpha-ENaC_Vicugna_pacos                    TYSSGVDAVREWYRFHYINILARRQ-DTSPSLEEDALGKFIFACRFNQVSCNEANYSHFH  278
alpha-ENaC_Equus_callabus                   TYSSGVDAVREWYRFHYINILSRLPVDASVSLEKDQLDNFIFACRFNQVSCNQANYSHFH  278
                                             ******** ***********:*    :    **.: *..***:***** **.:****:**

alpha-ENaC_Homo_sapiens                     HPMYGNCYTFNDKNNSNLWMSSMPGINNGLSLMLRAEQNDFIPLLSTVTGARVMVHGQDE  358
alpha-ENaC_Globicephala_melas               HPMYGNCYTFNDKNNSNLWMSFMPGVNNGLSLMLRTEQNDFIPLLSTVTGARVMVHGQDE  337
alpha-ENaC_Lagenorhynchus_obliquidens       HPMYGNCYTFNDKNNSNLWMSFMPGVNNGLSLMLRTEQNDFIPLLSTVTGARVMVHGQDE  337
alpha-ENaC_Tursiops_truncatus               HPMYGNCYTFNDKNNSNLWMSFMPGVNNGLSLMLRTEQNDFIPLLSTVTGARVMVHGQDE  337
alpha-ENaC_Orcinus_orca                     HPMYGNCYTFNDKNNSNLWMSFMPGVNNGLSLMLRTEQNDFIPLLSTVTGARVMVHGQDE  337
alpha-ENaC_Phocoena_sinus                   HPMYGNCYTFNDKNTSNLWMSFMPGVNNGLSLTLRTEQNDFIPLLSTVTGARVMVHGQDE  337
alpha-ENaC_Neophocaena_asiaeorientalis      HPMYGNCYTFNDKNTSNLWMSFMPGVNNGLSLTLRTEQNDFIPLLSTVTGARVMVHGQDE  337
alpha-ENaC_Monodon_monocerons               HPMYGNCYTFNNKNNSNLWMSFMPGVNNGLSLTLRTEQNDFIPLLSTVTGARVMVHGQDE  337
alpha-ENaC_Delphinapterus_leucas            HPMYGNCYTFNNKNNSNLWMSFMPGVNNGLSLTLRTEQNDFIPLLSTVTGARVMVHGQDE  337
alpha-ENaC_Pontoporia_blainvillei           HPMYGNCYTFNHKNSSKLWMSFMPGVNNGLSLTLRTEQNDFIPLLSTVTGARVMVHGQDE  337
alpha-ENaC_Inia_geoffrensis                 HPMYGNCYTFNDKNSSNLWMSFMPGVNNGLSLTLRTEQNDFIPLLSTVTGARVMVHGQDE  337
alpha-ENaC_Mesoplodon_bidens                HPMYGNCYTFNDKNSSNLWMSSMPGVNNGLSLTLRTEQNDFIPLLSTVTGARVMVHGQDE  337
alpha-ENaC_Ziphius_cavirostris              HPMYGNCYTFNDKNSSNLWMSSMPGVNNGLSLTLRTEQNDFIPLLSTVTGARVMVHGQDE  337
alpha-ENaC_Platanista_gangetica             HPMYGNCYTFNDKNSSNLWMSSMPGVNNGLSLTLRTEQNDFIPLLSTVTGARVMVHGQDE  338
alpha-ENaC_Platanista_minor                 HPMYGNCYTFNDKNSSNLWMSSMPGVNNGLSLTLRTEQNDFIPLLSTVTGARVMVHGQDE  338
alpha-ENaC_Kogia_breviceps                  HPMYGNCYTFNDKNSSNLWMSSMTGVNNGLSLMLRTEQNDFIPPLSTVTGARVMVHGQDE  338
```

```
alpha-ENaC_Physeter_catodon                   HPMYGNCYTFNDKNSSNLWMSSMTGVNNGLSLTLRTEQNDFIPLLSTVTGARVMVHGQDE  338
alpha-ENaC_Balaenoptera_musculus              HPMYGNCYTFNDKNSSNLWMSSISGVNNGLSLTLRTEQNDFIPLLSTVTGARVMVHGQDE  338
alpha-ENaC_Eubalaena_japonica                 HPMYGNCYTFNDKNSSNLWMSSISGVNNGLSLTLRTEQNDFIPLLSTVTGARVMVHGQDE  338
alpha-ENaC_Hippopotamus_amphibius             HPIYGNCYTFNDKNSSNLWMSSMPGVNNGLSLTLRTEQNDFIPLLSTVTGARVMVHGQDE  338
alpha-ENaC_Tragulus_javanicus                 HPMYGNCYTFNDKNSSNRWMSSRPGVSNGLSLTLRTEQNDFIPLLSTVTGARVMVHERDE  339
alpha-ENaC_Antilocapra_americana              HPMFGNCYTFNDKNSSNLWMSSMPGVNNGLSLTLRTEQNDFIPLLSTVTGARVMVHERDE  339
alpha-ENaC_Giraffa_camelopardalis             HPMYGNCYTFNDKNSSNLWMSSMPGVNNGLSLTLRTEQNDFIPLLSTVTGARVMVHERDE  336
alpha-ENaC_Giraffa_tippelskirchi              HPMYGNCYTFNDKNSSNLWMSSMPGVNNGLSLTLRTEQNDFIPLLSTVTGARVMVHERDE  336
alpha-ENaC_Capreolus_pygargus                 HPMYGNCYTFNDKNSSNLWMSSMPGVNNGLSLTLRTEQNDFIPLLSTVTGARVMVHERDE  339
alpha-ENaC_Cervus_elaphus                     HPMYGNCYTFNDKNSSNLWMSSMPGVNNGLSLTLRTEQNDFIPLLSTVTGARVMVHERDE  339
alpha-ENaC_Moschus_moschiferus                HPMYGNCYTFNDKNSSTLWMSSMPGVNNGLSLTLRTEQNDFIPLLSTVTGARVMVHERDE  339
alpha-ENaC_Moschus_berezovskii                HPMYGNCYTFNDKNSSTLWMSSMPGVNNGLSLTLRTEQNDFIPLLSTVTGARVMVHERDE  339
alpha-ENaC_Bos_grunniens                      HPMYGNCYTFNDKNSSNLWMSSMPGVNNGLSLTLRTEQNDFIPLLSTVTGARVMVHERDE  339
alpha-ENaC_Bos_taurus                         HPMYGNCYTFNDKNSSNLWMSSMPGVNNGLSLTLRTEQNDFIPLLSTVTGARVMVHERDE  339
alpha-ENaC_Bubalus_bubalis                    HPMYGNCYTFNDKNNSNLWMSSMPGVNNGLSLTLRTEQNDFIPLLSTVTGARVMVHERDE  339
alpha-ENaC_Nanger_granti                      HPMYGNCYTFNDKNSSNLWMSSMPGVNNGLSLTLRTEQNDFIPLLSTVTGARVMVHERDE  339
alpha-ENaC_Capra_hircus                       HPMYGNCYTFNDKNSSNLWMSSMPGVNNGLSLTLRTEQNDFIPLLSTVTGARVMVHERDE  339
alpha-ENaC_Ovis_aries                         HPMYGNCYTFNDKNSSNLWISSVPGVNNGLSLTLRTEQNDFIPLLSTVTGARVMVHERDE  339
alpha-ENaC_Ovis_canadensis                    HPMYGNCYTFNDKNSSNLWISSVPGVNNGLSLTLRTEQNDFIPLLSTVTGARVMVHERDE  339
alpha-ENaC_Hippotragus_niger                  HPMYGNCYTFNDKNSSNLWMSSMAGVNNGLSLTLRTEQNDFIPLLSTVTGARVMVHERDE  339
alpha-ENaC_Damaliscus_lunatus                 HPMYGNCYTFNDKNSSNLWMSSVPGVNNGLSLTLRTEQNDFIPLLSTVTGARVMVHERDE  339
alpha-ENaC_Sus_scrofa                         HPIYGNCYTFNDKNSSNLWMSSMPGVNNGLSLTLRTEQNDFIPLLSTVTGARVMVHGQDE  339
alpha-ENaC_Vicugna_pacos                      HPIYGNCYTFNDKNSSNLWMSSMPGVNHGLSLTLRTEQNDFIPLVSTVTGARVMVHGQNE  338
alpha-ENaC_Equus_callabus                     HPMYGNCYTFNDKNNSNLWMSSMPGINNGLSLTLRTEQNDFIPLLSTVTGARVMVHGQDE  338
                                              **::*******.**.*. *:*   *:.:**** **:******* :*********** ::*

alpha-ENaC_Homo_sapiens                       PAFMDDGGFNLRPGVETSISMRKETLDRLGGDYGDCTKNGSDVPVENLYPSKYTQQVCIH  418
alpha-ENaC_Globicephala_melas                 PPFMDDGGFNLRPGMETSISMSKEAVERLGGDYGDCTKNGSDVPVENLYGTKYTQQVCIH  397
alpha-ENaC_Lagenorhynchus_obliquidens         PAFMDDGGFNLRPGMETSISMSKEAVDRLGGDYGDCTKNGSDVPVENLYGTKYTQQVCIH  397
alpha-ENaC_Tursiops_truncatus                 PAFMDDGGFNLRPGMETSISMSKEAVDRLGGDYGDCTKNGSDVPVENLYGTKYTQQVCIH  397
alpha-ENaC_Orcinus_orca                       PAFMDDGGFNLRPGMETSISMSKEAVDRLGGDYGDCTKNGSDVPVENLYGTKYTQQVCIH  397
alpha-ENaC_Phocoena_sinus                     PAFMDDGGFNLRPGVETSISMSKEAVDRLGGDYGDCTKNGSDVPVENLYGTKYTQQVCIH  397
alpha-ENaC_Neophocaena_asiaeorientalis        PAFMDDGGFNLRPGVETSISMSKEAVDRLGGDYGDCTKNGSDVPVENLYGTKYTQQVCIH  397
alpha-ENaC_Monodon_monocerons                 PAFMDDGGFNLRPGMETSISMSKEAVDRLGGDYGDCTKNGSDVPVENLYGTKYTQQVCIH  397
alpha-ENaC_Delphinapterus_leucas              PAFMDDGGFNLRPGMETSISMSKEAVDRLGGDYGDCTKNGSDVPVENLYGTKYTQQVCIH  397
alpha-ENaC_Pontoporia_blainvillei             PAFMDDGGFNLRPGMETSISMSKEAVGRLGGDYGDCTKNGSDVPVENLYGTKYSQQVCIH  397
alpha-ENaC_Inia_geoffrensis                   PAFMDDGGFNLRPGMETSISMSKEAMDRLGGDYGDCTKNGSDVPVENLYGTKYTQQVCIY  397
alpha-ENaC_Mesoplodon_bidens                  PAFMDDGGFNLRPGMETSISMSKETMDRLGGDYGDCTKNGSDVPVENLYGTKYTQQVCIH  397
alpha-ENaC_Ziphius_cavirostris                PAFMDDGGFNLRPGMETSISMSKETMDRLGGDYGDCTKNGSDVPVENLYGTKYTQQVCIH  397
alpha-ENaC_Platanista_gangetica               PAFMDDGGFNLRPGMETSISMSKETVDRLGGDYGDCTKNGSDVPVENLYGAKYTQQVCIH  398
alpha-ENaC_Platanista_minor                   PAFMDDGGFNLRPGMETSISMSKETVDRLGGDYGDCTKNGSDVPVENLYGAKYTQQVCIH  398
alpha-ENaC_Kogia_breviceps                    PAFMDDGGFNLRPGMETSISMSKEAVKRLGGDYGDCTKNGSDVPVENLYGTKYTQQVCIH  398
alpha-ENaC_Physeter_catodon                   PAFMDDGGFNLRPGMETSISMSKEAVKRLGGDYGDCTKNGSDVPVENLYGTKYTQQVCIH  398
alpha-ENaC_Balaenoptera_musculus              PAFMDDGGFNLRPGMETSISMSKEAVARLGGDYGDCTKNGSDVPVENLYGTKYTQQVCIH  398
alpha-ENaC_Eubalaena_japonica                 PAFMDDGGFNLRPGMETSISMSKEAVERLGGNYGDCTKNGSDVPVENLYGTKYTQQVCIH  398
alpha-ENaC_Hippopotamus_amphibius             PAFMDDGGFNLRPGVETSISMSKEAVDRLGGDYGDCTKNGSEVPVKNLYGTKYTQQVCIH  398
alpha-ENaC_Tragulus_javanicus                 PAFMDDAGFNLRPGVETSISMRKEVVGRLGGDYGDCTRNGSEVPVENLYNTKYTQQVCIR  399
alpha-ENaC_Antilocapra_americana              PAFMDDAGFNLRPGVETSISMSKEAVDRLGGDYGDCTKNGSDIPVENLYNTKYTQQVCIH  399
alpha-ENaC_Giraffa_camelopardalis             PAFMDDAGFNLRPGVETSISMSKEAVDRLGGDYGDCTKNGSEVPVENLYNTKYTQQVCIH  396
alpha-ENaC_Giraffa_tippelskirchi              PAFMDDAGFNLRPGVETSISMSKEAVDRLGGDYGDCTKNGSEVPVENLYNTKYTQQVCIH  396
alpha-ENaC_Capreolus_pygargus                 PAFMDDAGFNLRPGVETSISMSKEAVDRLGGDYGDCTKNGSEVPVENLYNTKYTQQVCIH  399
alpha-ENaC_Cervus_elaphus                     PAFMDDSGFNLRPGVETSISMSKEAVDRLGGDYGDCTKNGSEVPVENLYNTKYTQQVCIH  399
alpha-ENaC_Moschus_moschiferus                PAFMDDAGFNLRPGVETSISMSKEAVDRLGGDYGDCTKNGSEVPVENLYNTKYTQQVCIH  399
alpha-ENaC_Moschus_berezovskii                PAFMDDAGFNLRPGVETSISMSKEAVDRLGGDYGDCTKNGSEVPVENLYNTKYTQQVCIH  399
alpha-ENaC_Bos_grunniens                      PAFMDDAGFNLRPGVETSISMSKEAVDRLGGDYGDCTKNGSEVPVENLYNTKYTQQVCIH  399
alpha-ENaC_Bos_taurus                         PAFMDDAGFNLRPGVETSISMSKEAVDRLGGDYGDCTKNGSEVPVENLYNTKYTQQVCIH  399
alpha-ENaC_Bubalus_bubalis                    HPFMDDAGFNLRPGVETSISISKEAVDRLGGDYGDCTKNGSEVPVENLYNTKYTQQVCIH  399
alpha-ENaC_Nanger_granti                      PAFMDDAGFNLRPGVETSISMSKEAVDRLGGDYGDCTKNGSEVPVENLYNTKYTQQVCIH  399
alpha-ENaC_Capra_hircus                       PAFMDDAGFNLRPGVETSISMSKEALDRLGGDYGDCTKNGSEVPVENLYNTKYTQQVCIH  399
alpha-ENaC_Ovis_aries                         PAFMDDAGFNLRPGVETSISMSKEAVDRLGGDYGDCTKNGSEVPVENLYNTKYTQQVCIH  399
alpha-ENaC_Ovis_canadensis                    PAFMDDAGFNLRPGVETSISMSKEAVDRLGGDYGDCTKNGSEVPVENFYNTKYTQQVCIH  399
alpha-ENaC_Hippotragus_niger                  PAFMDDAGFNLRPGVETSISMSKEAVDRLGGDYGDCTKNGSEVPVENLYNTKYTQQVCIH  399
alpha-ENaC_Damaliscus_lunatus                 PAFMDDAGFNLRPGVETSISMSKEAVDRLGGDYGDCTKNGSEVPVENLYNTKYTQQVCIH  399
alpha-ENaC_Sus_scrofa                         PAFMDDGGFNLRPGVESSISMSKEAVDRLGGDYSDCTKNGSEVPVKNLYGSKYTQQVCIH  399
alpha-ENaC_Vicugna_pacos                      PAFMDDGGFNLRPGVETSISMSKEAVDRLGDNYGDCTENGSEIPVENLYLTKYTQQVCIH  398
alpha-ENaC_Equus_callabus                     PAFMDDGGFNLRPGVETSISMRKETLDRLGGTYGDCTKNGSDIPVQNLYGSKYTQQVCIY  398
                                                ****.*******:*:***: **.: ***. *.***.***::**:*:* :**:*****

alpha-ENaC_Homo_sapiens                       SCFQESMIKECGCAYIFYPRPQNVEYCDYRKHSSWGYCYYKLQVDFSSDHLGCFTKCRKP  478
alpha-ENaC_Globicephala_melas                 SCFQVNMIRECGCAYIFYPQPRGVEFCDYRKHNSWGYCYYKLQDAFSSDRLGCFTKCRKP  457
alpha-ENaC_Lagenorhynchus_obliquidens         SCFQVNMIRECGCAYIFYPQPRGVEFCDYRKHNSWGYCYYKLQDAFSSDRLGCFTKCRKP  457
alpha-ENaC_Tursiops_truncatus                 SCFQVNMIRECGCAYIFYPQPRGVEFCDYRKHNSWGYCYYKLQDAFSSDRLGCFTKCRKP  457
alpha-ENaC_Orcinus_orca                       SCFQVNMIRECGCAYIFYPQPRGVEFCDYRKHNSWGYCYYKLQDAFSSDRLGCFTKCRKP  457
alpha-ENaC_Phocoena_sinus                     SCFQVNMIRECGCAYIFYPQPRGVEFCDYRKHNSWGYCYYKLQDAFSSDRLGCFTKCRKP  457
alpha-ENaC_Neophocaena_asiaeorientalis        SCFQVNMIRECGCAYIFYPQPRGVEFCDYRKHNSWGYCYYKLQDAFSSDRLGCFTKCRKP  457
alpha-ENaC_Monodon_monocerons                 SCFQVNMIRECGCAYIFYPQPRGVEFCDYRKHNSWGYCYYKLQDAFSSDRLGCFTKCRKP  457
alpha-ENaC_Delphinapterus_leucas              SCFQVNMIRECGCAYIFYPQPRGVEFCDYRKHNSWGYCYYKLQDAFSSDRLGCFTKCRKP  457
alpha-ENaC_Pontoporia_blainvillei             SCFQVNMIRECGCAYIFYPQPRGVEFCDYRKHNSWGYCYYKLQDAFSSDRLGCFTKCRKP  457
alpha-ENaC_Inia_geoffrensis                   SCFQVNMIRECGCAYIFYPQPRGVEFCDYRKHNSWGYCYYKLQDAFSSDRLGCFTKCRKP  457
alpha-ENaC_Mesoplodon_bidens                  SCFQVNMVRECGCAYIFYPPHRGAEFCDYRKHTSWGYCYYKLQDAFSSDRLGCFTKCRKP  457
alpha-ENaC_Ziphius_cavirostris                SCFQVNMVRECGCAYIFYPPHRGAEFCDYRKHTSWGYCYYKLQDAFSSDRLGCFTKCRKP  457
alpha-ENaC_Platanista_gangetica               SCFQVNMIRECGCAYIFYPLHRGVEFCDYRKHNSWGYCYYKLQDAFSSDRLGCFTKCRKP  458
alpha-ENaC_Platanista_minor                   SCFQVNMIRECGCAYIFYPLHRGVEFCDYRKHNSWGYCYYKLQDAFSSDRLGCFTKCRKP  458
alpha-ENaC_Kogia_breviceps                    SCFQVNMVRECGCAYIFYPPHRGAEFCDYRKHNSWGYCYYKLQDAFSSDRLGCFTKCRKP  458
```

```
alpha-ENaC_Physeter_catodon                SCFQVNMIRECGCAYIFYPLHRGVEFCDYRKHSSWGYCYYKLQDAFSSDRLGCFTKCRKP  458
alpha-ENaC_Balaenoptera_musculus           SCFQVNMIRECGCAYIFYPPPRGVEFCDYRKHNSWGYCYYKLQDAFSSDRLGCFTKCRKP  458
alpha-ENaC_Eubalaena_japonica              SCFQVNMIRECGCAYIFYPPHRGVEFCDYRKHNSWGYCYYKLQDAFSSDRLGCFTKCRKP  458
alpha-ENaC_Hippopotamus_amphibius          SCFQESMIRECGCAYIFYPPSNNSEFCDYRKHNSWGYCYYKLQDAFSSDRLGCFTKCRKP  458
alpha-ENaC_Tragulus_javanicus              SCFQKSMIKKCGCAYILYPRPEGVEFCDYRKHSSWGYCYYKLQDAFSSDRLGCFTKCRKP  459
alpha-ENaC_Antilocapra_americana           SCFQESMIKECGCAYIFYPRPDGVEFCDYRKHNSWGYCYYKLQDAFSSDRLGCFTKCRKP  459
alpha-ENaC_Giraffa_camelopardalis          SCFQESMIKECGCAYIFYPRPHGVEFCDYRKHNSWGYCYYKLQDAFSSDRLGCFTKCRKP  456
alpha-ENaC_Giraffa_tippelskirchi           SCFQESMIKECGCAYIFYPRPHGVEFCDYRKHNSWGYCYYKLQDAFSSDRLGCFTKCRKP  456
alpha-ENaC_Capreolus_pygargus              SCFQESMIQECGCAYIFYPRPDGVEFCDYRKHNSWGYCYYKLQDAFSSDRLGCFTKCRKP  459
alpha-ENaC_Cervus_elaphus                  SCFQESMIKECGCAYIFYPRPDGVEFCDYRKHNSWGYCYYKLQDAFSSDRLGCFTKCRKP  459
alpha-ENaC_Moschus_moschiferus             SCFQESMIKECGCAYIFYPRPDGVEFCDYRKHNSWGYCYYKLQDAFSSDRLGCFTKCRKP  459
alpha-ENaC_Moschus_berezovskii             SCFQESMIKECGCAYIFYPRPDGVEFCDYRKHNSWGYCYYKLQDAFSSDRLGCFTKCRKP  459
alpha-ENaC_Bos_grunniens                   SCFQESMIKECGCAYIFYPRPDGVEFCDYRKHNSWGYCYYKLQDAFSSDRLGCFTKCRKP  459
alpha-ENaC_Bos_taurus                      SCFQESMIKECGCAYIFYPRPDGVEFCDYRKHNSWGYCYYKLQDAFSSDRLGCFTKCRKP  459
alpha-ENaC_Bubalus_bubalis                 SCFQESMIKECGCAYIFYPLPDGVEFCDYRKHSSWGYCYYKLQDAFSSDRLGCFTKCRKP  459
alpha-ENaC_Nanger_granti                   SCFQESMIKKCGCAYIFYPKPEGVEFCDYKKHDSWGYCYYKLQDAFSSDRLGCFTKCRKP  459
alpha-ENaC_Capra_hircus                    SCFQESMIKECGCAYIFYPRPEGVEFCDYKKHNSWGYCYYKLQDAFSSDRLGCFTKCRKP  459
alpha-ENaC_Ovis_aries                      SCFQESMIKECGCAYIFYPRPEGVEFCDYKKHNSWGYCYYKLQDAFSSDRLGCFTKCRKP  459
alpha-ENaC_Ovis_canadensis                 SCFQESMIKECGCAYIFYPRPEGVEFCDYKKHNSWGYCYYKLQDAFSSDRLGCFTKCRKP  459
alpha-ENaC_Hippotragus_niger               SCFQESMIKECGCAYIFYPRPEGVEFCDYKKHNSWGYCYYKLQDAFSSDRLGCFTKCRKP  459
alpha-ENaC_Damaliscus_lunatus              SCFQESMIKECGCAYIFYPRPEGIEFCDYKKHNSWGYCYYKLQDAFSSDRLGCFTKCRKP  459
alpha-ENaC_Sus_scrofa                      SCFQQNMVKECGCAYIFYPLPPGMEFCDYRKHNSWGYCYYKLQDAFSSDRLGCFTKCRKP  459
alpha-ENaC_Vicugna_pacos                   SCFQESMVRECGCAYIFYPRPHNVDFCDYRKHDSWGYCYYKLQDAFSSDRLGCFTKCRKP  458
alpha-ENaC_Equus_callabus                  SCFQENMIKECGCAYLFYPPLPGVDFCDYRKHNSWGYCYYKLQDAFASNRLGCFTKCRKP  458
                                           **** .*:::*****::**   . ::***:** **********  *:*::**********

alpha-ENaC_Homo_sapiens                    CSVTSYQLSAGYSRWPSVTSQEWVFQMLSRQNNYTVNNKRNGVAKVNIFFKELNYKTNSE  538
alpha-ENaC_Globicephala_melas              CNMTTYKLSAGYSRWPSVTSQDWVFQMLSLQNNYTVKNKRDGIAKLNIFFKELNYKTNSE  517
alpha-ENaC_Lagenorhynchus_obliquidens      CNMTTYKLSAGYSRWPSVTSQDWVFQMLSLQNNYTVKNKRDGIAKLNIFFKELNYKTNSE  517
alpha-ENaC_Tursiops_truncatus              CNMTTYKLSAGYSRWPSVTSQDWVFQMLSLQNNYTVKNKRDGIAKLNIFFKELNYKTNSE  517
alpha-ENaC_Orcinus_orca                    CKMTTYKLSAGYSRWPSVTSQDWVFQMLSLQNNYTVKNKRDGIAKLNIFFKELNYKTNSE  517
alpha-ENaC_Phocoena_sinus                  CNMTTYKLSAGYSRWPSVTSQDWVFQMLSLQNNYTVKNKRDGVAKLNIFFKELNYKTNSE  517
alpha-ENaC_Neophocaena_asiaeorientalis     CNMTTYKLSAGYSRWPSVTSQDWVFQMLSLQNNYTVKNKRDGVAKLNIFFKELNYKTNSE  517
alpha-ENaC_Monodon_monocerons              CNMTTYKLSAGYSRWPSVTSQDWVFQMLSLQNNYTVKNKRDGVAKLNIFFKELNYKTNSE  517
alpha-ENaC_Delphinapterus_leucas           CNMTTYKLSAGYSRWPSVTSQDWVFQMLSLQNNYTVKNKRDGVAKLNIFFKELNYKTNSE  517
alpha-ENaC_Pontoporia_blainvillei          CNVTTYKLSAGYSRWPSVTSQDWVFQMLSRQNSYTVKNKRDGVAKLNIFFKELNYKTNSE  517
alpha-ENaC_Inia_geoffrensis                CNVITYKLSAGYSRWPSVTSQDWVFQMLSRQNNYTVKNKRDGVAKLNIFFKELNYKTNSE  517
alpha-ENaC_Mesoplodon_bidens               CRVTTYKLSAGYSRWPSVTSQDWVFRMLSRQNNYTIKNKRDGVAKLNIFFKELNYKTNSE  517
alpha-ENaC_Ziphius_cavirostris             CRVTTYKLSAGYSRWPSVTSQDWVFRMLSRQNNYTIKNKREGVAKLNIFFKELNYKTNSE  517
alpha-ENaC_Platanista_gangetica            CSVTTYKLSAGYSRWPSVTSQDWVFQMLSRQNNYTIKNKRDGVAKLNIFFKELNYKTNSE  518
alpha-ENaC_Platanista_minor                CSVTTYKLSAGYSRWPSVTSQDWVFQMLSRQNNYTIKNKRDGVAKLNIFFKELNYKTNSE  518
alpha-ENaC_Kogia_breviceps                 CSVTTYKLSAGYSRWPSVTSQDWVFQMLSRQNSYTVKNKRDGVAKLNIFFKELNYKTNSE  518
alpha-ENaC_Physeter_catodon                CSVTTYKLSAGYSRWPSVTSQDWVFQMLSRQNNYTIKNKRDGVAKLNIFFKELNYKTNSE  518
alpha-ENaC_Balaenoptera_musculus           CSVTTYKLSAGYSRWPSVTSQDWVFKMLSRQNNYTIKNKRDGVAKLNIFFKELNYKTNSE  518
alpha-ENaC_Eubalaena_japonica              CSVTTYKLSAGYSRWPSVTSQDWVFKMLSRQNNYTIKNKRDGVAKLNIFFKELNYKTNSE  518
alpha-ENaC_Hippopotamus_amphibius          CSVTSYKLSAGYSRWPSVTSQDWVFQMLSRQNNYTIKNKRDGVAKLNIFFKELNYKTNSE  518
alpha-ENaC_Tragulus_javanicus              CSLTIYRLSASYSQWPSVTSQDWVFEMLSRQNNYTIKNKRDGVAKLNIFFKELNYKSNTE  519
alpha-ENaC_Antilocapra_americana           CSVTIYKLSASYSQWPSVTSQDWVFQMLSRQNNYTIKNKRDGVAKLNIFFKELNYKSNSE  519
alpha-ENaC_Giraffa_camelopardalis          CSVTIYKLSASYSQWPSVTSQDWVFQMLSRQNNYTIKNKRDGVAKLNIFFKELNYKSNSE  516
alpha-ENaC_Giraffa_tippelskirchi           CSVTIYKLSASYSQWPSVTSQDWVFQMLSRQNNYTIKNKRDGVAKLNIFFKELNYKSNSE  516
alpha-ENaC_Capreolus_pygargus              CSVTIYKLSASYSQWPSMTSQDWVFQMLSRQNNYTIKNKRDGVAKLNIFFKELNYKSNSE  519
alpha-ENaC_Cervus_elaphus                  CSVTIYKLSASYSQWPSVTSQDWVFQMLSRQNNYTIKNKRDGVAKLNIFFKELNYKSNSE  519
alpha-ENaC_Moschus_moschiferus             CSVTIYKLSASYSQWPSVTSQDWVFQMLSRQNNYTIKNKRDGVAKLNIFFKELNYKSNSE  519
alpha-ENaC_Moschus_berezovskii             CSVTIYKLSASYSQWPSVTSQDWVFQMLSRQNNYTIKNKRDGVAKLNIFFKELNYKSNSE  519
alpha-ENaC_Bos_grunniens                   CSVTIYKLSASYSQWPSATSQDWVFQMLSRQNNYTIKNKRDGVAKLNIFFKELNYKSNSE  519
alpha-ENaC_Bos_taurus                      CSVTIYKLSASYSQWPSATSQDWVFQMLSRQNNYTIKNKRDGVAKLNIFFKELNYKSNSE  519
alpha-ENaC_Bubalus_bubalis                 CSVTIYKLSASYSQWPSVTSQDWVFQMLSRQNNYTVKNKRDGVAKLNIFFKELNYKSNSE  519
alpha-ENaC_Nanger_granti                   CSVTIYKLSASYSQWPSMTSQDWVFQMLSRQNNYTIKNKRDGVAKLNIFFKELNYKSNSE  519
alpha-ENaC_Capra_hircus                    CSVTIYKLSASYSQWPSVTSQDWVFQMLSRQNNYTIKNKRDGVAKLNIFFKELNYKSNSE  519
alpha-ENaC_Ovis_aries                      CSVTIYKLSASYSQWPSVTSQDWVFQMLSRQNNYTIKNKRDGVAKLNIFFKELNYKSNSE  519
alpha-ENaC_Ovis_canadensis                 CSVTIYKLSASYSQWPSVTSQDWVFQMLSRQNNYTIKNKRDGVAKLNIFFKELNYKSNSE  519
alpha-ENaC_Hippotragus_niger               CSVTIYKLSASYSQWPSVTSQDWVFEMLSRQNNYTIKNKRDGVAKLNIFFKELNYKSNSE  519
alpha-ENaC_Damaliscus_lunatus              CSVTIYKLSASYSQWPSVTSQDWVFEMLSRQNNYTIKNKRDGVAKLNIFFKELNYKSNSE  519
alpha-ENaC_Sus_scrofa                      CSVTTYKLSAGYSRWPSVTSQDWVFQMLSRQNNYTIKNKRNGVAKVNIFFKELNYKTNSE  519
alpha-ENaC_Vicugna_pacos                   CSVTMYKLSAGYSRWPSVTSQDWVFEMLSRQNNYTIKDKRNGVAKLNIFFKELNYKTNSE  518
alpha-ENaC_Equus_callabus                  CSVTSYQLSAGYSRWPSVTSQDWVFQMLSLQNNYTINNKRNGVAKLNIFFEELNYKTNSE  518
                                           * :  *:***.**:*** ***:***.*** **.**:::**:*:**:****:*****:*:*

                                                TM2
alpha-ENaC_Homo_sapiens                    SPSVTMVTLLSNLGSQWSLWFGSSVLSVVEMAELVFDLLVIMFLMLLRRFRSRYWSPGRG  598
alpha-ENaC_Globicephala_melas              SPSVTMVTLLSNLGSQWSLWFGSSVLSVVEMAELIFDLLAITFFMLLRRFQSQYWSPGRG  577
alpha-ENaC_Lagenorhynchus_obliquidens      SPSVTMVTLLSNLGSQWSLWFGSSVLSVVEMAELIFDLLAITFFMLLRRFQSQYWSPGRG  577
alpha-ENaC_Tursiops_truncatus              SPSVTMVTLLSNLGSQWSLWFGSSVLSVVEMAELIFDLLAITFFMLLRRFQSQYWSPGRG  577
alpha-ENaC_Orcinus_orca                    SPSVTMVTLLSNLGSQWSLWFGSSVLSVVEMAELIFDLLAITFFMLLRRFQSQYWSPGRG  577
alpha-ENaC_Phocoena_sinus                  SPSVTMVTLLSNLGSQWSLWFGSSVLSVVEMAELIFDLLAITFLMLLRRFQSQYWSPGRG  577
alpha-ENaC_Neophocaena_asiaeorientalis     SPSVTMVTLLSNLGSQWSLWFGSSVLSVVEMAELIFDLLAITFLMLLRRFQSQYWSPGRG  577
alpha-ENaC_Monodon_monocerons              SPSVTMVTLLSNLGSQWSLWFGSSVLSVVEMAELIFDLLAITFLMLLRRFQSQYWSPGRG  577
alpha-ENaC_Delphinapterus_leucas           SPSVTMVTLLSNLGSQWSLWFGSSVLSVVEVAELIFDLLAITFLMLLRRFQSQYWSPGRG  577
alpha-ENaC_Pontoporia_blainvillei          SPSVTMVTLLSNLGSQWSLWFGSSVLSVVEMAELVFDLLAITFLMLLRRFQSQYWSPGRG  577
alpha-ENaC_Inia_geoffrensis                SPSVTMVTLLSKLGSQWSLWFGSSVLSVVEMAELVFDLLAITFLMLLRRFQSQYWSPGRG  577
alpha-ENaC_Mesoplodon_bidens               SPSVTMVTLLSNLGSQWSLWFGSSVLSVVEMAELIFDLLAITFLMLLRRFQSQYWSPGRG  577
alpha-ENaC_Ziphius_cavirostris             SASITMVTLLSNLGSQWSLWFGSSVLSVVEMAELIFDLLAITFLMLLRRLQSQYWSPGRG  577
alpha-ENaC_Platanista_gangetica            SPSVTMVTLLSNLGSQWSLWFGSSVLSVVEMAELIFDLLAITFLMLLRRFQSQYWSPGRG  578
alpha-ENaC_Platanista_minor                SPSVTMVTLLSNLGSQWSLWFGSSVLSVVEMAELIFDLLAITFLMLLRRFQSQYWSPGRG  578
```

```
alpha-ENaC_Kogia_breviceps              SPSVTMVTLLSNLGSQWSLWFGSSVLSVVEMAELIFDLLAITFLMLLRRFQSRYWSPGRG  578
alpha-ENaC_Physeter_catodon             SPSVTMVTLLSNLGSQWSLWFGSSVLSVVEMAELIFDLLAITFLMLLRRVQSRYWSPGRG  578
alpha-ENaC_Balaenoptera_musculus        SPSVTTVTLLSNLGSQWSLWFGSSVLSVVEVAELIFDLLAITFLMLLRRFQSQYWSPGRG  578
alpha-ENaC_Eubalaena_japonica           SPSVTTVTLLSNLGSQWSLWFGSSVLSVVEVAELIFDLLAITFLMLLRRFQSQYWSPGRG  578
alpha-ENaC_Hippopotamus_amphibius       SPSVTMVTLLSNLGSQWSLWFGSSVLSVVEMAELLFDLLVITFLMLLRRCRSRYWSPGRG  578
alpha-ENaC_Tragulus_javanicus           SPSVKMATLLSNLGSQWSLWFGSSVLSVVEMAELIFDLLVITILMLLRRLRSRYWSPGRG  579
alpha-ENaC_Antilocapra_americana        SPSVTMVTLLSNLGSQWSLWFGSSVLSVVEMAELIFDLLVITFLMLLRRFQSRYWSPGRG  579
alpha-ENaC_Giraffa_camelopardalis       SPLVTMVTLLSKLGSQWSLWFGSSVLSVVEMAELIFDLLVITFLMLLRRFRSRYWSPSRG  576
alpha-ENaC_Giraffa_tippelskirchi        SPLVTMVTLLSKLGSQWSLWFGSSVLSVVEMAELIFDLLVITFLMLLRRFRSRYWSPSRG  576
alpha-ENaC_Capreolus_pygargus           SPSVTMVTLLSNLGSQWSLWFGSSVLSVVEMAELIFDLLVITFLMLLRRFRSRYWSPGRG  579
alpha-ENaC_Cervus_elaphus               SPSVTMVTLLSNLGSQWSLWFGSSVLSVVEMAELIFDLLVITFLMLLRRFRSRYWSPGRG  579
alpha-ENaC_Moschus_moschiferus          SPSVTMVTLLSNLGSQWSLWFGSSVLSVVEMAELIFDLLVITFLMLLRRFRSRYWSPGRG  579
alpha-ENaC_Moschus_berezovskii          SPSVTMVTLLSNLGSQWSLWFGSSVLSVVEMAELIFDLLAITFLMLLRRFRSRYWSPGRG  579
alpha-ENaC_Bos_grunniens                SPSVTMVTLLSNLGSQWSLWFGSSVLSVVEMAELIIDLLVITFLMLLRRFRSRYWSPGRG  579
alpha-ENaC_Bos_taurus                   SPSVTMVTLLSNLGSQWSLWFGSSVLSVVEMAELIIDLLVITFLMLLRRFRSRYWSPGRG  579
alpha-ENaC_Bubalus_bubalis              SPSVTMVTLLSNLGSQWSLWFGSSVLSVVEMAELIIDLLVITFLMLLRRFRSRYWSPGRG  579
alpha-ENaC_Nanger_granti                SPSVTMVTLLSNLGSQWSLWFGSSVLSVVEMAELIFDLLVITFLMLLRRFRSRYWSPGRG  579
alpha-ENaC_Capra_hircus                 SPSVTMVTLLSNLGSQWSLWFGSSVLSVVEMAEFIFDLLVITFLMLLRRFRSRYWSPGRG  579
alpha-ENaC_Ovis_aries                   SPSVTMVTLLSNLGSQWSLWFGSSVLSVVEMAELIFDLLVITFLMLLRRFRSRYWSPGRG  579
alpha-ENaC_Ovis_canadensis              SPSVTMVTLLSNLGSQWSLWFGSSVLSVVEMAELIFDLLVITFLMLLRRFRSRYWSPGRG  579
alpha-ENaC_Hippotragus_niger            SPSVTMVTLLSNLGSQWSLWFGSSVLSVVEMAELIFDLLVITFLMLLRRFRSRYWSPGRG  579
alpha-ENaC_Damaliscus_lunatus           SPSVTMVTLLSNLGSQWSLWFGSSVLSVVEMAELIFDLLVITFLMLLRRFRSRYWSPGRG  579
alpha-ENaC_Sus_scrofa                   SPSVTMVTLLSNLGSQWSLWFGSSVLSVVEMAELIFDLLVITFLMLLRRFRSRYWSPGRG  579
alpha-ENaC_Vicugna_pacos                SPSVTMVTLLSNLGGQWSLWFGSSVLSVVEMAELVFDLLAITFLMLIRRVRSRYWSPGRG  578
alpha-ENaC_Equus_callabus               SPSVTMVTLLSNLGSQWSLWFGSSVLSVVEMAELIFDLLVITFLMLLRRIRSRYWSPGRG  578
                                        *  :. .****:**.***************:**:::***.* ::**:** :*:****.**

alpha-ENaC_Homo_sapiens                 GRGAQEVASTLASSPPSHFCPHPMSLSLSQPGPAPSPALTAPPPAYATLGPRPSPGGSAG  658
alpha-ENaC_Globicephala_melas           GRGAQEVASTPPSSLPSRFCPHPASPSSSAAGPATSLALSAPPPAYATLGPRLAPSGSTE  637
alpha-ENaC_Lagenorhynchus_obliquidens   GRGAQEVASTPPSSLPSRFCPHPASPSSSAAGPATSLALSAPPPAYATLGPRLAPSGSTE  637
alpha-ENaC_Tursiops_truncatus           GRGAQEVASTPPSSLPSRFCPHPASPSSSAAGPATSLALSAPPPAYATLGPRLAPSGSTE  637
alpha-ENaC_Orcinus_orca                 GRGAQEVASTPPSSLPSRFCPHPASPSSSAVGPATSLALSAPPPAYATLGPRLAPSGSTE  637
alpha-ENaC_Phocoena_sinus               GRGAQEVASTPPSSLPSRFCPHPASPPASLADPATSLALSAPPPAYATLGPRLAPSGSME  637
alpha-ENaC_Neophocaena_asiaeorientalis  GRGAQEVASTPPSSLPSRFCPHPASPPASLADPATSLALSAPPPAYATLGPRLAPSGSME  637
alpha-ENaC_Monodon_monocerons           GRGAQEVASTPASSLPSRFCPHPASPSSSLAGPATSLALSAPPPAYATLGPRLAPSGSME  637
alpha-ENaC_Delphinapterus_leucas        GRGAQEVASTPASSLPSRFCPHPASPSSSLGGPATSLALSAPPPAYATLGPRLAPSGSME  637
alpha-ENaC_Pontoporia_blainvillei       GRGAQEVASTPASSLPSRFCPHPATPSSSPAGPATSLALSAPPPAYATLGPHLAPSGSTE  637
alpha-ENaC_Inia_geoffrensis             GRGAQEVASTPASSLPSRFCPHPTSPSSSPAGPATSLALSAPPPAYATLGPRLAPSGSTE  637
alpha-ENaC_Mesoplodon_bidens            GRGSQEVASTPASSLPSRFCPHPASPSSSPPGPATSLALSAPPPAYATLGPRLALSGSTE  637
alpha-ENaC_Ziphius_cavirostris          GRGSQEVASTPASSLPSRFCPHPASPSSSPPGPATTLALSAPPPAYATLGPRLALSGSTE  637
alpha-ENaC_Platanista_gangetica         GRGAQEVASTPASSLPSRFCPHPASPSSSPPGPATSLALSAPPPAYATLGPRLAPSGSTE  638
alpha-ENaC_Platanista_minor             GRGAQEVASTPASSLPSRFCPHPASPSSSPPGPATSLALSAPPPAYATLGPRLAPSGSTE  638
alpha-ENaC_Kogia_breviceps              GSGAREVASMPTSAFPSRFCPHPASPSSFPPGPATSLALSAPPPAYATLGPRWTPSGSTE  638
alpha-ENaC_Physeter_catodon             GRGAQEVASTPTSALPSRFCPHPASPSSFPPGPAASLALSAPPPAYATLGPRRAPSGSTE  638
alpha-ENaC_Balaenoptera_musculus        RRGAQEVASTPASSLPSRFCPHPASPSSSPPGPATSLALSAPPPAYATLGPRLAPSGSTE  638
alpha-ENaC_Eubalaena_japonica           RRGAQEVASTPASSLPSRFCPHPASPSSSPLGPATSLALSAPPPAYATLGPRLAPSGSTE  638
alpha-ENaC_Hippopotamus_amphibius       RGGAQEVASTPASSLPSRFCPHPASPSSSLPGPAPSPALSAPPPAYATLGPHPAPSGSTE  638
alpha-ENaC_Tragulus_javanicus           GRGAQEVASTPASSPPSSVCPDPASSSSP-GPATAPALSAPPPAYNTLGPHPAPLGLAE  638
alpha-ENaC_Antilocapra_americana        GRGTQEVASTPAASLPSSFCPHPASSSSSPPEPAISPALSAPPPAYATLGPHPGPSGLAE  639
alpha-ENaC_Giraffa_camelopardalis       GRGAQEVASTPASSLPSSFCPHPASSSSSLPHPAISPALSAPPPAYATLGPHPAPSGLVE  636
alpha-ENaC_Giraffa_tippelskirchi        GRGAQEVASTPASSLPSSFCPHPASSSSSLPHPAISPALSAPPPAYATLGPHPAPSGLVE  636
alpha-ENaC_Capreolus_pygargus           GRGSQEVASTPAASLPSSFCPHLASSSSSPPDPAISPALSAPPPAYATLGPHPVPSGLAE  639
alpha-ENaC_Cervus_elaphus               GRGTQEVASTPAASLPSSFCPHPASSSSSPPDPAISPALSAPPPAYATLGPHPAPSGLAE  639
alpha-ENaC_Moschus_moschiferus          GRGTQELASTPAASLPSSFCPHPASSSSSPPDPAISPALSAPPPAYATLGPHPAPSGLAE  639
alpha-ENaC_Moschus_berezovskii          GRGTQEVASTPAASLPSSFCPHPASSSSSPPDPAISPALSAPPPAYATLGPHPAPSGLAE  639
alpha-ENaC_Bos_grunniens                GKGTQEVASTPATSLPSSFCPHPAFSSSYPPDPAISPALSAPPPAYATLGPHPAPSGLAE  639
alpha-ENaC_Bos_taurus                   GKGTQEVASTPAASLPSSFCPHPAFFSSSPPDPAISPALSAPPPAYATLGPHPAPSGLAE  639
alpha-ENaC_Bubalus_bubalis              GKGTQEVASTLAASLPSSFCPHPASSSSSLPDPAISPALSAPPPAYATLGPHPAPSGLAE  639
alpha-ENaC_Nanger_granti                GRGTQEVASTPATSLPSSFCPHQASSSSSPLDPAVSPALSAPPPAYATLGPHPAPSGLAE  639
alpha-ENaC_Capra_hircus                 GRGTQEVASTPATSLPSSSCPYQASSSSSPPDPAISPALSAPPPAYSTLGPHPAPSGLAE  639
alpha-ENaC_Ovis_aries                   GRGTQEVASTPATSLPSSSCPYQASSSSFPPDPAISPALSAPPPAYSTLGPHPAPTGLAE  639
alpha-ENaC_Ovis_canadensis              GRGTQEVASTPATSLPSSSCPYQASSSSFPPDPAISLALSAPPPAYSTLGPHPAPSGLAE  639
alpha-ENaC_Hippotragus_niger            GRGTQEVASTPATSLPSSFCPYQASSSSSPPDPAISLALSAPPPAYATLGPHPAPSGLAE  639
alpha-ENaC_Damaliscus_lunatus           GRGTQEVASTPATSLPSSFCPYQASSSSSSPDPAISLALSAPPPAYATLGPHPAPSGLAE  639
alpha-ENaC_Sus_scrofa                   GRGAQEVASTPPSSLPSRFCPHPASPSSSPPGPAANPALSAPPPAYATLGPCPAPSGSAA  639
alpha-ENaC_Vicugna_pacos                ARAGQEVASASAPPLPSHFCPHPTSPSSSLPGPAPAPALAAPPPAYATLGPPSSLSGSAA  638
alpha-ENaC_Equus_callabus               RRGAREVASTPASSLPAHFCPNPASPSSSPPGPAASPALTAPPPAYATLGPCPSLASSAG  638
                                          . :*:**      *:  **            **   **:****** ****      .

alpha-ENaC_Homo_sapiens                 ASSSTCPLGGP    669
alpha-ENaC_Globicephala_melas           ASSSAHSPGEP    648
alpha-ENaC_Lagenorhynchus_obliquidens   ASSSAHTPGEP    648
alpha-ENaC_Tursiops_truncatus           ASSSAHTPGEP    648
alpha-ENaC_Orcinus_orca                 ASSSAHTPGEP    648
alpha-ENaC_Phocoena_sinus               ASSSAHTPGEP    648
alpha-ENaC_Neophocaena_asiaeorientalis  ASSSAHTPGEP    648
alpha-ENaC_Monodon_monocerons           ASSSAHTPGEP    648
alpha-ENaC_Delphinapterus_leucas        ASSSAHTPGEP    648
alpha-ENaC_Pontoporia_blainvillei       ASSSAHTPGEP    648
alpha-ENaC_Inia_geoffrensis             ASSSAHTPGEP    648
alpha-ENaC_Mesoplodon_bidens            ASSSAHTPGEP    648
alpha-ENaC_Ziphius_cavirostris          ASSSAHTPGEP    648
alpha-ENaC_Platanista_gangetica         ASSSAHTPGKP    649
alpha-ENaC_Platanista_minor             ASSSAHTPGKP    649
```

| | | |
|---|---|---|
| alpha-ENaC_Kogia_breviceps | ASASAHTAGEP | 649 |
| alpha-ENaC_Physeter_catodon | ASSSAHTLREP | 649 |
| alpha-ENaC_Balaenoptera_musculus | ASSSAHTPGEP | 649 |
| alpha-ENaC_Eubalaena_japonica | ASSSAHAPGEP | 649 |
| alpha-ENaC_Hippopotamus_amphibius | ASSSAHTPGEP | 649 |
| alpha-ENaC_Tragulus_javanicus | ANLSAHAPGEP | 649 |
| alpha-ENaC_Antilocapra_americana | ASASAHAPGEP | 650 |
| alpha-ENaC_Giraffa_camelopardalis | ASASAHTPGEP | 647 |
| alpha-ENaC_Giraffa_tippelskirchi | ASASAHTPGEP | 647 |
| alpha-ENaC_Capreolus_pygargus | ASASAHALGEP | 650 |
| alpha-ENaC_Cervus_elaphus | ASASAHTSGEP | 650 |
| alpha-ENaC_Moschus_moschiferus | ASASAHAPGEP | 650 |
| alpha-ENaC_Moschus_berezovskii | ASASAHAPGEP | 650 |
| alpha-ENaC_Bos_grunniens | ASTSAHAPGEP | 650 |
| alpha-ENaC_Bos_taurus | ASTSAHAPGEP | 650 |
| alpha-ENaC_Bubalus_bubalis | ASTSAHAPGEP | 650 |
| alpha-ENaC_Nanger_granti | ASIFAHAQGEP | 650 |
| alpha-ENaC_Capra_hircus | ASASAHAPGEP | 650 |
| alpha-ENaC_Ovis_aries | ASASAHAPGEP | 650 |
| alpha-ENaC_Ovis_canadensis | ASASAHGPGEP | 650 |
| alpha-ENaC_Hippotragus_niger | ASAP------- | 643 |
| alpha-ENaC_Damaliscus_lunatus | ASASAHAPGEP | 650 |
| alpha-ENaC_Sus_scrofa | AGSSAHPLGEP | 650 |
| alpha-ENaC_Vicugna_pacos | ASNPAYTPEEV | 649 |
| alpha-ENaC_Equus_callabus | ASSSACTPGEP | 649 |
| | *. | |

**β-ENaC (*SCNN1B*) amino acid sequence alignment** created with Clustal Omega (v.1.2.4, accessed 24.03.2024). Key structural motifs are highlighted in the human ENaC subunit based on the Cryo-EM derived structure (Noreng et al. 2018). Transmembrane domains (TM1/TM2) are highlighted in yellow. An N-Terminal HG-motif affecting ENaC open probability is marked in magenta. Cysteines involved in secondary tertiary structure formation are indicated in red. Residues putatively forming the selectivity filter within TM2 are shown in red font. The C-terminal PPPxY motif regulating membrane abundance is shown in **gray**.

```
                                                                                        TM1
beta-ENaC_Homo_sapiens                                    -MHVKKYLLKGLHRLQKGPGYTYKELLVWYCDNTNTHGPKRIICEGPKKKAMWFLLTLLF    59
beta-ENaC_Globicephala_melas                              -MHIKKYLLKCLHRLQKGPGYTYKELLVWYCDNTNTHGPKRIICEGPKKKAMWFVLTLLF    59
beta-ENaC_Lagenorhynchus_obliquidens                      -MHIKKYLLKCLHRLQKGPGYTYKELLVWYCDNTNTHGPKRIICEGPKKKAMWFVLTLLF    59
beta-ENaC_Tursiops_truncatus                              -MHIKKYLLKCLHRLQKGPGYTYKELLVWYCDNTNTHGPKRIICEGPKKKAMWFVLTLLF    59
beta-ENaC_Orcinus_orca                                    -MHIKKYLLKCLHRLQKGPGYTYKELLVWYCDNTNTHGPKRIICEGPKKKAMWFVLTLLF    59
beta-ENaC_Phocoena_sinus                                  -MHIKKYLLKCLHRLQKGPGYTYKELLVWYCDNTNTHGPKRIICEGPKKKAMWFVLTLLF    59
beta-ENaC_Neophocaena_asiaeorientalis                     -MHIKKYLLKCLHRLQKGPGYTYKELLVWYCDNTNTHGPKRIICEGPKKKAMWFVLTLLF    59
beta-ENaC_Monodon_monocerons                              MMHIKKYLLKCLHRLQKGPGYTYKELLVWYCDNTNTHGPKRIICEGPKKKAMWFVLTLLF    60
beta-ENaC_Delphinapterus_leucas                           -MHIKKYLLKCLHRLQKGPGYTYKELLVWYCDNTNTHGPKRIICEGPKKKAMWFVLTLLF    59
beta-ENaC_Pontoporia_blainvillei                          -MHIKKYLLKCLHRLQKGPGYTYKELLVWYCDNTNTHGPKRIICEGPKKKAMWFVLTLLF    59
beta-ENaC_Inia_geoffrensis                                -MHIKKYLLKCLHRLQKGPGYTYKELLVWYCDNTNTHGPKRIICEGPKKKAMWFVLTLLF    59
beta-ENaC_Lipotes_vexillifer                              -MHIKRYLLKCLHRLQKGPGYTYKELLVWYCDNTNTHGPKRIICEGPKKKAMWFVLTLLF    59
beta-ENaC_Hyperoodon_ampullatus                           -MHIKKYLLKCLHRLQKGPGYTYKELLVWYCDNTNTHGPKRIICEGPKKKAMWFVLTLLF    59
beta-ENaC_Mesoplodon_bidens                               -MHIKKYLLKCLHRLQKGPGYTYKELLVWYCDNTNTHGPKRIICEGPKKKAMWFVLTLLF    59
beta-ENaC_Ziphius_cavirostris                             -MHIKKYLLKCLHRLQKGPGYTYKELLVWYCDNTNTHGPKRIICEGPKKKAMWFVLTLLF    59
beta-ENaC_Platanista_gangetica                            -MHIKKYLLKCLHRLQKGPGYTYKELLVWYCDNTNTHGPKRIICEGPKKKAMWFVLTLLF    59
beta-ENaC_Platanista_minor                                -MHIKKYLLKCLHRLQKGPGYTYKELLVWYCDNTNTHGPKRIICEGPKKKAMWFVLTLLF    59
beta-ENaC_Kogia_breviceps                                 -MHIKKYLLKCLHRLQKGPGYTYKELLVWYCDNTNTHGPKRIICEGPKKKAMWFVLTLLF    59
beta-ENaC_Physeter_catodon                                -MHIKKYLLKCLHRLQKGPGYTYKELLVWYCDNTNTHGPKRIICEGPKKKAMWFVLTLLF    59
beta-ENaC_Balaenoptera_musculus                           -MHIKKYLLKCLHRLQKGPGYTYKELLVWYCDNTNTHGPKRIICEGPKKKAMWFVLTLLF    59
beta-ENaC_Balaenoptera_acutorostrata_scammoni             -MHVKKYLLKCLHRLQKGPGYTYKELLVWYCDNTNTHGPKRIICEGPKKKAMWFVLTLLF    59
beta-ENaC_Eubalaena_japonica                              -MHIKKYLLKCLHRMQKGPGYTYKELLVWYCDNTNTHGPKRIICEGPKKKAMWFVLTLLF    59
beta-ENaC_Hippopotamus_amphibius                          MVPIKKYLLKCLHRLQKGPGYTYKELLVWYCDNTNTHGPKRIICEGPKKKAMWFVLTLLF    60
beta-ENaC_Tragulus_javanicus                              -MHVKKYLLKGLHRLQKGPGYTYKELLVWYCDNTNTHGPKRIICEGPKKKAMWFVLTLLF    59
beta-ENaC_Tragulus_kanchil                                -MHVKKYLLKGLHRLQKGPGYTYKELLVWYCDNTNTHGPKRIICEGPKKKAMWFVLTLLF    59
beta-ENaC_Antilocapra_americana                           -MHVKKYLLKGLHRLQKGPGYTYKELLVWYCDNTNTHGPKRIICEGPKKKAMWFVLTLLF    59
beta-ENaC_Giraffa_camelopardalis                          -MHVKKYLLKGLHRLQKGPGYTYKELLVWYCDNTNTHGPKRIICEGPKKKAMWFMLTLLF    59
beta-ENaC_Giraffa_tippelskirchi                           -MHVKKYLLKGLHRLQKGPGYTYKELLVWYCDNTNTHGPKRIICEGPKKKAMWFMLTLLF    59
beta-ENaC_Capreolus_pygargus                              MMHVKKYLLKGLHRLQKGPGYTYKELLVWYCDNTNTHGPKRIICEGPKKKAMWFVLTLLF    60
beta-ENaC_Cervus_elaphus                                  -MHVKKYLLKGLHRLQKGPGYTYKELLVWYCDNTNTHGPKRIICEGPKKKAMWFVLTLLF    59
beta-ENaC_Moschus_moschiferus                             -MHVKKYLLKGLHRLQKGPGYTYKELLVWYCDNTNTHGPKRIICEGPKKKAMWFVLTLLF    59
beta-ENaC_Moschus_berezovskii                             -MHVKKYLLKGLHRLQKGPGYTYKELLVWYCDNTNTHGPKRIICEGPKKKAMWFVLTLLF    59
beta-ENaC_Bos_grunniens                                   -MHVKKYLLKGLHRLQKGPGYTYKELLVWYCDNTNTHGPKRIICEGPKKKAMWFVLTLLF    59
beta-ENaC_Bos_taurus                                      -MHVKKYLLKGLHRLQKGPGYTYKELLVWYCDNTNTHGPKRIICEGPKKKAMWFVLTLLF    59
beta-ENaC_Bubalus_bubalis                                 -MHVKKYLLKGLHRLQKGPGYTYKELLVWYCDNTNTHGPKRIICEGPKKKAMWFVLTLLF    59
beta-ENaC_Nanger_granti                                   -MHVKKYLLKGLHRLQKGPGYTYKELLVWYCDNTNTHGPKRIICEGPKKKAMWFVLTLLF    59
beta-ENaC_Capra_hircus                                    -MHLKKYLLKGLHRLQKGPGYTYKELLVWYCDNTNTHGPKRIICEGPKKKAMWFVLTLLF    59
beta-ENaC_Ovis_aries                                      -MHLKKYLLKGLHRLQKGPGYTYKELLVWYCDNTNTHGPKRIICEGPKKKAMWFVLTLLF    59
beta-ENaC_Ovis_canadensis                                 -MHLKKYLLKGLHRLQKGPGYTYKELLVWYCDNTNTHGPKRIICEGPKKKAMWFVLTLLF    59
beta-ENaC_Hippotragus_niger                               MMHVKKYLLKGLHRLQKGPGYTYKELLVWYCGNTNTHGPKRIICEGPKKKAMWFVLTLLF    60
beta-ENaC_Sus_scrofa                                      -MHVKKYLLKCLHRLQKGPGYTYKELLVWYCDNTNTHGPKRIIREGPKKKAMWFVLTLLF    59
beta-ENaC_Vicugna_pacos                                   -MHVKKYLLKCLHRLQKGPGYTYKELLVWYCDNTNTHGPKRIICEGPKKKAMWFVITLLF    59
beta-ENaC_Equus_callabus                                  -MHVKKYLLKGLHRLQKGPGYTYKELLVWYCDNTNTHGPKRIICEGPKKKAMWFLLTLLF    59
                                                           : :*:**** ***:****************.*********** **********::****

beta-ENaC_Homo_sapiens                                    AALVCWQWGIFIRTYLSWEVSVSLSVGFKTMDFPAVTICNASPFKYSKIKHLLKDLDELM    119
beta-ENaC_Globicephala_melas                              TSLVCWQWGVFIRTYLNWEVSVSLSIGFKAMDFPAVTVCNTSPFQYSKVKHLLKDLDELM    119
beta-ENaC_Lagenorhynchus_obliquidens                      TSLVCWQWGIFIRTYLNWEVSVSLSIGFKAMDFPAVTVCNTSPFQYSKVKHLLKDLDELM    119
beta-ENaC_Tursiops_truncatus                              TSLVCWQWGVFIRTYLNWEVSVSLSIGFKAMDFPAVTVCNTSPFQYSKVKHLLKDLDELM    119
beta-ENaC_Orcinus_orca                                    TSLVCWQWGVFIRTYLNWEVSVSLSIGFKAMDFPAVTVCNTSPFQYSKVKHLLKDLDELM    119
beta-ENaC_Phocoena_sinus                                  TSLVCWQWGVFIRTYLSWEVSVSLSIGFKAMDFPAVTICNTSPFQYSKVKHLLKDLDELM    119
beta-ENaC_Neophocaena_asiaeorientalis                     TSLVCWQWGVFIRTYLSWEVSVSLSIGFKAMDFPAVTICNTSPFQYSKVKHLLKDLDELM    119
beta-ENaC_Monodon_monocerons                              TSLVCWQWGVFIRTYLSWEVSVSLSIGFKAMDFPAVTICNTSPFQYSKVKHLLKDLDELM    120
beta-ENaC_Delphinapterus_leucas                           TSLVCWQWGVFIRTYLSWEVSVSLSIGFKAMDFPAVTICNTSPFQYSKVKHLLKDLDELM    119
beta-ENaC_Pontoporia_blainvillei                          TSLVCWQWGVFIRTYLSWEVSVSLSIGFKAMDFPAVTICNTSPFQYSKVKHLLKDLDELM    119
beta-ENaC_Inia_geoffrensis                                TSLVCWQWGVFIRTYLSWEVSVSLSIGFKAMDFPAVTICNTSPFQYSKAKHLLKDLDELM    119
beta-ENaC_Lipotes_vexillifer                              TSLVCWQWGVFIRTYLSWEVSVSLSIGFKAMDFPAVTICNTSPFQYSKVKHLLKDLDELM    119
beta-ENaC_Hyperoodon_ampullatus                           ASLVCWQWGVFIRTYLSWEVSVSLSIGFKAMDFPAVTVCNTSPFRYSKVKHLLKDLDELM    119
beta-ENaC_Mesoplodon_bidens                               ASLVCWQWGVFIRTYLSWEVSVSLSIGFKAMDFPAVTVCNTSPFRYSKVKHLLKDLDELM    119
beta-ENaC_Ziphius_cavirostris                             ASLVCWQWGVFIRTYLSWEVSVSLSIGFKAMDFPAVTVCNTSPFRYSKVKHLLKDLDELM    119
beta-ENaC_Platanista_gangetica                            ASLVCWQWGVFIRTYLSWEVSVSLSVGFKAMDFPAVTICNTSPFQYSKVKHLLKDLDELM    119
beta-ENaC_Platanista_minor                                ASLVCWQWGVFIRTYLSWEVSVSLSVGFKAMDFPAVTICNTSPFQYSKVKHLLKDLDELM    119
beta-ENaC_Kogia_breviceps                                 ASLVCWQWGVFIRTYLSWEVSVSLSIGFKAMDFPAVTICNTNPFQYSKVKHLLKDLDELM    119
beta-ENaC_Physeter_catodon                                TSLVCWQWGVFIRTYLSWEVSVSLSIGFKAMDFPAVTICNTSPFQYSKVKHLLKDLDELM    119
beta-ENaC_Balaenoptera_musculus                           TSLVCWQWGVFIRTYLSWEVSVSLSIGFKAMDFPAVTICNTSPFQYSKVKHLLKDLDELM    119
beta-ENaC_Balaenoptera_acutorostrata_scammoni             TSLVCWQWGVFIRTYLSWEVSVSLSIGFKAMDFPAVTICNTSPFQYSKIKHLLKDLDELM    119
beta-ENaC_Eubalaena_japonica                              TSLVCWQWGVFIRTYLSWEVSVSLSIGFKAMDFPAVTICNTSPFQYSKVKHLLKDLDELM    119
beta-ENaC_Hippopotamus_amphibius                          TSLVCWQWGVFIKTYLSWEVSVSLSVGFKAMDFPAVTICNTSPFQYSKVKHLLKDLDELM    120
beta-ENaC_Tragulus_javanicus                              ASIVFWQWGVFIKNYLNWDVSVSLSIGFKTMDFPAVTICNTSPFQYPKVQHLLKDLDELM    119
beta-ENaC_Tragulus_kanchil                                ASIVFWQWGVFIKNYLNWDVSVSLSIGFKTMDFPAVTICNTSPFQYPKVQHLLKDLDELM    119
beta-ENaC_Antilocapra_americana                           ASLVCWQWGLFIKTYLNWEVSVSLSIGFKTMDFPAVTICNASPFQYSKVQHLLKDLDELM    119
beta-ENaC_Giraffa_camelopardalis                          ASLVCWQWGLFIKTYLNWEVSVSLSIGFKTMDFPAVTICNASPFQYSKVQHLLKDLDELM    119
beta-ENaC_Giraffa_tippelskirchi                           ASLVCWQWGLFIKTYLNWEVSVSLSIGFKTMDFPAVTICNASPFQYSKVQHLLKDLDELM    119
beta-ENaC_Capreolus_pygargus                              TSLVCWQWGLFIKTYLNWEVSVSLSIGFKTMNFPAVTICNASPFQYSKVQHLLKDLDELM    120
beta-ENaC_Cervus_elaphus                                  TSLVCWQWGLFIKTYLNWEVSVSLSIGFKTMDFPAVTICNASPFQYSKVQHLLKDLDELM    119
beta-ENaC_Moschus_moschiferus                             TSLVCWQWGLFIKTYLNWEVSVSLSIGFKTMDFPAVTICNASPFQYSKVQHLLKDLDELM    119
beta-ENaC_Moschus_berezovskii                             TSLVCWQWGLFIKTYLNWEVSVSLSIGFKTMDFPAVTICNASPFQYSKVQHLLKDLDELM    119
```

```
beta-ENaC_Bos_grunniens                                 TSLVCWQWGLFIKTYLNWEVSVSLSIGFKTMDFPAVTICNASPFQYSKVQHLLKDLDELM    119
beta-ENaC_Bos_taurus                                    TSLVCWQWGLFIKTYLNWEVSVSLSIGFKTMDFPAVTICNASPFQYSKVQHLLKDLDELM    119
beta-ENaC_Bubalus_bubalis                               TSLVCWQWGLFIKTYLNWEVSVSLSIGFKTMDFPAVTICNASPFQYSKVQHLLKDLDELM    119
beta-ENaC_Nanger_granti                                 TSLVCWQWGLFIKTYLNWEVSVSLSIGFKTMDFPAVTICNASPFQYSKVQHLLKDLDALM    119
beta-ENaC_Capra_hircus                                  TSLVCWQWGLFIKTYLNWEVSVSLSIGFKTMDFPAVTICNASPFQYSKVQHLLKDLDALM    119
beta-ENaC_Ovis_aries                                    TSLVCWQWGLFIKTYLNWEVSVSLSIGFKTMDFPAVTICNASPFQYSKVQHLLKDLDALM    119
beta-ENaC_Ovis_canadensis                               TSLVCWQWGLFIKTYLNWEVSVSLSIGFKTMDFPAVTICNASPFQYSKVQHLLKDLDALM    119
beta-ENaC_Hippotragus_niger                             TSLVCWQWGLFIKTYLNWEVSVSLSIGFKTMDFPAVTICNASPFQYSKVQHLLKDLDALM    120
beta-ENaC_Sus_scrofa                                    TSLVCWQWGVFIRTYLSWEVSVSLSMGFKTMDFPAVTICNASPFQYSKVQHLLKDLDELM    119
beta-ENaC_Vicugna_pacos                                 AAIVCWQWGVFIRTYLSWEVSVSLSIGFKTMDFPAVTICNASPFRYSKVKHLLKDLDELM    119
beta-ENaC_Equus_callabus                                AALVCWQWGIFIRTYLSWEVSVSLSVGFKTMDFPAVTICNASPFQYSKVRHLLKDLDELM    119
                                                        ::* ****:**:.**.*:******:***:*:*****:**:.**:* * :******* **

beta-ENaC_Homo_sapiens                                  EAVLERILAPELSHANATRNLNFSIWNHTPLVLIDERNPHHPMVLDLFGDNHNGLTSSS-    178
beta-ENaC_Globicephala_melas                            EAVLARILGPELSHVNATRALNLTIWNHVPLVFIDEQNPRHPVVLDLFEDNYNGSASSTP    179
beta-ENaC_Lagenorhynchus_obliquidens                    EAVLARILGPELSHVNATRALNLTIWNHVPLVFIDEQNPRHPVVLDLFEDNYNGSASSSP    179
beta-ENaC_Tursiops_truncatus                            EAVLARILGPELSHVNATRALNLTIWNHVPLVFIDEQNPHHPVVLDLFEDNYNGSAGSTP    179
beta-ENaC_Orcinus_orca                                  EAVLARILGPELSHVNATRALNLTIWNHVPLVFIDEQNPRHPVVLDLFEDNYNGSASSTP    179
beta-ENaC_Phocoena_sinus                                EAVLERILGPELSHVNATRALNLTIWNHVPLVFIDEQNPHHPVVLDLFEDNYNGSASSTP    179
beta-ENaC_Neophocaena_asiaeorientalis                   EAVLERILGPELSHVNTTRALNLTIWNHVPLVFIDEQNPHHPVVLDLFEDNYNGSASSTP    179
beta-ENaC_Monodon_monocerons                            EAVLERILGPELSHVNTTRALNLTIWNHVPLVFINEQNPHHPVVLDLFEDNYSGSASSTP    180
beta-ENaC_Delphinapterus_leucas                         EAVLERILGPELSHVNATRALNLTIWNHVPLVFIDEQNPHHPVVLDLFEDNYNGSASSTP    179
beta-ENaC_Pontoporia_blainvillei                        EAVLERILGPELSPVNATRALNLTIWNHVPLVFINEQNPHHPVVLDLFEDSYNGSASSTP    179
beta-ENaC_Inia_geoffrensis                              EAVLERILGPELSPVNATRALNLTIWNHMPLVFINEQNPHHPVVLDLFEDSYNGSASSTP    179
beta-ENaC_Lipotes_vexillifer                            EAVLERILGPELSHVNATRALNLTLWNHMPLVFINEQNPHHPVVLDLFEDNYNGSASSTP    179
beta-ENaC_Hyperoodon_ampullatus                         EAVLERILGPELSHVNTTRVLNLTIWNHMPLVFINEQNPHHPVVLDLFEDNYNGSASSSP    179
beta-ENaC_Mesoplodon_bidens                             EAVLERILGPELSHVNTTRVLNLTIWNHMPLVFINEQNPHHPVVLDLFEDNYNGSASSSP    179
beta-ENaC_Ziphius_cavirostris                           EAVLERILGPELSHVNTTRVLNLTIWNHMPLVFINEQNPHHPVVLDLFEDNYNGSASSSP    179
beta-ENaC_Platanista_gangetica                          EAVLERILGPELSHVNATRALNLTIWNHMPLVFISEENPHHPVVLDLFEDNYNGSASSSP    179
beta-ENaC_Platanista_minor                              EAVLERILGPELSHVNATRALNLTIWNHMPLVFISEENPHHPVVLDLFEDNYNGSASSSP    179
beta-ENaC_Kogia_breviceps                               EAVLERILGPELSHVNATRALNLTIWNHMPLVFINEQNPHHPVVLDLFEDNYNGSASSSP    179
beta-ENaC_Physeter_catodon                              EAVLERILGPELSHVNATRALNLTIWNHMPLVFINEQNPHHPVVLDLFEDNYNGSASSSP    179
beta-ENaC_Balaenoptera_musculus                         EAVLERILGPELSHGNATRALNLTVWNYMPLVFINEQNPHHPVVLDLFEDDYNGSASSSP    179
beta-ENaC_Balaenoptera_acutorostrata_scammoni           EAVLERILGPELSHVNATRALNLTIWNYMPLVFINEQNPHHPVVLDLFEDNYNGSASSSP    179
beta-ENaC_Eubalaena_japonica                            EAVLERILGPELSHVNTTRALNLTIWNYMPLVFINEQNPHHPVVLDLFEDNYNGSASSSP    179
beta-ENaC_Hippopotamus_amphibius                        EAVLERILGPELSHFNATRALNLTIWNHMPLVFINEQNPHHPVVLELFEDNYNGSASNSS    180
beta-ENaC_Tragulus_javanicus                            GAVLERILGPELSRVNTTRTLNLSIWRHTPLVFINEQNPHHPVVLDLFEDNYNGSTSSSS    179
beta-ENaC_Tragulus_kanchil                              GAVLERILGPELSRVNTTRTLNLSIWRHTPLVFINEQNPHHPVVLDLFEDNYNGSTSSSS    179
beta-ENaC_Antilocapra_americana                         EAVLGRILGPELSQVNATRALNLSIWHHTPLVFINEQNPHHPVVLDLFDDNFNGSASSSP    179
beta-ENaC_Giraffa_camelopardalis                        EAVLGRILGPELSQVNATRALNLSIWHHTPLVFINEQNPHHPLVLDLFEDNFNGSASSSP    179
beta-ENaC_Giraffa_tippelskirchi                         EAVLGRILGPELSQVNATRALNLSIWHHTPLVFINEQNPHHPVVLDLFEDNFNGSASSSP    179
beta-ENaC_Capreolus_pygargus                            EAVLERILGPELSQVNATRTLNLSIWHHMPLVFINEQNPHHPVVLDLFEDNFNGSASSSP    180
beta-ENaC_Cervus_elaphus                                EAVLGRILGPELSQVNATRTLNLSIWHHTPLVFINEQNPHHPVVLDLFEDNFNGSASSSP    179
beta-ENaC_Moschus_moschiferus                           EAVLGRILGPELSQVNATRALNLSIWHHTPLVFINEQNPHHPVVLDLFEDNVNGSASSSP    179
beta-ENaC_Moschus_berezovskii                           EAVLGRILGPELSQVNATRALNLSIWHHTPLVFINEQNPHHPVVLDLFEDNVNGSASSSP    179
beta-ENaC_Bos_grunniens                                 EAVLGRILGPELSQVNDTRALNLSIWHHTPLVFINEQNPHHPVVLDLFEDNFNGSASNSP    179
beta-ENaC_Bos_taurus                                    EAVLGRILGPELSQVNDTRALNLSIWHHTPLVFINEQNPHHPVVLDLFEDNFNGSASNSP    179
beta-ENaC_Bubalus_bubalis                               EAVLGRILGPELSQVNATRALNLSIWHHTPLVFINEQNPHHPVVLDLFEDNFNGSASNSP    179
beta-ENaC_Nanger_granti                                 EAVLERILGPELSQVNATRALNLSIWHHTPLVFINEQNPHHPVVLDLFEDNVNGSASSSP    179
beta-ENaC_Capra_hircus                                  EAVLGRILGPELSQVNATTALNLSIWHHTPLVFINEQNPHHPVVLDLFEDNFNGSASSSP    179
beta-ENaC_Ovis_aries                                    EAVLGRILGPELSQVNATTALNLSIWHHTPLVFINEQNPHHPVVLDLFEDNFNGSASSSP    179
beta-ENaC_Ovis_canadensis                               EAVLGRILGPELSQVNATTALNLSIWHHTPLVFINEQNPHHPVVLDLFEDNFNGSASSSP    179
beta-ENaC_Hippotragus_niger                             EAVLGRILGPELSQVNATRALNLSIWHHTPLVFINEQNPHHPVVLDLFEDNFNGSASSSP    180
beta-ENaC_Sus_scrofa                                    EAVLQRILAPELSHVNATETLNFTIWNHTPLVFINEQNPQHPVVLDLFEDNYNGSASSGP    179
beta-ENaC_Vicugna_pacos                                 EAVLERILAPELSHINATRALNFTIWNHTPLVFIDEQNPHQPVVLDLFGDNHNDSASSSL    179
beta-ENaC_Equus_callabus                                DAVLERILAPEPSHANATEALNFTIWNHTPLVLIDEQNPHKPVVLDLFGDNHNSSASSSP    179
                                                         *** ***.** *  * *  **:::*.: ***:*.*.**::*:**:** *. .. :..

beta-ENaC_Homo_sapiens                                  ASEKICNA-HGCKMAMRLCSLNRTQCTFRNFTSATQALTEWYILQATNIFAQVPQQELVE    237
beta-ENaC_Globicephala_melas                            APGRTCSA-HGCKVAMKLCSHNGTTCNFRNFSSATQAVTEWYALQATNIFAQVPNQELVA    238
beta-ENaC_Lagenorhynchus_obliquidens                    APGRICSA-HGCKVAMKLCSHNGTTCNFRNFSSATQAVTEWYALQATNIFAQVPNEELVA    238
beta-ENaC_Tursiops_truncatus                            APGRTCSA-HGCKVAMKLCSHNGTTCNFRNFSSATQAVTEWYALQATNIFAQVPNQELVA    238
beta-ENaC_Orcinus_orca                                  APGRTCSA-HGCKVAMKLCSHNGTTCNFRNFSSATQAVTEWYALQATNIFAQVPNQELVA    238
beta-ENaC_Phocoena_sinus                                APGRTCSA-HGCKVAMKLCSHNGTTCNFRNFSSATQAVTEWYALQATNIFAQVPNQELVA    238
beta-ENaC_Neophocaena_asiaeorientalis                   APGRTCSA-HGCKVAMKLCSHNGTTCNFRNFSSATQAVTEWYALQATNIFAQVPNQELVA    238
beta-ENaC_Monodon_monocerons                            APGRTCSA-HGCKVAMKLCSHNGTTCNFRNFSSATQAVTEWYALQATNIFAQVPNQELVA    239
beta-ENaC_Delphinapterus_leucas                         APGRTCSA-HGCKVAMKLCSHNGTTCNFRNFSSATQAVTEWYALQATNIFAQVPNQELVA    238
beta-ENaC_Pontoporia_blainvillei                        APGRTCSA-HGCKVAMRLCSHNGTTCSFRNFSSATQAVTEWYTLQATNIFAQVPNQELVA    238
beta-ENaC_Inia_geoffrensis                              APGRTCSA-HGCKVAMRLCSHNGTTCSFRNFSSATQAVTEWYTLQATNIFAQVPNQELVA    238
beta-ENaC_Lipotes_vexillifer                            APGRTCSA-HGCKVAMRLCSHNGTTCNFRNFSSATQAVTEWYTLQATNIFAQVPNKELVA    238
beta-ENaC_Hyperoodon_ampullatus                         APGRTCSA-RECKVAMRLCSHNGTTCNFRNFSSATQAVMEWYTLQATNIFAQVPNQELVA    238
beta-ENaC_Mesoplodon_bidens                             APGRTCSA-RGCKVAMRLCSHNGTTCNFRNFSSATQAVTEWYTLQATNIFAQVPNQELVA    238
beta-ENaC_Ziphius_cavirostris                           APGRTCSA-RGCKVAMRLCSHNGTTCNFRNFSSATQAVMEWYTLQATNIFAQVPHQELVA    238
beta-ENaC_Platanista_gangetica                          APGRTCSA-HGCKVAMRLCSHSGTTCNFRNFSSATQAVTEWYTLQATNIFAQVPNQELVA    238
beta-ENaC_Platanista_minor                              APGRTCSA-HGCKVAMRLCSHSGTTCNFRNFSSATQAVTEWYTLQATNIFAQVPNQELVA    238
beta-ENaC_Kogia_breviceps                               APGRTCSA-HECKVAMRLCSHNGTMCNFRNFSSATQAVTEWYMLQATNIFAQVPNQELVA    238
beta-ENaC_Physeter_catodon                              APGRTCSA-HECKVAMRLCSHNGTMCNFRNFSSATQAVTEWYTLQATNIFAQVPNQELVA    238
beta-ENaC_Balaenoptera_musculus                         APGRTCSA-QGCKVAMRLCSHNGTTCNFRNFSSATQAVTEWYTLQATNIFAQVPKQELVA    238
beta-ENaC_Balaenoptera_acutorostrata_scammoni           APGRTCSA-QGCKVAMRLCSHNGTTCNFRNFSSATQAVTEWYTLQATNIFAQVPKQELVA    238
beta-ENaC_Eubalaena_japonica                            APERTCSA-QGCKVAMRLCSQNGTTCNFRNFSSATQAVTEWYTLQATNIFAQVPKQELVA    238
beta-ENaC_Hippopotamus_amphibius                        TPGKTCSAHHGCKVAMRLCSHNGTTCTFRNFSSATQAVTEWYTLQATNIFAQVPNQELVA    240
beta-ENaC_Tragulus_javanicus                            APGRPCSA-HRCKIAMRLCSHNGTACTFRNFSSATQAVTEWYTLQATNIFAQVPNQELVA    238
beta-ENaC_Tragulus_kanchil                              APGRPCSA-HRCKIAMRLCSHNGTACTFRNFSSATQAVTEWYTLQATNIFAQVPNQELVA    238
beta-ENaC_Antilocapra_americana                         APGRPCSA-HRCKVAMRLCSHNGTMCNFRNFSSATQAVMEWYTLQATNIFAQVPKQELVA    238
beta-ENaC_Giraffa_camelopardalis                        APGRPCSA-HRCKVAMRLCSHNGTTCTFRNFSSATQAVTEWYALQATNIFAQVPNQELVA    238
beta-ENaC_Giraffa_tippelskirchi                         APGRPCSA-HRCKVAMRLCSHNGTTCTFRNFSSATQAVTEWYALQATNIFAQVPNQELVA    238
beta-ENaC_Capreolus_pygargus                            APGRPCSA-HRCKVAIRLCSHNGTTCTFRNFSSATQAVTEWYTLQATNIFAQVPNPELVA    239
beta-ENaC_Cervus_elaphus                                APGRPCSA-HRCKVAMRLCSHNGTTCTFRNFSSATQAVTEWYTLQATNIFAQVPNQELVA    238
beta-ENaC_Moschus_moschiferus                           APGRPCSA-HRCKVAIRLCSHNGTTCTFRNFSSATQAVTEWYSLQATNIFAQVPNQELVA    238
beta-ENaC_Moschus_berezovskii                           APGRPCSA-HRCKVAIRLCSHNGTTCTFRNFSSATQAVTEWYSLQATNIFAQIPNQELVA    238
beta-ENaC_Bos_grunniens                                 APGRPCSA-HRCKVAMRLCSHNGTTCTFRNFSSATQAVTEWYTLQATNIFAQVPNQELVA    238
```

```
beta-ENaC_Bos_taurus                              APGRPCSA-HRCKVAMRLCSHNGTTCTFRNFSSATQAVTEWYTLQATNIFAQVPNQELVA    238
beta-ENaC_Bubalus_bubalis                         APGRPCSA-HRCKVAMRLCSHNGTTCTFRNFSSATQAVTEWYTLQATNIFAQVPNQELVA    238
beta-ENaC_Nanger_granti                           APGRPCSA-HRCKVAMRLCSHNGTTCTFRNFSSATQAVTEWYTLQATNIFAQVPNQELVA    238
beta-ENaC_Capra_hircus                            APGRPCSA-HRCKVAMRLCSHNGTTCTFRNFSSATQAVTEWYTLQATNIFAQVPNQELVA    238
beta-ENaC_Ovis_aries                              APGRPCSA-HRCKVAMRLCSHNGTTCTFRNFSSATQAVTEWYTLQATNIFAQVPNQELVA    238
beta-ENaC_Ovis_canadensis                         APGRPCSA-RRCKVAMRLCSHNGTTCTFRNFSSATQAVTEWYTLQATNIFAQVPNQELVA    238
beta-ENaC_Hippotragus_niger                       APGRPCSA-HRCKVAMRLCSHNGTTCTFRNFSSATQAVTEWYTLQATNIFAQVPNQELVA    239
beta-ENaC_Sus_scrofa                              APERTCRA-PGCKIAMKLCSHNGTTCTFRSFSSATRAVTEWYTLQATNIFSQVPNKELVA    238
beta-ENaC_Vicugna_pacos                           APGRSCSA-HACKVAMRLCSRGGTLCTFRNFSSATQAVTEWYTLQATNIFSQVPRRELVT    238
beta-ENaC_Equus_callabus                          APGRTCDA-QGCKVAMRLCSLNGTVCTFRNFTSATQAVMEWYVLQATNIFSQVPRQELVA    238
                                                  :  : * *   **:*::*** . * *.**.*:***:*: *** *******:*:*. ***

beta-ENaC_Homo_sapiens                            MSYPGEQMILACLFGAEPCNYRNFTSIFYPHYGNCYIFNWGMTEKALPSANPGTEFGLKL    297
beta-ENaC_Globicephala_melas                      MGYSAEQLILACLFGAEACTYRNFTRIFHPDYGNCYIFNWGMKEKALPSANPGAEFGLKL    298
beta-ENaC_Lagenorhynchus_obliquidens              MGYSAEQLILACLFGAEACTYRNFTRIFHPDYGNCYIFNWGMKEKALPSANPGAEFGLKL    298
beta-ENaC_Tursiops_truncatus                      MGYSAEQLILACLFGAEACTYRNFTRIFHPDYGNCYIFNWGMKEKALPSANPGAEFGLKL    298
beta-ENaC_Orcinus_orca                            MGYSAEQLILACLFGAEACTYRNFTRIFHPDYGNCYIFNWGMKEKALPSANPGAEFGLKL    298
beta-ENaC_Phocoena_sinus                          MGYSAEQLILACLFGAEACTYRNFTRIFHPDYGNCYIFNWGMKEKALPSANPGAEFGLKL    298
beta-ENaC_Neophocaena_asiaeorientalis             MGYSAEQLILACLFGAEACTYRNFTRIFHPDYGNCYIFNWGMKEKALPSANPGAEFGLKL    298
beta-ENaC_Monodon_monocerons                      MGYSAEQLILACLFGAEACTYRNFTPIFHPDYGNCYIFNWGMKEKALPSANPGAEFGLKL    299
beta-ENaC_Delphinapterus_leucas                   MGYSAEQLILACLFGAEACTYRNFTPIFHPDYGNCYIFNWGMKEKALPSANPGAEFGLKL    298
beta-ENaC_Pontoporia_blainvillei                  MGYPAEQLILACLFGAEPCTYRNFTPIFHPDYGNCYIFNWGMKEKALPSANPGAEFGLKL    298
beta-ENaC_Inia_geoffrensis                        MGYPAEQLILACLFGAEPCTYRNFTPIFHPDYGNCYIFNWGMTEKALPSANPGAEFGLKL    298
beta-ENaC_Lipotes_vexillifer                      MGYPAEQLILACLFGAEPCTYRNFTPIFNPDYGNCYIFNWGMKEKALPSANPGDDFGLKL    298
beta-ENaC_Hyperoodon_ampullatus                   MGYPAEQLILACLFGAEPCNYRNFTHIFHPEYGNCYIFNWGMREKALPSANPGAEFGLKL    298
beta-ENaC_Mesoplodon_bidens                       MGYPAEQLILACLFGAEPCNYRNFTHIFHPEYGNCYIFNWGMREKALPSANPGAEFGLKL    298
beta-ENaC_Ziphius_cavirostris                     MGYPAERLILACLFGAEPCNYRNFTHIFHPEYGNCYIFNWGMREKALPSANPGAEFGLKL    298
beta-ENaC_Platanista_gangetica                    MGYPAERLILACLFGAEPCNYRNFTPIFHPDYGNCYIFNWGMTEKALPSANPGAEFGLKL    298
beta-ENaC_Platanista_minor                        MGYPAERLILACLFGAEPCNYRNFTPIFHPDYGNCYIFNWGMTEKALPSANPGAEFGLKL    298
beta-ENaC_Kogia_breviceps                         MGYPAERLILACLFGAEPCNYRNFTPIFDPDYGNCYIFNWGMTEKALPSANPGAEFGLKL    298
beta-ENaC_Physeter_catodon                        MGYPAERLILACLFGAEPCNYRNFTPIFDPDYGNCYIFNWGMTEKALPSANPGAEFGLKL    298
beta-ENaC_Balaenoptera_musculus                   MGYPAERLILACLFGAEPCNYRNFTPIFHPDYGNCYIFNWGMREKALPSANPGAEFGLKL    298
beta-ENaC_Balaenoptera_acutorostrata_scammoni     MGYPAERLILACLFGAEPCNYRNFTPIFHPDYGNCYIFNWGMREKALPSANPGAEFGLKL    298
beta-ENaC_Eubalaena_japonica                      MGYPAERLILACLFGAEPCNYRNFTPIFHPDYGNCYIFNWGMREKALPSANPGAEFGLKL    298
beta-ENaC_Hippopotamus_amphibius                  MGYPAERLILACLFGAEPCNYRNFTPIFHPDYGNCYIFNWGMTEKALPSANPGAEFGLKL    300
beta-ENaC_Tragulus_javanicus                      MGYPAERLILACLFGAEPCNYRNFTRIFHPDYGNCYIFNWGMTEKALPSANPGTEFGLKL    298
beta-ENaC_Tragulus_kanchil                        MGYPAERLILACLFGAEPCNYRNFTRIFHPDYGNCYIFNWGMTEKALPSANPGTEFGLKL    298
beta-ENaC_Antilocapra_americana                   MGYPAERLILACLFGAEPCNYRNFTPIFHPDYGNCYIFNWGMREKALPSANPGTEFGLKL    298
beta-ENaC_Giraffa_camelopardalis                  MGYPAERLILACLFGAEPCNYRNFTPIFHPDYGNCYIFNWGMTEKALPSANPGTEFGLKL    298
beta-ENaC_Giraffa_tippelskirchi                   MGYPAERLILACLFGAEPCNYRNFTPIFHPDYGNCYIFNWGMTEKALPSANPGTEFGLKL    298
beta-ENaC_Capreolus_pygargus                      MGYPAERLILACLFGAEPCNHRNFTPIFHPDYGNCYIFNWGMTEKALPSANPGPEFGLKL    299
beta-ENaC_Cervus_elaphus                          MGYPAERLILACLFGAEPCNYRNFTPIFHPDYGNCYIFNWGMTEKALPSANPGTEFGLKL    298
beta-ENaC_Moschus_moschiferus                     MGYPAERLILACLFGAEPCNYRNFTPIFHPDYGNCYIFNWGMTEKALPSANPGTEFGLKL    298
beta-ENaC_Moschus_berezovskii                     MGYPAERLILACLFGAEPCNYRNFTPIFHPDYGNCYIFNWGMTEKALPSANPGTEFGLKL    298
beta-ENaC_Bos_grunniens                           MGYPAERLILACLFGAEPCNYRNFTPIFHPDYGNCYIFNWGMTEKALPSANPGTEFGLKL    298
beta-ENaC_Bos_taurus                              MGYPAERLILACLFGAEPCNYRNFTPIFHPDYGNCYIFNWGMTEKALPSANPGTEFGLKL    298
beta-ENaC_Bubalus_bubalis                         MGYPAERLILACLFGAEPCNYRNFTPIFHPDYGNCYIFNWGMTEKALPSANPGTEFGLKL    298
beta-ENaC_Nanger_granti                           MGYPAERLILACLFGAEPCNYRNFTPIFHPDYGNCYIFNWGMTEKALPSANPGTEFGLKL    298
beta-ENaC_Capra_hircus                            MGYPAERLILACLFGAEPCNYRNFTPIFHPDYGNCYIFNWGMTEKALPSANPGTEFGLKL    298
beta-ENaC_Ovis_aries                              MGYPAERLILACLFGAEPCNYRNFTPIFHPDYGNCYIFNWGMTEKALPSANPGTEFGLKL    298
beta-ENaC_Ovis_canadensis                         MGYPAERLILACLFGAEPCNYRNFTPIFHPDYGNCYIFNWGMTEKALPSANPGTEFGLKL    298
beta-ENaC_Hippotragus_niger                       MGYPAERLILACLFGAEPCNYRNFTPIFHPDYGNCYIFNWGMTEKALPSANPGTEFGLKL    299
beta-ENaC_Sus_scrofa                              MGYPAERMILACLFGAEPCNYRNFTPIFHPDYGNCYIFNWGMKEKALPSANPGAEFGLKL    298
beta-ENaC_Vicugna_pacos                           MGYSGEQLILACLFGAEPCSYRNFTPIFHPDYGNCYIFNWGMTEKALLSSNPGAEFGLKL    298
beta-ENaC_Equus_callabus                          MSYPGEQLILACLFGAEPCSHRNFTSIFHPDYGNCHIFNWGMTEKALPSSNPGAEFGLKL    298
                                                  *.* .*::********* *.:**** ** *.****:****** **** *:*** :*****

beta-ENaC_Homo_sapiens                            ILDIGQEDYVPFLASTAGVRLMLHEQRSYPFIRDEGIYAMSGTETSIGVLVDKLQRMGEP    357
beta-ENaC_Globicephala_melas                      ILDVGQEDYVPFLTSTAGARLMLHEQMSYPFIKEEGIFAMSGMETSIGVLVDKLERKGEP    358
beta-ENaC_Lagenorhynchus_obliquidens              ILDVGQEDYVPFLTSTAGARLMLHEQMSYPFIKEEGIFAMSGMETSIGVLVDKLERKGEP    358
beta-ENaC_Tursiops_truncatus                      ILDVGQEDYVPFLTSTAGARLMLHEQMSYPFIKEEGIFAMSGMETSIGVLVDKLERKGEP    358
beta-ENaC_Orcinus_orca                            ILDVGQEDYVPFLTSTAGARLMLHEQMSYPFIKEEGIFAMSGMETSIGVLVDKLERKGEP    358
beta-ENaC_Phocoena_sinus                          ILDVGQEDYVPFLTSTAGARLMLHEQMSYPFIKEEGIFAMSGMETSIGVLVDKLERKGEP    358
beta-ENaC_Neophocaena_asiaeorientalis             ILDVGQEDYVPFLTSTAGARLMLHEQMSYPFIKEEGIFAMSGMETSIGVLVDKLERKGEP    358
beta-ENaC_Monodon_monocerons                      ILDVGQEDYVPFLTSTAGARLMLHEQMSYPFIKEEGIFAMSGMETSIGVLVDKLERKGEP    359
beta-ENaC_Delphinapterus_leucas                   ILDVGQEDYVPFLTSTAGARLMLHEQMSYPFIKEEGIFAMSGMETSIGVLVDKLERKGEP    358
beta-ENaC_Pontoporia_blainvillei                  ILDVDQEDYVPFLTSTAGARLMLHEQMSYPFIKEEGIFAMSGMETSIGVLVDKLERKGEP    358
beta-ENaC_Inia_geoffrensis                        ILDVGQEDYVPFLTSTAGARLMLHEQMSYPFIKEEGIFAMSGMETSIGVLVDKLERKGEP    358
beta-ENaC_Lipotes_vexillifer                      ILDVGQEDYVPFLTSTAGARLMLHEQMSYPFIKEEGIFAMSGMETSIGVLVDKLERKGEP    358
beta-ENaC_Hyperoodon_ampullatus                   ILDMGQEDYVPFLTTTAGARLMLHEQMSYPFIKEEGIFAMSGMETSIGVLVDKLERKGEP    358
beta-ENaC_Mesoplodon_bidens                       ILDMGQEDYVPFLTTTAGARLMLHEQMSYPFIKEEGIFAMSGMETSIGVLVDKLERKGEP    358
beta-ENaC_Ziphius_cavirostris                     ILDMGQEDYVPFLTTTAGARLMLHEQMSYPFIKEEGIFAMSGMETSIGVLVDKLERKGEP    358
beta-ENaC_Platanista_gangetica                    ILDVGQEDYVPFLTPTAGARLMLHEQMSYPFIKEEGIFAMSGMETSIGVLVDKLERKGEP    358
beta-ENaC_Platanista_minor                        ILDVGQEDYVPFLTPTAGARLMLHEQMSYPFIKEEGIFAMSGMETSIGVLVDKLERKGEP    358
beta-ENaC_Kogia_breviceps                         ILDMGQEDYVPFLTSTAGARLMLHEQMSYPFIKEEGIFAMSGMETSIGVLVDKLERKGEP    358
beta-ENaC_Physeter_catodon                        ILDMGQEDYVPFLTSTAGARLMLHEQMSYPFIKEEGIFAMSGMETSIGVLVDKLERKGEP    358
beta-ENaC_Balaenoptera_musculus                   ILDMGQEDYVPFLTSTAGARLMLHEQMSYPFIKEEGIFAMSGMETSIGVLVDRLERKGEP    358
beta-ENaC_Balaenoptera_acutorostrata_scammoni     ILDMGQEDYVPFLTSTAGARLMLHEQMSYPFIKEEGIFAMSGMETSIGVLVDRLERKGEP    358
beta-ENaC_Eubalaena_japonica                      ILDMGQEDYVPFLTSTAGARLMLHEQMSYPFIKEEGIFAMSGMETSIGVLVDRLERKGEP    358
beta-ENaC_Hippopotamus_amphibius                  ILDMGQEDYMPFLTSTAGARLMLHEQRSYPFIKEEGIYAMSGMETSIGVLVDKLERKGEP    360
beta-ENaC_Tragulus_javanicus                      ILDMGQEDYVPYLTPTAGARLLLHEQRSYPFIKEEGIYAMAGMETSIGVLVDKLQRKGKP    358
beta-ENaC_Tragulus_kanchil                        ILDMGQEDYVPYLTPTAGARLLLHEQRSYPFIKEEGIYAMAGMETSIGVLVDKLQRKGKP    358
beta-ENaC_Antilocapra_americana                   ILDMGQEDYVPFLTSTAGARLMLHEQRSYPFIKEEGIYAMAGMETSIGVLVDKLQRKGKP    358
beta-ENaC_Giraffa_camelopardalis                  ILDMGQEDYVPFLTSTAGARLMLHEQRSYPFIKEEGIYAMVGMETSIGVLVDKLQRKGEP    358
beta-ENaC_Giraffa_tippelskirchi                   ILDMGQEDYVPFLTSTAGARLMLHEQRSYPFIKEEGIYAMVGMETSIGVLVDKLQRKGEP    358
beta-ENaC_Capreolus_pygargus                      ILDMGQEDYVPFLTSTAGARLMLHEQRSYPFIKEEGIYAMAGMETSIGVLVDKLQRKGEP    359
beta-ENaC_Cervus_elaphus                          ILDMGQEDYVPFLTSTAGARLMLHEQRSYPFIKEEGIYAMAGMETSIGVLVDKLQRKGKP    358
beta-ENaC_Moschus_moschiferus                     ILDMGQEDYVPFLTSTAGARLMLHEQRSYPFIKEEGIYAMAGMETSIGVLVDKLQRKGEP    358
beta-ENaC_Moschus_berezovskii                     ILDMGQEDYVPFLTSTAGARLMLHEQRSYPFIKEEGIYAMAGMETSIGVLVDKLQRKGEP    358
beta-ENaC_Bos_grunniens                           ILDMGQEDYVPFLTSTAGARLMLHEQRSYPFIKEEGIYAMAGMETSIGVLVDKLQRKGEP    358
beta-ENaC_Bos_taurus                              ILDMGQEDYVPFLTSTAGARLMLHEQRSYPFIKEEGIYAMAGMETSIGVLVDKLQRKGEP    358
```

```
beta-ENaC_Bubalus_bubalis                          ILDMGQEDYVPFLTSTAGARLMLHEQRSYPFIKEEGIYAMAGMETSIGVLVDKLQRKGEP   358
beta-ENaC_Nanger_granti                            ILDMGQEDYVPFLTSTAGARLMLHEQRSYPFIKEEGIYAMAGMETSIGVLVDKLQRKGEP   358
beta-ENaC_Capra_hircus                             ILDMGQEDYVPFLTSTAGARLMLHEQRSYPFIKEEGIYAMAGMETSIGVLVDKLQRKGEP   358
beta-ENaC_Ovis_aries                               ILDMGQEDYVPFLTSTAGARLMLHEQRSYPFIKEEGIYAMAGMETSIGVLVDKLQRKGEP   358
beta-ENaC_Ovis_canadensis                          ILDMGQEDYVPFLTSTAGARLMLHEQRSYPFIKEEGIYAMAGMETSIGVLVDKLQRKGEP   358
beta-ENaC_Hippotragus_niger                        ILDMGQEDYVPFLTSTAGARLMLHEQRSYPFIKEEGIYAMAGMETSIGVLVDKLQRKGEP   359
beta-ENaC_Sus_scrofa                               ILDMGQKDYVPFLTSTGGVRLMLHEQRSFPFIKDEGIYAMSGTETSIGVLVDKLERKGEP   358
beta-ENaC_Vicugna_pacos                            ILDMGQEDYVPFLTSTAGTRLMLHEQRSYPFIKDEGIYAMSGTETSIGVMVDRLQRKGEP   358
beta-ENaC_Equus_callabus                           ILDIGQEDYVPFLTSTAGARLLLHEQRSYPFIKEEGIYAMSGTETSIGVLVDRLERKGEP   358
                                                   ***:.*:**:*:*: *.*.**:**** *:***::***:** * ******:**:*:* *:*

beta-ENaC_Homo_sapiens                             YSPCTVNGSEVPVQNFYS---DYNTTYSIQACLRSCFQDHMIRNCNCGHYLYPLPRGEKY   414
beta-ENaC_Globicephala_melas                       YSQCTVNGSDVPIRNLYS---DYNTTYSIQACIRSCFQDHMIRNCSCGHYLYPLPRGEKY   415
beta-ENaC_Lagenorhynchus_obliquidens               YSQCTVNGSDVPIRNLYS---DYNTTYSIQACIRSCFQDHMIRNCSCGHYLYPLPRGEKY   415
beta-ENaC_Tursiops_truncatus                       YSQCTVNGSDVPIRNLYS---DYNTTYSIQACIRSCFQDHMIRNCSCGHYLYPLPRGEKY   415
beta-ENaC_Orcinus_orca                             YSQCTVNGSDVPIRNLYS---DYNTTYSIQACIRSCFQDHMIRNCSCGHYLYPLPRGEKY   415
beta-ENaC_Phocoena_sinus                           YSQCTVNGSDVPIRNLYS---DYNTTYSIQACIRSCFQDHMIRNCSCGHYLYPLPRGEKY   415
beta-ENaC_Neophocaena_asiaeorientalis              YSQCTVNGSDVPIRNLYS---DYNTTYSIQACIRSCFQDHMIRNCSCGHYLYPLPRGEKY   415
beta-ENaC_Monodon_monocerons                       YSQCTVNGSDVPIRNLYS---DYNTTYSIQACIRSCFQDHMIRNCSCGHYLYPLPRGEKY   416
beta-ENaC_Delphinapterus_leucas                    YSQCTVNGSDVPIRNLYS---DYNTTYSIQACIRSCFQDHMIRNCSCGHYLYPLPRGEKY   415
beta-ENaC_Pontoporia_blainvillei                   YSQCTVNGSDVPIRNLYS---DYNTTYSIQACIRSCFQDHMIHNCSCGHYLYPLPHGEKY   415
beta-ENaC_Inia_geoffrensis                         YSQCTVNGSDVPIRNLYS---DYNTTYSIQACIRSCFQDHMIHNCSCGHYLYPLPHGEKY   415
beta-ENaC_Lipotes_vexillifer                       YSQCTVNGSDVPIRNLYS---DYNTTYSIQACIRSCFQDHMIHNCSCGHYLYPLPRGEKY   415
beta-ENaC_Hyperoodon_ampullatus                    YSQCTMNGSDVPIRNLYS---DYNTTYSIQACIGSCFQDHMIHNCSCGHYLYPLPRGEKY   415
beta-ENaC_Mesoplodon_bidens                        YSQCTMNGSDVPIRNLYS---DYNTTYSIQACIGSCFQDHMIHNCSCGHYLYPLPRGEKY   415
beta-ENaC_Ziphius_cavirostris                      YSQCTMNGSDVPIRNLYS---DYNTTYSIQACIGSCFQDHMIHNCSCGHYLYPLPRGEKY   415
beta-ENaC_Platanista_gangetica                     YSQCTMNGSDVPIRNLYS---DYNTTYSIQACIRSCFQDHMIRNCSCGHYLYPLPRGEKY   415
beta-ENaC_Platanista_minor                         YSQCTMNGSDVPIRNLYS---DYNTTYSIQACIRSCFQDHMIRNCSCGHYLYPLPRGEKY   415
beta-ENaC_Kogia_breviceps                          YSQCTVNGSDVPIRNLYS---DYNTTYSIQACIRSCFQDHMIHNCSCGHYLYPLPRGEKY   415
beta-ENaC_Physeter_catodon                         YSQCTVNGSDVPIRNLYS---DYNTTYSIQACIRSCFQDHMIHNCSCGHYLYPLPRGEKY   415
beta-ENaC_Balaenoptera_musculus                    YSQCTMNGSDVPIRNLYS---DYNTTYSIQACIRSCFQDHMIHNCSCGHYLYPLPRGEKY   415
beta-ENaC_Balaenoptera_acutorostrata_scammoni      YSQCTMNGSDVPIRNLYS---DYNTTYSIQACIRSCFQDHMIRNCSCGHYLYPLPRGEKY   415
beta-ENaC_Eubalaena_japonica                       YSQCTMNGSDVPIRNLYS---DYNTTYSIQACIRSCFQDHMIHNCSCGHYLYPLPRGEKY   415
beta-ENaC_Hippopotamus_amphibius                   YSQCTMNGSDVPIQNLYSS--DYNTTYSIQACIRSCFQDHMIRNCSCGHYLYPLPRGQKY   418
beta-ENaC_Tragulus_javanicus                       YSQCTKNGSDVPVPNLYS---NYNTTYSIQACIRSCFQDRMIRDCGCGHYLYPLPPGRRY   415
beta-ENaC_Tragulus_kanchil                         YSQCTKNGSDVPVPNLYS---NYNTTYSIQACIRSCFQDRMIRDCGCGHYLYPLPPGKRY   415
beta-ENaC_Antilocapra_americana                    YSQCTKNGSDVPIQNLYS---GYNTTYSIQACIRSCFQEHMIRECGCGHYLYPLPHEEKY   415
beta-ENaC_Giraffa_camelopardalis                   YSQCTKNGSDVPIKNLYS---NYNTTYSIQACIRSCFQEYMIRECGCGHYLYPLPHEKKY   415
beta-ENaC_Giraffa_tippelskirchi                    YSQCTKNGSDVPIKNLYS---NYNTTYSIQACIRSCFQEYMIRECGCGHYLYPLPHEKKY   415
beta-ENaC_Capreolus_pygargus                       YSQCTKNGSDVPVQNLYS---SYNTTYSIQACIRSCFQEHMIRECGCGHYLYPLPHKSKY   416
beta-ENaC_Cervus_elaphus                           YSQCTKNGSDVPIPNLYS---SYNTTYSIQACIRSCFQEHMIRECGCGHYLYPLPHKRKY   415
beta-ENaC_Moschus_moschiferus                      YSQCTKNGSDVPIQNLYS---NYNTTYSIQACIRSCFQEHMIRECGCGHYLYPLPRKRKY   415
beta-ENaC_Moschus_berezovskii                      YSQCTKNGSDVPIQNLYS---NYNTTYSIQACIRSCFQEHMIRECGCGHYLYPLPRKRKY   415
beta-ENaC_Bos_grunniens                            YSQCTKNGSDVPIQNLYS---NYNTTYSIQACIRSCFQEHMIRECGCGHYLYPLPHKRKY   415
beta-ENaC_Bos_taurus                               YSQCTKNGSDVPIQNLYS---NYNTTYSIQACIRSCFQEHMIRECGCGHYLYPLPHKRKY   415
beta-ENaC_Bubalus_bubalis                          YSQCTKNGSDVPIQNLYS---SYNTTYSIQACIRSCFQEHMIRECGCGHYLYPLPHKRKY   415
beta-ENaC_Nanger_granti                            YSQCTKNGSDVPIQNLYS---NYNTTYSIQACIRSCFQEHMIRECGCGHYLYPLPHKRKY   415
beta-ENaC_Capra_hircus                             YSQCTKNGSDVPIQNLYS---SYNTTYSIQACIRSCFQEHMIRECGCGHYLYPLPDKKKY   415
beta-ENaC_Ovis_aries                               YSQCTKNGSDVPIQNLYS---SYNTTYSIQACIRSCFQEHMIRECGCGHYLYPLPDKRKY   415
beta-ENaC_Ovis_canadensis                          YSQCTKNGSDVPIQNLYS---SYNTTYSIQACIRSCFQEHMIRECGCGHYLYPLPDKRKY   415
beta-ENaC_Hippotragus_niger                        YSQCTKNGSDVPIQNLYS---SYNTTYSIQACIRSCFQEHMIRECGCGHYLYPLPHKRKY   416
beta-ENaC_Sus_scrofa                               YSQCTMNGSDVPIQNLYSNYGDYNTTYSIQACIRSCFQEHMIRHCSCGHYLYPLPPGEKY   418
beta-ENaC_Vicugna_pacos                            YSQCTVNGSDVPIRNLYS---YYNTTYSIQACIRSCFQDHMITKCSCGHYLYPLPRGEKY   415
beta-ENaC_Equus_callabus                           YSRCTKNGSDVPIPNLYS---DHNTTYSIQACIHSCFQDHMIRNCSCGHYLYPLPRGEKY   415
                                                   ** ** ***:**: *:**    :*********: ****: ** .*.*********   :*

beta-ENaC_Homo_sapiens                             CNNRDFPDWAHCYSDLQMSVAQRETCIGMCKES NDTQYKMTISMADWPSEASEDWIFHV   474
beta-ENaC_Globicephala_melas                       CNNQEFPDWAYCYSALRTSLAQRETCIDVCKESCNDTQYKMTISMAVWPSESSEDWIFHV   475
beta-ENaC_Lagenorhynchus_obliquidens               CNNQEFPDWAYCYSALRTSLAQRETCIDVCKESCNDTQYKMTISMAVWPSESSEDWIFHV   475
beta-ENaC_Tursiops_truncatus                       CNNQEFPDWAYCYSALRTSLAQRETCIDVCKESCNDTQYKMTISMAVWPSESSEDWIFHV   475
beta-ENaC_Orcinus_orca                             CNNQEFPDWAYCYSALRTSLAQRETCIDVCKESCNDTQYKMTISMAVWPSESSEDWIFHV   475
beta-ENaC_Phocoena_sinus                           CNNQEFPDWAYCYSALRMSLAQRETCIDVCKESCNDTQYKMTISMAVWPSESSEDWIFHV   475
beta-ENaC_Neophocaena_asiaeorientalis              CNNQEFPDWAYCYSALRMSLAQRETCIDVCKESCNDTQYKMTISMAVWPSESSEDWIFHV   475
beta-ENaC_Monodon_monocerons                       CNNQEFPDWAYCYSALRMSLAQRETCIDVCKESCNDTQYKMTISMAVWPSESSEDWIFHV   476
beta-ENaC_Delphinapterus_leucas                    CNNQEFPDWAYCYSALRMSLAQRETCIDVCKESCNDTQYKMTISMAVWPSESSEDWIFHV   475
beta-ENaC_Pontoporia_blainvillei                   CNNQEFPDWAYCYSALRMSLAQRETCIDVCKESCNDTQYKMTISMAVWPSEASEDWIFHV   475
beta-ENaC_Inia_geoffrensis                         CNNQEFPDWAYCYSALRMSLAQRETCIDVCKESCNDTQYKMTISMAAWPSEASEDWIFHV   475
beta-ENaC_Lipotes_vexillifer                       CNNQEFPDWAYCYSALRMSLAQRETCIDVCKESCNDTQYKMTISMAVWPSEASEDWIFHV   475
beta-ENaC_Hyperoodon_ampullatus                    CNNREFPDWVYCYSALRMSLAQRETCIDICKESCNDTQYKMTISMAVWPSESSEDWIFHV   475
beta-ENaC_Mesoplodon_bidens                        CNNRDFPDWVYCYSALRMSLAQRETCIDICKESCNDTQYKMTISMAVWPSESSEDWIFHV   475
beta-ENaC_Ziphius_cavirostris                      CNNREFPDWVYCYSALRMSLAQRETCIDICKESCNDTQYKMTISMAVWPSESSEDWIFHV   475
beta-ENaC_Platanista_gangetica                     CNNQEFPDWAYCYSALRVSLAQRETCIDVCKESCNDTQYKMTISMAVWPSEASEDWIFHV   475
beta-ENaC_Platanista_minor                         CNNQEFPDWAYCYSALRVSLAQRETCIDVCKESCNDTQYKMTISMAVWPSEASEDWIFHV   475
beta-ENaC_Kogia_breviceps                          CNNQEFPDWAYCYSALRMSLAQRETCIDLCKESCNDTQYKMTISMAVWPSEASEDWIFHV   475
beta-ENaC_Physeter_catodon                         CNNQEFPDWAYCYSALRMSLAQRETCIDVCKESCNDTQYKMTISMAVWPSEASEDWIFHV   475
beta-ENaC_Balaenoptera_musculus                    CNNQEFPDWAYCYSALRMSLAQRETCIDVCKESCNDTQYKMTISMAVWPSEASEDWIFHV   475
beta-ENaC_Balaenoptera_acutorostrata_scammoni      CNNQEFPDWAYCYSALRMSLAQRETCIDVCKESCNDTQYKMTISMAVWPSEASEDWIFHV   475
beta-ENaC_Eubalaena_japonica                       CNNQEFPDWAYCYSALRMSLAQRETCIDVCKESCNDTQYKMTISMAVWPSEASEDWIFHV   475
beta-ENaC_Hippopotamus_amphibius                   CSNQEFPDWALCYSALRMSLAQRETCIDMCKESCNDTQYKMTISMAVWPSEASEDWIFHV   478
beta-ENaC_Tragulus_javanicus                       CHSQEFPDWAHCYSALRISMEQREDCIYTCKESCNDTQYKMTISMAVWPSEASEDWILHV   475
beta-ENaC_Tragulus_kanchil                         CHSQEFPDWAHCYSALRISMEQREDCIYTCKESCNDTQYKMTISMAVWPSEASEDWILHV   475
beta-ENaC_Antilocapra_americana                    CNNQEFPDWAHCYSALRISMAQRETCIYACKESCNDTQYKMTISMAVWPSEASEDWIFHV   475
beta-ENaC_Giraffa_camelopardalis                   CNNQDFPDWAHCYSALRISLAQRETCIYACKESCNDTQYKMTISMAVWPSEASEDWIFHV   475
beta-ENaC_Giraffa_tippelskirchi                    CNNQDFPDWAHCYSALRISLAQRETCIYACKESCNDTQYKMTISMAVWPSEASEDWIFHV   475
beta-ENaC_Capreolus_pygargus                       CNNQEFPDWAHCYSALRISMAQREACIYACKESCNDTQYKMTISMAVWPSEASEDWIFHV   476
beta-ENaC_Cervus_elaphus                           CNNQEFPDWAHCYSALRISMAQREACIYACKESCNDTQYKMTISMAVWPSEASEDWIFHV   475
beta-ENaC_Moschus_moschiferus                      CSNQEFPDWAHCYSALRISMAQRETCIYACKESCNDTQYKMTISMAVWPSEASEDWIFHV   475
beta-ENaC_Moschus_berezovskii                      CSNQEFPDWAHCYSALRISMAQRETCIYACKESCNDTQYKMTISMAVWPSEASEDWIFHV   475
beta-ENaC_Bos_grunniens                            CNNQEFPDWAHCYSALRISLAQRETCIYACKESCNDTQYKMTISMAVWPSEASEDWIFHV   475
beta-ENaC_Bos_taurus                               CNNQEFPDWAHCYSALRISLAQRETCIYACKESCNDTQYKMTISMAVWPSEASEDWIFHV   475
beta-ENaC_Bubalus_bubalis                          CNNQEFPDWAHCYSALRISMAQRETCIYACKESCNDTQYKMTISMAVWPSEASEDWIFHV   475
```

```
beta-ENaC_Nanger_granti                                    CNNQEFPDWAHCYSALRISMAQRETCIYTCKESCNDTQYKMTISMAVWPSEASEDWIFHV    475
beta-ENaC_Capra_hircus                                     CNNQEFPDWAHCYSALRISMAQRETCIYACKESCNDTQYKMTISMAVWPSEASEDWIFQV    475
beta-ENaC_Ovis_aries                                       CNNQEFPDWAHCYSALRISMAQRETCIYACKESCNDTQYKMTISMAVWPSEASEDWIFHV    475
beta-ENaC_Ovis_canadensis                                  CNNQEFPDWAHCYSALRISMAQRETCIYACKESCNDTQYKMTISMAVWPSEASEDWIFHV    475
beta-ENaC_Hippotragus_niger                                CNNQEFPDWAHCYSALRVSMAQREACIYTCKESCNDTQYKMTISMAVWPSEASEDWIFHV    476
beta-ENaC_Sus_scrofa                                       CNSQEFPDWAYCYSDLRMSLAQRETCIDVCKESCNDTQYKMTISMADWPSESSEDWIFHV    478
beta-ENaC_Vicugna_pacos                                    CSNREFPDWAYCYSALRMSLAQRESCIDVCKESCNDTQYKMTISMADWPSEASEDWIFHV    475
beta-ENaC_Equus_callabus                                   CNNQDFPDWAYCYSDLRINVAQRETCINLCKESCNDTQYKMTISMAEWPSEASEDWILHV    475
                                                           * .::****. *** *: .: *** **  ***************** ****:*****::*

                                                                                                   TM2
beta-ENaC_Homo_sapiens                                     LSQERDQSTNITLSRKGIVKLNIYFQEFNYRTIEESAANNIVWLLSNLGGQFGFWMGGSV    534
beta-ENaC_Globicephala_melas                               LSEERDQSPNITMNRKGVIKLNIYFQEYNYRTIEESAANNIVWLLSNLGGQFGFWMGGSV    535
beta-ENaC_Lagenorhynchus_obliquidens                       LSEERDQSPNITMNRKGVIKLNIYFQEYNYRTIEESAANNIVWLLSNLGGQFGFWMGGSV    535
beta-ENaC_Tursiops_truncatus                               LSEERDQSPNITMNRKGVIKLNIYFQEYNYRTIEESAANNIVWLLSNLGGQFGFWMGGSV    535
beta-ENaC_Orcinus_orca                                     LSEERDQSPNITMNRKGVIKLNIYFQEYNYRTIEESAANNIVWLLSNLGGQFGFWMGGSV    535
beta-ENaC_Phocoena_sinus                                   LSEERDQSPNITLNRKGVIKLNIYFQEYNYRTIEESAANNIVWLLSNLGGQFGFWMGGSV    535
beta-ENaC_Neophocaena_asiaeorientalis                      LSEERDQSPNITLNRKGVIKLNIYFQEYNYRTIEESAANNIVWLLSNLGGQFGFWMGGSV    535
beta-ENaC_Monodon_monocerons                               LSEERDQSPNITLNRKGIIKLNIYFQEYNYRTIEESAANNIVWLLSNLGGQFGFWMGGSV    536
beta-ENaC_Delphinapterus_leucas                            LSEERDQSPNITLNRKGIIKLNIYFQEYNYRTIEESAANNIVWLLSNLGGQFGFWMGGSV    535
beta-ENaC_Pontoporia_blainvillei                           LSEERDQSPNITLNRKGIIKLNIYFQEYNYRTIEESEANNIVWLLSNLGGQFGFWMGGSV    535
beta-ENaC_Inia_geoffrensis                                 LSEERDQSPNITLNRKGIIKLNIYFQEYNYRTIEESAANNIVWLLSNLGGQFGFWMGGSV    535
beta-ENaC_Lipotes_vexillifer                               LSEERDQSPNITLNRKGIIKLNIYFQEYNYRTIEESAANNIVWLLSNLGGQFGFWMGGSV    535
beta-ENaC_Hyperoodon_ampullatus                            LSQERDQSTNITLNRKGVIKLNIYFQEYNYRTIAESAANNIVWLLSNLGGQFGFWMGGSV    535
beta-ENaC_Mesoplodon_bidens                                LSQERDQSTNITLNRKGVIKLNIYFQEYNYRTIAESAANNIVWLLSNLGGQFGFWMGGSV    535
beta-ENaC_Ziphius_cavirostris                              LSQERDHSTNITLNRKGVIKLNIYFQEYNYRTIAESAANNIVWLLSNLGGQFGFWMGGSV    535
beta-ENaC_Platanista_gangetica                             LSQERDQSTNITLNRKGIIKLNIYFQEYNYRTIEESAANNIVWLLSNLGGQFGFWMGGSV    535
beta-ENaC_Platanista_minor                                 LSQERDQSTNITLNRKGIIKLNIYFQEYNYRTIEESAANNIVWLLSNLGGQFGFWMGGSV    535
beta-ENaC_Kogia_breviceps                                  LSQERDQSTNITLNRKGIIKLNIYFQEYNYRTIEESAANNIVWLLSNLGGQFGFWMGGSV    535
beta-ENaC_Physeter_catodon                                 LSQERDQSTNITLNRKGIIKLNIYFQEYNYRTIEESAANNIVWLLSNLGGQFGFWMGGSV    535
beta-ENaC_Balaenoptera_musculus                            LSQERDQSTNITLNRKGIIKLNIYFQEYNYRTIEESAANNIVWLLSNLGGQFGFWMGGSV    535
beta-ENaC_Balaenoptera_acutorostrata_scammoni              LSQERDQSTNITLNRKGIIKLNIYFQEYNYRTIEESAANNIVWLLSNLGGQFGFWMGGSV    535
beta-ENaC_Eubalaena_japonica                               LSQERDQSTNITLNRKGIIKLNIYFQEYNYRTIEESAANNIVWLLSNLGGQFGFWMGGSV    535
beta-ENaC_Hippopotamus_amphibius                           LSQERDQSTNITLSRKGIVKLNIYFQEFNYRTIGESAANNIVWLLSNLGGQFGFWMGGSV    538
beta-ENaC_Tragulus_javanicus                               LSQERDQSTNITISRKGIVKLNIYFQEFNYRTIEESAANNIVWLLSNLGGQFGFWMGGSV    535
beta-ENaC_Tragulus_kanchil                                 LSQERDQSTNITISRKGIVKLNIYFQEFNYRTIEESAANNIVWLLSNLGGQFGFWMGGSV    535
beta-ENaC_Antilocapra_americana                            LSQERDQSSNITLSRKGIVKLNIYFQEFNYRTIEESAANNIVWLLSNLGGQFGFWMGGSV    535
beta-ENaC_Giraffa_camelopardalis                           LSQERDQSSNITLSRKGIVKLNIYFQEFNYRTIEESAANNIVWLLSNLGGQFGFWMGGSV    535
beta-ENaC_Giraffa_tippelskirchi                            LSQERDQSSNITLSRKGIVKLNIYFQEFNYRTIEESAANNIVWLLSNLGGQFGFWMGGSV    535
beta-ENaC_Capreolus_pygargus                               LSQERDQSSNITLSRKGIVKLNIYFQEFNYRTIEESAANNIVWLLSNLGGQFGFWMGGSV    536
beta-ENaC_Cervus_elaphus                                   LSQERDQSSNITLSRKGIVKLNIYFQEFNYRTIEESAANNIVWLLSNLGGQFGFWMGGSV    535
beta-ENaC_Moschus_moschiferus                              LSQERDQSSNITLSRKGIVKLNIYFQEFNYRTIEESAANNIVWLLSNLGGQFGFWMGGSV    535
beta-ENaC_Moschus_berezovskii                              LSQERDQSSNITLSRKGIVKLNIYFQEFNYRTIEESAANNIVWLLSNLGGQFGFWMGGSV    535
beta-ENaC_Bos_grunniens                                    LSQERDQSSNITLSRKGIVKLNIYFQEFNYRTIEESAANNIVWLLSNLGGQFGFWMGGSV    535
beta-ENaC_Bos_taurus                                       LSQERDQSSNITLSRKGIVKLNIYFQEFNYRTIEESAANNIVWLLSNLGGQFGFWMGGSV    535
beta-ENaC_Bubalus_bubalis                                  LSQERDQSSNITLSRKGIVKLNIYFQEFNYRTIEESAANNIVWLLSNLGGQFGFWMGGSV    535
beta-ENaC_Nanger_granti                                    LSQERDQSSNITLSRKGIVKLNIYFQEFNYRTIEESAANNIVWLLSNLGGQFGFWMGGSV    535
beta-ENaC_Capra_hircus                                     LSQERDQSSNITLSRKGIVKLNIYFQEFNYRTIEESAANNIVWLLSNLGGQLGFWMGGSV    535
beta-ENaC_Ovis_aries                                       LSQERDQSSNITLSRKGIVKLNIYFQEFNYRTIEESAANNIVWLLSNLGGQFGFWMGGSV    535
beta-ENaC_Ovis_canadensis                                  LSQERDQSSNITLSRKGIVKLNIYFQEFNYRTIEESAANNIVWLLSNLGGQFGFWMGGSV    535
beta-ENaC_Hippotragus_niger                                LSQERDQSSNITLSRKGIVKLNIYFQEFNYRTIEESAANNIVWLLSNLGGQFGFWMGGSV    536
beta-ENaC_Sus_scrofa                                       LSEERDQSTNITLSRQGIVKLNIYFQEFNYRTIEESAANNIVWLLSNLGGQFGFWMGGSV    538
beta-ENaC_Vicugna_pacos                                    LSQERDQSTNITLSRKGIVKLNIYFQEFNYRTIEESAANNIVWLLSNLGGQFGFWMGGSV    535
beta-ENaC_Equus_callabus                                   LSQERDQSTNITLSRKGIVKLNIYFQEFNYRTIEESAANNIVWLLSNLGGQFGFWMGGSV    535
                                                           **:***:* ***:.*:*::********:***** ** **************:********

beta-ENaC_Homo_sapiens                                     LCLIEFGEIIIDFVWITIIKLVALAKSLRQRRAQASYAGPPPTVAELVEAHTNFGFQPDT    594
beta-ENaC_Globicephala_melas                               LCLIEFAEIIIDFVWITIIKLVALAKSLRRRRTQARYDGPPPTVAELVEAHTNFGFQPDT    595
beta-ENaC_Lagenorhynchus_obliquidens                       LCLIEFAEIIIDFVWITIIKLVALAKSLRRRRTQARYDGPPPTVAELVEAHTNFGFQPDT    595
beta-ENaC_Tursiops_truncatus                               LCLIEFAEIIIDFVWITIIKLAALAKSLRRRRTQARYDGPPPTVAELVEAHTNFGFQPDT    595
beta-ENaC_Orcinus_orca                                     LCLIEFAEIIIDFVWITIIKLVALAKSLRRKRTQARYDGPPPTVAELVEAHTNFGFQPDT    595
beta-ENaC_Phocoena_sinus                                   LCLIEFAEIIIDFVWITIIKLVALAKSLRRRRAQTCYDGPPPTVAELVEAHTNFGFQPDT    595
beta-ENaC_Neophocaena_asiaeorientalis                      LCLIEFAEIIIDFVWITIIKLVALAKSLRRRRAQACYDGPPPTVAELVEAHTNFGFQPDT    595
beta-ENaC_Monodon_monocerons                               LCLIEFAEIIIDFVWITIIKLVALAKSLRRRRAQACYDGPPPTVAELVEAHTNFGFQPDT    596
beta-ENaC_Delphinapterus_leucas                            LCLIEFAEIIIDFVWITIIKLVALAKSLRRRRAQACYDGPPPTVAELVEAHTNFGFQPDT    595
beta-ENaC_Pontoporia_blainvillei                           LCLIEFAEIVIDFVWITIIKLVALAKSLWRRRAQARYDGPPPTVAELVEAHTNFGFQPDT    595
beta-ENaC_Inia_geoffrensis                                 LCLIEFAEIIIDFVWITIIKLVALAKSLRQRRAQARYDGPPPTVAELVEAHTNFGFQPDT    595
beta-ENaC_Lipotes_vexillifer                               LCLIEFAEIIIDFVWITIIKLVALAKSLRRRRAEARYNGPPPTVAELVEAHTNFGFQPDR    595
beta-ENaC_Hyperoodon_ampullatus                            LCLIEFGEIIIDFLWITIIRLVAFAKRLRQRRSQACYDGPPPTVAELVEAHTNFGFQPDT    595
beta-ENaC_Mesoplodon_bidens                                LCLIEFGEIIIDFLWITIIRLVAFAKRLRQRRAQACYDGPPPTVAELVEAHTNFGFQPDT    595
beta-ENaC_Ziphius_cavirostris                              LCLIEFGEIIIDFLWITIIRLVAFAKRLRQRRAQACYDGPPPTVAELVEAHTNFGFQPDT    595
beta-ENaC_Platanista_gangetica                             LCLIEFGEIIIDFVWITIIRLVALTKSLRQRRAQARYDGPPPTVAELVEAHTNFGFQPDT    595
beta-ENaC_Platanista_minor                                 LCLIEFGEIIIDFVWITIIRLVALTKSLRQRRAQARYDGPPPTVAELVEAHTNFGFQPDT    595
beta-ENaC_Kogia_breviceps                                  LCLIEFGEIIIDFVWITIIRLVALAKSLRQRRARARYDGPPPTVAELVEAHTNFGFQPDR    595
beta-ENaC_Physeter_catodon                                 LCLIEFGEIIIDFVWITIIRLVALAKSLRQRRTRARYDGPPPTVAELVEAHTNFGFQPDT    595
beta-ENaC_Balaenoptera_musculus                            LCLIEFGEIIIDFVWITIIRLVALAKSLRQRRAQACYDGPPPTVAELVEAHTNFGFQPDT    595
beta-ENaC_Balaenoptera_acutorostrata_scammoni              LCLIEFGEIIIDFVWITIIRLVALAKSLRQRRAQACYDGPPPTVAELVEAHTNFGFQPDT    595
beta-ENaC_Eubalaena_japonica                               LCLIEFGEIIIDFVWITIIRLVALAKSLRRRRAQACYDGPPPTVAELVEAHTNFGFQPDT    595
beta-ENaC_Hippopotamus_amphibius                           LCLIEFGEIVIDFVWITIIRLVALAKSLRRKRAQARYDGPPPTVAELVEAHTNFGFQPDV    598
beta-ENaC_Tragulus_javanicus                               LCIIEFGEIIIDFVWITIIKLVALAKGTRQKQAQARYDGPPPTVAELVEAHTNFAFQPDS    595
beta-ENaC_Tragulus_kanchil                                 LCIIEFGEIIIDFVWITIIKLVALAKGTRQKQAQARYDGPPPTVAELVEAHTNFAFQPDS    595
beta-ENaC_Antilocapra_americana                            LCLIEFGEIIIDFVWITIIKLVALAKSVRQKRAQARYEGPPPTVAELVEAHTNFGFQPDS    595
beta-ENaC_Giraffa_camelopardalis                           LCLIEFGEIIIDFMWITIIKLVALAKSVRQKRAQARYEGPPPTVAELVEAHTNFGFQPDS    595
beta-ENaC_Giraffa_tippelskirchi                            LCLIEFGEIIIDFMWITIIKLVALAKSVRQKRAQARYEGPPPTVAELVEAHTNFGFQPDS    595
beta-ENaC_Capreolus_pygargus                               LCLIEFGEIIIDFVWITIIKLVALAKSVRQKRAQARYEGPPPTVAELVEAHTNFGFQPDS    596
beta-ENaC_Cervus_elaphus                                   LCLIEFGEIIIDFVWITIIKLVALAKSVRQKRAQARYQGPPPTVAELVEAHTNFGFQPDS    595
beta-ENaC_Moschus_moschiferus                              LCLIEFGEIIIDFVWITIIKLVALAKSVRQKRALARYDGPPPTVAELVEAHTNFGFQPDP    595
beta-ENaC_Moschus_berezovskii                              LCLIEFGEIIIDFVWITIIKLVALAKSVRQKRALARYDGPPPTVAELVEAHTNFGFQPDP    595
beta-ENaC_Bos_grunniens                                    LCLIEFGEIIIDFVWITIIKLVALAKSVRQKQAQARYEGPPPTVAELVEAHTNFGFQPDL    595
beta-ENaC_Bos_taurus                                       LCLIEFGEIIIDFVWITIIKLVALAKSVRQKRAQARYEGPPPTVAELVEAHTNFGFQPDL    595
beta-ENaC_Bubalus_bubalis                                  LCLIEFGEIIIDFVWITIIKLVALAKSVRQKRAQARYDGPPPTVAELVEAHTNFGFQPDL    595
```

```
beta-ENaC_Nanger_granti                            LCLIEFGEIIIDFVWITIIKLVALAKGVRQKRAQARYDGPPPTVAELVEAHTNFGFQPDS    595
beta-ENaC_Capra_hircus                             LCLIEFGEIIIDFVWITIIKLVALAQSMRQKRAQARYDGPPPTVAELVEAHTNFGFQPDS    595
beta-ENaC_Ovis_aries                               LCLIEFGEIIIDFVWITIIKLVALAKSVRQKRAQARYNGPPPTVAELVEAHTNFGFQPDS    595
beta-ENaC_Ovis_canadensis                          LCLIEFGEIIIDFVWITIIKLVALAKSVQQKRAQARYDGPPPTVAELVEAHTNFGFQPDS    595
beta-ENaC_Hippotragus_niger                        LCLIEFGEIIIDFVWITIIRLVALAKGVRQKRAQARYDGPPPTVAELVEAHTNFGFQPDL    596
beta-ENaC_Sus_scrofa                               LCLIEFAEIIIDFVWITIIKLVALAKSLRQRRAQARYVGPPPTVAELVEAHTNFSFQPDT    598
beta-ENaC_Vicugna_pacos                            LCLIEFGEIIIDFVWITVIKLVALAKSLRQRRAQARYAGPPPTVAELVEAHTNFSFQPDV    595
beta-ENaC_Equus_callabus                           LCLIEFGEIIIDFVWITVIKLVAFSKSLQRRRAQAHYASPPPTVAELVEAHTNFGFQPDV    595
                                                   **:***.**:***:***:*:*.*:::   :::: : * .***************.****

beta-ENaC_Homo_sapiens                             APRSPNTGPYPSEQALPIPGTPPPNYDSLRLQPLDVIESDSEGDAI    640
beta-ENaC_Globicephala_melas                       ATHGPDAGPHPHEQTPPIPGTPPPNYDSLRLQPLDVIESDNEGDAI    641
beta-ENaC_Lagenorhynchus_obliquidens               ATHGPDAGPHPHEQTPPIPGTPPPNYDSLRLQPLDVIESDNEGDAI    641
beta-ENaC_Tursiops_truncatus                       ATHGPDAGPHPHEQTPPIPGTPPPNYDSLRLQPLDVIESDNEGDAI    641
beta-ENaC_Orcinus_orca                             ATHGPDAGPHPHEQTPPIPGTPPPNYDSLRLQPLDVIESDNEGDAI    641
beta-ENaC_Phocoena_sinus                           APHGPDAGPHAHEQTPPIPGTPPPNYDSLRLQPLDVTESDNEGDAI    641
beta-ENaC_Neophocaena_asiaeorientalis              APHGPDAGPHPHEQTPPIPGTPPPNYDSLRLQPLDVTESDNEGDAI    641
beta-ENaC_Monodon_monocerons                       ATHGPDAGPHPHEQTPPIPGTPPPNYDSLRLQPLDVTESDNEGDAI    642
beta-ENaC_Delphinapterus_leucas                    ATHGPDAGPHPHEQTPPIPGTPPPNYDSLRLQPLDVTESDNEGDAI    641
beta-ENaC_Pontoporia_blainvillei                   ATHGPDAGPHPHEQTPPIPGTPPPNYDSLRLQPLDVIESDNEGDAI    641
beta-ENaC_Inia_geoffrensis                         ATHSPDAGPHPHKQTPPIPGTPPPNYDSLRLQPLDVIESDNEGDAI    641
beta-ENaC_Lipotes_vexillifer                       ATHGPDAGPHPHEQTPPIPGTPPPNYDSLRLRPLDVIESDNEGDAI    641
beta-ENaC_Hyperoodon_ampullatus                    ATHGPDAGPHPHEQTPPIPGTPPPNYDSLRLQPLDIIESDNEGDAI    641
beta-ENaC_Mesoplodon_bidens                        ATHGPDAGPHPHEQTPPIPGTPPPNYDSLRLQPLDIIESDNEGDAI    641
beta-ENaC_Ziphius_cavirostris                      ATHGPDAGPHPHEQTLPIPGTPPPNYDSLRLQPLDTIESDNEGDAI    641
beta-ENaC_Platanista_gangetica                     ATHGPDAGPHPREQTPSIPGTPPPNYDSLRLQPLDVIESDNEGDAI    641
beta-ENaC_Platanista_minor                         ATHGPDAGPHPREQTPSIPGTPPPNYDSLRLQPLDVIESDNEGDAI    641
beta-ENaC_Kogia_breviceps                          ATHGPDARPHPHEQAPPIPGTPPPNYDSLRLQPLDVIESDNEDDAI    641
beta-ENaC_Physeter_catodon                         ATHGPDARPHPHEQTPPIPGTPPPNYDSLRLQPLDVIESDNEGDAI    641
beta-ENaC_Balaenoptera_musculus                    ATHGPDAGPHPHEQTPPIPGTPPPNYDSLRLQPLDVIESDNEGDAI    641
beta-ENaC_Balaenoptera_acutorostrata_scammoni      ATHGPDAGPHPHEQTPPIPGTPPPNYDSLRLQPLDVIESDNEGDAI    641
beta-ENaC_Eubalaena_japonica                       ATHGPDAGPHPHEQTPPIPGTPPPNYDSLRLQPLDVIESDNEGDAI    641
beta-ENaC_Hippopotamus_amphibius                   ATRGPSAGPHSHEPTPPIPGTPPPNYDSLRLQPLDVIESDSEGDAV    644
beta-ENaC_Tragulus_javanicus                       ATPGPDTGPYTQEQTLPIPGTPPPNYDSLRLQPLDVIESDSEADAI    641
beta-ENaC_Tragulus_kanchil                         ATPGPDTGPYTQEQTLPIPGTPPPNYDSLRLQPLDVIESDSEADAI    641
beta-ENaC_Antilocapra_americana                    ATPGPDAGAYPHEQNPPIPGTPPPNYDSLRLQPLDIIESDSEGDAI    641
beta-ENaC_Giraffa_camelopardalis                   AMPGPNVGAYPHEQNPPIPGTPPPNYDSLRLQPLDVIESDSEGDAI    641
beta-ENaC_Giraffa_tippelskirchi                    AMPGPNVGAYPHEQNPPIPGTPPPNYDSLRLQPLDVIESDSEGDAI    641
beta-ENaC_Capreolus_pygargus                       ATTSPDAGTYTREQNPPIPGTPPPNYDSLRLQPLDVIESDSEGDAI    642
beta-ENaC_Cervus_elaphus                           ATPGPDAGAYPHEQNPPIPGTPPPNYDSLRLQPLDVIESDSEGDAI    641
beta-ENaC_Moschus_moschiferus                      ATPGPDAGAYPHEQNPPIPGTPPPNYDSLRLQPLDVIESDSDGDAI    641
beta-ENaC_Moschus_berezovskii                      ATPGPDAGAYPHEQNPPIPGTPPPNYDSLRLQPLDVIESDSDGDAI    641
beta-ENaC_Bos_grunniens                            ATPGPDVGAYPHEQNPPIPGTPPPNYDSLRLQPLDVIESDSEGDAI    641
beta-ENaC_Bos_taurus                               ATPGPDVEAYPHEQNPPIPGTPPPNYDSLRLQPLDVIESDSEGDAI    641
beta-ENaC_Bubalus_bubalis                          ATPGPDVGAYPHEQNPPIPGTPPPNYDSLRLQPLDVIESDSEGDAI    641
beta-ENaC_Nanger_granti                            ATPGPDAGAYPHEQNPPIPGTPPPNYESLRLQPLDVIESDSEGDAI    641
beta-ENaC_Capra_hircus                             AMPGPDAGAYSREQNPPIPGTPPPNYDSLRLQPLDVIESDSEGDAI    641
beta-ENaC_Ovis_aries                               AMPGPDAGAYRREQNPPIPGTPPPNYDSLRLQPLDVIESDSEGDAI    641
beta-ENaC_Ovis_canadensis                          AMPGPDAGAYRREQNPPIPGTPPPNYDSLRLQPLDVIESDSEGDAI    641
beta-ENaC_Hippotragus_niger                        ATPGPDTGAYPREQNPPIPGTPPPNYDSLRLQPLDVIESDSEGDAI    642
beta-ENaC_Sus_scrofa                               PAPSADPGPYVQEQAPPIPGTPPPNYDSLRLQPLDVIESDGEGDAV    644
beta-ENaC_Vicugna_pacos                            VTPGPSAGPYPDEQLPPIPGTPPPNYDSLRLQPLDVIESDTEGDAV    641
beta-ENaC_Equus_callabus                           ISHSPTTGTYPDEQPVPIPGTPPPNYDSLRLQPLDVVESDNEGDAI    641
                                                      .     :  :    *********:****:***  *** : **:
```

**γ-ENaC (*SCNN1G*) amino acid sequence alignment** created with Clustal Omega (v.1.2.4, accessed 24.03.2024). Key structural motifs are highlighted in the human ENaC subunit based on the Cryo-EM derived structure (Noreng et al. 2018). Transmembrane domains (TM1/TM2) are highlighted in yellow. An N-Terminal HG-motif affecting ENaC open probability is marked in magenta. Protease cleavage sites are highlighted in blue. Cysteines involved in tertiary structure formation are indicated in red. Residues putatively forming the selectivity filter within TM2 are shown in red font. The C-terminal PPPxY motif regulating membrane abundance is shown in **gray**.

```
                                                                                                 TM1
gamma-ENaC_Homo_sapiens                     MAPGEKIKAKIKKNLPVTGPQAPTIKELMRWYCLNTNTHGCRRIVVSRGRLRRLLWIGFT  60
gamma-ENaC_Globicephala_melas               MAAGEKIKAKIKKNLPVRGPQAPTIKELMRWYCLNTNTHGCRRIVVSRGRLRRLLWILFT  60
gamma-ENaC_Lagenorhynchus_obliquidens       MAAGEKIKAKIKKNLPVRGPQAPTIKELMRWYCLNTNTHGCRRIVVSRGRLRRLLWILFT  60
gamma-ENaC_Tursiops_truncatus               MAAGEKIKAKIKKNLPVRGPQAPTIKELMRWYCLNTNTHGCRRIVVSRGRLRRLLWILFT  60
gamma-ENaC_Orcinus_orca                     MAAGEKIKAKIKKNLPVRGPQAPTIKELMRWYCLNTNTHGCRRIVVSRGRLRRLLWILFT  60
gamma-ENaC_Phocoena_sinus                   MAAGEKIKAKIKKNLPVRGPQAPTIKELMRWYCLNTNTHGCRRIVVSRGRLRRLLWILLT  60
gamma-ENaC_Neophocaena_asiaeorientalis      MAAGEKIKAKIKKNLPVRGPQAPTIKELMRWYCLNTNTHGCRRIVVSRGRLRRLLWILLT  60
gamma-ENaC_Monodon_monocerons               MAAGEKIKAKIKKNLPVRGPQAPTIKELMRWYCLNTNTHGCRRIVVSRGRLRRLLWILFT  60
gamma-ENaC_Delphinapterus_leucas            MAAGEKIKAKIKKNLPVRGPQAPTIKELMRWYCLNTNTHGCRRIVVSRGRLRRLLWILFT  60
gamma-ENaC_Pontoporia_blainvillei           MAAGEKIKAKIKKNLPVRGPQAPTIKELMRWYCLNTNTHGCRRIVVSRGRLRRLLWILFT  60
gamma-ENaC_Inia_geoffrensis                 MAAGEKIKAKIKKNLPVRGPQAPTIKELMRWYCLNTNTHGCRRIVVSRGRLRRLLWILFT  60
gamma-ENaC_Lipotes_vexillifer               MAAGEKIKAKIKKNLPVRGPQAPTIKELMRWYCLNTNTHGCRRIVVSRGRLRRLLWILFT  60
gamma-ENaC_Hyperoodon_ampullatus            MAAGEKIKAKIKENLPVRGPQAPTIKELMRWYCLNTNTHGCRRIVVSRGRLRRLLWILFT  60
gamma-ENaC_Mesoplodon_bidens                MAAGEKIKAKIKENLPVRGPQAPTIKELMRWYCLNTNTHGCRRIVVSRGRLRRLLWILFT  60
gamma-ENaC_Ziphius_cavirostris              MAAGEKIKAKIKENLPVRGPQAPTIRELMRWYCLNTNTHGCRRIVVSRGRLRRLLWILFT  60
gamma-ENaC_Platanista_gangetica             MAAGEKIKAKIKKNLPVRGPQAPTIKELMRWYCLNTNTHGCRRIVVSRGRLRRLLWILFT  60
gamma-ENaC_Platanista_minor                 MAAGEKIKAKIKKNLPVRGPQAPTIKELMRWYCLNTNTHGCRRIVVSRGRLRRLLWILFT  60
gamma-ENaC_Kogia_breviceps                  MAAGEKIKAKIKKNLPVRGPQAPTIKELMRWYCLNTNTHGCRRIVVSRGRLRRLLWILFT  60
gamma-ENaC_Physeter_catodon                 MAAGEKIKAKIKKNLPVRGPQAPTIKELMRWYCLNTNTHGCRRIVVSRGRLRRLLWILFT  60
gamma-ENaC_Balaenoptera_musculus            MAAGEKIKAKIKKNLPVRGPQAPTIKELMQWYCLNTNTHGCRRIVVSRGRLRRLLWILFT  60
gamma-ENaC_Eubalaena_japonica               MAAGEKIKAKIKKNLPVRGPQAPTIKELMQWYCLNTNTHGCRRIVVSRGRLRRLLWILFT  60
gamma-ENaC_Hippopotamus_amphibius           MAPGEKIKAKIKKNLPVTGPQAPTIKELMRWYCLNTNTHGCRRIVVSRGRLRRLLWILFT  60
gamma-ENaC_Tragulus_javanicus               MAPGEKIKAKIKKNLPVTGPQAPTIKELMRWYCLNTNTHGCRRIVVSRGRLRRLLWILFT  60
gamma-ENaC_Tragulus_kanchil                 MAPGEKIKAKIKKNLPVTGPQAPTIKELMRWYCLNTNTHGCRRIVVSRGRLRRLLWILFT  60
gamma-ENaC_Antilocapra_americana            MAPGEKIKAKIKKNLPVTGPQAPNIKELMQWYCLNTNTHGCRRIVVSRGRLRRLLWILFT  60
gamma-ENaC_Giraffa_camelopardalis           MAPGEKIKAKIKKNLPVTGPQAPNIKELMQWYCLNTNTHGCRRIVVSRGRLRRLLWILFT  60
gamma-ENaC_Giraffa_tippelskirchi            MAPGEKIKAKIKKNLPVTGPQAPNIKELMQWYCLNTNTHGCRRIVVSRGRLRRLLWILFT  60
gamma-ENaC_Capreolus_pygargus               MAPGEKIKAKIKKNLPVTGPQAPNIKELMQWYCLNTNTHGCRRIVVSRGRLRRLLWILFT  60
gamma-ENaC_Cervus_elaphus                   MAPGEKIKAKIKKNLPVTGPQAPNIKELMQWYCLNTNTHGCRRIVVSRGRLRRLLWILFT  60
gamma-ENaC_Moschus_moschiferus              MAPGEKIKAKIKKNLPVTGPQAPNIKELMQWYCLNTNTHGCRRIVVSRGRLRRLLWILFT  60
gamma-ENaC_Moschus_berezovskii              MAPGEKIKAKIKKNLPVTGPQAPNIKELMQWYCLNTNTHGCRRIVVSRGRLRRLLWILFT  60
gamma-ENaC_Bos_grunniens                    MAPGEKIKAKIKKNLPVTGPQAPNIKELMQWYCLNTNTHGCRRIVVSRGRLRRLLWILFT  60
gamma-ENaC_Bos_taurus                       MAPGEKIKAKIKKNLPVTGPQAPNIKELMQWYCLNTNTHGCRRIVVSRGRLRRLLWILFT  60
gamma-ENaC_Bubalus_bubalis                  MAPGEKIKAKIKKNLPVTGPQAPNIKELMQWYCLNTNTHGCRRIVVSRGRLRRLLWILFT  60
gamma-ENaC_Nanger_granti                    MAPGEKIKAKIKKNLPVTGPQAPNIKELMQWYCLNTNTHGCRRIVVSRGRLRRLLWILFT  60
gamma-ENaC_Kobus_leche                      MAPGEKIKAKIKKNLPVTGPQAPSIKELMHWYCLNTNTHGCRRIVVSRGRLRRLLWILFT  60
gamma-ENaC_Capra_hircus                     MAPGEKIKAKIKKNLPVTGPQAPNIKELMQWYCLNTNTHGCRRIVVSRGRLRRLLWILFT  60
gamma-ENaC_Ovis_aries                       MAPGEKIKAKIKKNLPVTGPQAPNIKELMQWYCLNTNTHGCRRIVVSRGRLRRLLWILFT  60
gamma-ENaC_Ovis_canadensis                  MAPGEKIKAKIKKNLPVTGPQAPNIKELMQWYCLNTNTHGCRRIVVSRGRLRRLLWILFT  60
gamma-ENaC_Oreamnos_americanus              MAPGEKIKAKIKKNLPVTGPQAPNIKELMQWYCLNTNTHGCRRIVVSRGRLRRLLWILFT  60
gamma-ENaC_Hippotragus_niger                MAPGEKIKAKIKKNLPVTGPQAPNIKELMQWYCLNTNTHGCRRIVVSRGRLRRLLWILFT  60
gamma-ENaC_Damaliscus_lunatus               MAPGEKIKAKIKKNLPVTGPQAPNIKELMQWYCLNTNTHGCRRIVVSRGRLRRLLWILFT  60
gamma-ENaC_Sus_scrofa                       MAPGEKIKAKIKKNLPVTGPQAPTIKELMQWYCLNTNTHGCRRIVVSRGRLRRLLWILFT  60
gamma-ENaC_Vicugna_pacos                    MAPGEKIKAKIKKNLPVKGPQAPTIKELMRWYCLNTNTHGCRRIVVSRGRLRRLLWILFT  60
gamma-ENaC_Equus_callabus                   MAPGEKIKAKIKKNLPVTGPQAPTIKELMRWYCLNTNTHGCRRIVVSRGRLRRLLWILFT  60
                                            ** *********:**** *****.*:***:*************************** :*


gamma-ENaC_Homo_sapiens                     LTAVALILWQCALLVFSFYTVSVSIKVHFRKLDFPAVTICNINPYKYSTVRHLLADLEQE  120
gamma-ENaC_Globicephala_melas               LTAVALIFWQCALLISSFYTVSVSIKVHFQKLDFPAVTICNINPYRYSAVRDLLAELEQE  120
gamma-ENaC_Lagenorhynchus_obliquidens       LTAVALIFWQCALLISSFYTVSVSIKVHFQKLDFPAVTICNINPYRYSAVRDLLAELEQE  120
gamma-ENaC_Tursiops_truncatus               LTAVALIFWQCALLISSFYTVSVSIKVHFQKLDFPAVTICNINPYRYSAVRDLLAELEQE  120
gamma-ENaC_Orcinus_orca                     LTAVALIFWQCALLISSFYTVSVSIKVHFQKLDFPAVTICNINPYRYSAVRDLLAELEQE  120
gamma-ENaC_Phocoena_sinus                   LTAVALIFWQCALLISSFYTVSVSIKVHFQKLDFPAVTICNINPYRYSAVRDLLAELEQE  120
gamma-ENaC_Neophocaena_asiaeorientalis      LTAVALIFWQCALLISSFYTVSVSIKVHFQKLDFPAVTICNINPYRYSAVRDLLAELEQE  120
gamma-ENaC_Monodon_monocerons               LTAVALIFWQCALLISSFYTVSVSIKVHFQKLDFPAVTICNINPYRYSAVRDLLAELEQE  120
gamma-ENaC_Delphinapterus_leucas            LTAVALIFWQCALLISSFYTVSVSIKVHFQKLDFPAVTICNINPYRYSAVRDLLAELEQE  120
gamma-ENaC_Pontoporia_blainvillei           LTAVALIFWQCALLISSFYTVSVSIKVHFQKLDFPAVTICNINPYRYSAVRDLLAELEQE  120
gamma-ENaC_Inia_geoffrensis                 LTAVALIFWQCALLISSFYTVSVSIKVHFQKLDFPAVTICNINPYRYSAVRDLLAELEQE  120
gamma-ENaC_Lipotes_vexillifer               LTAVALIFWQCALLISTFYTVSVSIKVHFQKLDFPAVTICNINPYRYSAVRHLLAELEQE  120
gamma-ENaC_Hyperoodon_ampullatus            LTAVALIFWQCALLISSFYTVSVSIKVHFQKLDFPAVTICNINPYRYSAVRDLLADLEQE  120
gamma-ENaC_Mesoplodon_bidens                LTAVALIFWQCALLISSFYTVSVSIKVHFQKLDFPAVTICNINPYRYSAVRDLLADLEQE  120
gamma-ENaC_Ziphius_cavirostris              LTAVALIFWQCALLISSFYTVSVSIKVHFQKLDFPAVTICNINPYRYSAVRDLLADLEQE  120
gamma-ENaC_Platanista_gangetica             LTAVALIFWQCALLISSFYTVSVSIKVHFQKLDFPAVTICNINPYRYSAVRDLLADLEQE  120
gamma-ENaC_Platanista_minor                 LTAVALIFWQCALLISSFYTVSVSIKVHFQKLDFPAVTICNINPYRYSAVRDLLADLEQE  120
gamma-ENaC_Kogia_breviceps                  LTAVALIFWQCALLISSFYTVSVSIKVHFQKLDFPAVTICNINPYRYSAVRDLLADLEQE  120
gamma-ENaC_Physeter_catodon                 LTAVALIFWQCALLISSFYTVSVSIKVHFQKLDFPAVTICNINPYRYSAVRDLLADLEQE  120
gamma-ENaC_Balaenoptera_musculus            LTAVALIFWQCALLISSFYTVSVSIKVHFQKLDFPAVTICNINPYRYSAVRDLLADLEKE  120
gamma-ENaC_Eubalaena_japonica               LTAVALIFWQCALLISSFYTVSVSIKVHFQKLDFPAVTICNINPYRYSAVQDLLADLEKE  120
```

```
gamma-ENaC_Hippopotamus_amphibius            LTAVALIFWQCALLISSFYTVSVSIKVHFQKLDFPAVTICNINPYKYSAVRDLLADLERE  120
gamma-ENaC_Tragulus_javanicus                LTAVALIFWQCALLISSFYTVSVSIKVHFQKLDFPAVTICNMNPFKYSAVQHLLADLEQE  120
gamma-ENaC_Tragulus_kanchil                  LTAVALIFWQCALLISSFYTVSVSIKVHFQKLDFPAVTICNMNPFKYSAVQHLLADLEQE  120
gamma-ENaC_Antilocapra_americana             LTAVALIFWQCALLISSFYTVSVSIKVHFQKLDFPAVTICNINPYKYSAVRPLLADLEQE  120
gamma-ENaC_Giraffa_camelopardalis            LTAVALILWQCALLISSFYTVSVSIKVHFQKLDFPAVTICNINPYKYSAVRHLLADLEQE  120
gamma-ENaC_Giraffa_tippelskirchi             LTAVALILWQCALLISSFYTVSVSIKVHFQKLDFPAVTICNINPYKYSAVRHLLADLEQE  120
gamma-ENaC_Capreolus_pygargus                LTAVALIFWQCALLISSFYTVSVSIKVHFQKLDFPAVTICNINPYKYSAVRHLLADLEQE  120
gamma-ENaC_Cervus_elaphus                    LTAVALIFWQCALLISSFYTVSVSIKVHFQKLDFPAVTICNINPYKYSAVRHLLADLEQE  120
gamma-ENaC_Moschus_moschiferus               LTAVALIFWQCALLISSFYTVSVSIKVHFQKLDFPAVTICNINPYRYSAVRHLLADLEQE  120
gamma-ENaC_Moschus_berezovskii               LTAVALIFWQCALLISSFYTVSVSIKVHFQKLDFPAVTICNINPYRYSAVRHLLADLEQE  120
gamma-ENaC_Bos_grunniens                     LTAVALIFWQCALLISSFYTVSVSIKVHFQKLDFPAVTICNINPYKYSAVRHLLADLEQE  120
gamma-ENaC_Bos_taurus                        LTAVALIFWQCALLISSFYTVSVSIKVHFQKLDFPAVTICNINPYKYSAVRHLLADLEQE  120
gamma-ENaC_Bubalus_bubalis                   LTAVGLIFWQCALLISSFYTVSVSIKVHFQKLDFPAVTICNINPYKYSAVRHLLADLEQE  120
gamma-ENaC_Nanger_granti                     LTAVALIFWQCSLLISSFYTVSVSIKVHFQKLDFPAVTICNINPYKYSAVRHLLADLERE  120
gamma-ENaC_Kobus_leche                       LTAVALIFWQCALLISSFYTVSVSIKVHFQKLDFPAVTICNINPYKYSAVRHLLGDLEQE  120
gamma-ENaC_Capra_hircus                      LTAVALIFWQCALLISSFYTVSVSIKVHFQKLDFPAVTICNINPYKYSAVRHLLADLEQE  120
gamma-ENaC_Ovis_aries                        LTAVALIFWQCALLISSFYTVSVSIKVHFQKLDFPAVTICNINPYKYSAVRHLLADLEQE  120
gamma-ENaC_Ovis_canadensis                   LTAVALIFWQCALLISSFYTVSVSIKVHFQKLDFPAVTICNINPYKYSAVRHLLADLEQE  120
gamma-ENaC_Oreamnos_americanus               LTAVALIFWQCALLISSFYTVSVSIKVHFQKLDFPAVTICNINPYKYSAVRHLLADLEQE  120
gamma-ENaC_Hippotragus_niger                 LTAVALIFWQCALLISSFYTVSVSIKVHFQKLDFPAVTICNINPYKYSAVRHLLADLEQE  120
gamma-ENaC_Damaliscus_lunatus                LTAVALIFWQCALLISSFYTVSVSIKVHFQKLDFPAVTICNINPYKYSAVRHLLADLEQE  120
gamma-ENaC_Sus_scrofa                        LTAVALIFWQCALLILSFYTVSVSIKVHFQKLDFPAVTICNINPYKYSAVRDLLAELEQE  120
gamma-ENaC_Vicugna_pacos                     LTAVALIFWQCALLILSFYTVSVSIKVHFQKLDFPAVTICNINPYKYSAVRDLLADLEQE  120
gamma-ENaC_Equus_callabus                    LTAVALIFWQCALLVISFYTVSVSIKIHFQKLDFPAVTICNINPYKYSAVRDLLADLEQE  120
                                             ****.**:***:**: :*********:**:***********:**::**:*: **.:**:*

gamma-ENaC_Homo_sapiens                      TREALKSLYGFPE--SRKRREAESWNSVSEGKQPRFSHRIPLLIFDQDEKGKARDFFTGR  178
gamma-ENaC_Globicephala_melas                TRGSLKTLFGFSEITSRKRREAESWSSARKGTGSKFLNLIPLLAFEKGETSKARDFRTGR  180
gamma-ENaC_Lagenorhynchus_obliquidens        TRGSLKTLFGFSEITSRKRREAESWSSARKGTGSKFLNLIPLLAFEKGETSKARDFRTGR  180
gamma-ENaC_Tursiops_truncatus                TRGSLKTLFGFSEITSRKRREAESWSSARKGTGSKFLNLIPLLAFEKGETSKARDFRTGR  180
gamma-ENaC_Orcinus_orca                      TRGSLKTLFGFSEITSRKRREAESWSSARKGTGSKFLNLIPLLAFEKGETSKARDFRTGR  180
gamma-ENaC_Phocoena_sinus                    TRGSLKTLFGFSEITSRKRREAESWSSARKGTGSKFLNLIPLLAFDKGETSKARDFRTGR  180
gamma-ENaC_Neophocaena_asiaeorientalis       TRGSLKTLFGFSEITSRKRREAESWSSARKGTGSKFLNLIPLLAFDKGETSKARDFRTGR  180
gamma-ENaC_Monodon_monocerons                TRGSLKTLFGFSEITSRKRREAESWSSARKGTGSKFLNLIPLLAFDKGETSKARDFRTGR  180
gamma-ENaC_Delphinapterus_leucas             TRGSLKTLFGFSEITSRKRREAESWSSARKGTGSKFLNLIPLLAFDKGETSKARDFRTGR  180
gamma-ENaC_Pontoporia_blainvillei            TRGSLKTLFGFSEITSRKRREAESWSSAREGTGSKFLNLIPLLAFDKGETSKARDFRTGR  180
gamma-ENaC_Inia_geoffrensis                  TRGNLKTLFGFSEITSRKRREAESWSLAREGTGSKFLNLIPLLAFDKGETSKARDFRTGR  180
gamma-ENaC_Lipotes_vexillifer                TRGSLKTLFGFSEITSRKRREAESWSSAKEGTGSKFLNLIPLLAFDKGETSKARDFRTGR  180
gamma-ENaC_Hyperoodon_ampullatus             TRGTLKTLYGFSEIISRKRREAESWSSAREGTGSKFLNLIPLLAFDKGETSKARDFRTGR  180
gamma-ENaC_Mesoplodon_bidens                 TRGTLKTLYGFSEIISRKRREAESWSSAREGTGSKFLNLIPLLAFDKGETSKARDFRTGR  180
gamma-ENaC_Ziphius_cavirostris               TRGTLKTLYGFSEIISRKRREAESWSSAREGTGSKFLNLIPLLAFDKGETSKARDFRTGR  180
gamma-ENaC_Platanista_gangetica              TRGTLKTLYGFPEITSRKRREAESWSSAREGTGSKFLNLIPLLAFDKGETSKARDFRTGR  180
gamma-ENaC_Platanista_minor                  TRGTLKTLYGFPEITSRKRREAESWSSAREGTGSKFLNLIPLLAFDKGETSKARDFRTGR  180
gamma-ENaC_Kogia_breviceps                   TRGNLKTLYGFSEITSRKRRDAESWSSAREDTGSKFLNLVPLLAFDKGETSKARDFRTGR  180
gamma-ENaC_Physeter_catodon                  TRGTLKTLYGFSEITSRKRREAESWSSAREGTGSTFLNLVPLLAFEKGETSKATDFRTGR  180
gamma-ENaC_Balaenoptera_musculus             TRGTLKTLYGFSEITSRKRREAESWSSAREGTGSKFLNLIPLLAFDKGETSKARDFRTGR  180
gamma-ENaC_Eubalaena_japonica                TRGTLKTLYGFSEITSRKRREAESWSSAREGTGSKFLNLIPLLAFDKGETSKARDFRTGR  180
gamma-ENaC_Hippopotamus_amphibius            TRAALKTLYGFSEITHRKRREAESRSSARE---GKFLNLVPLLTFDTSETGKARDFRTGR  177
gamma-ENaC_Tragulus_javanicus                TRAALKTLYGFEEITSRKCREAESWSSARKDLQPKFLNLAPLMAFEKGETSKARDFFTGQ  180
gamma-ENaC_Tragulus_kanchil                  TRAALKTLYGFEEITSRKRREAESWSSARKDLQPKFLNLAPLMAFEKGETSKARDFFTGQ  180
gamma-ENaC_Antilocapra_americana             TRAALTTLYGFSEITSRKRREAQSWSSAREGTDPRFLNLAPLMAFEKGDKGKARDFFTGR  180
gamma-ENaC_Giraffa_camelopardalis            TRAALKTLYGFSEITSRKRREAQSWSSAREGNDPKFLNLAPLMAFEKGDTGKARDFFTGR  180
gamma-ENaC_Giraffa_tippelskirchi             TRAALKTLYGFSEITSRKRREAQSWSSAREGNDPKFLNLAPLMAFEKGDTGKARDFFTGR  180
gamma-ENaC_Capreolus_pygargus                TRAALKTLYGFSEITSRKRREAQSWSSVRKGTDPKFLNLAPLMAFEKGDTDKARDFFTGR  180
gamma-ENaC_Cervus_elaphus                    TRAALKTLYGFSEIKSRKRREAQPWSSVRKGADPKFLNLAPLMAFEKGDTGKARDFFTGR  180
gamma-ENaC_Moschus_moschiferus               TRAALKTLYGFSEITSRKRREAQSWSSARKGTDPKFLNLAPLMAFEKGDTGKARDFFTGR  180
gamma-ENaC_Moschus_berezovskii               TRAALKTLYGFSEITSRKRREAQSWSSARKGTDPKFLNLAPLMAFEKGDTGKARDFFTGR  180
gamma-ENaC_Bos_grunniens                     TRAALKTLYGFSEITSRKRREAQSWSSVRKGTDPKFLNLAPLMAFEKGDTGKARDFFTGR  180
gamma-ENaC_Bos_taurus                        TRAALKTLYGFSEITSRKRREAQSWSSVRKGTDPKFLNLAPLMAFEKGDTGKARDFFTGR  180
gamma-ENaC_Bubalus_bubalis                   TRAALKTLYGFSEITSRKRREAQSWSSVRRGTDPKFLNLAPLMAFEKGDTGKARDFFTGR  180
gamma-ENaC_Nanger_granti                     TRAALKTLYGFSEITSRKRREAQSQSSVRKDTDPKFLNLAPLMAFEEGDTGKARDFFTGR  180
gamma-ENaC_Kobus_leche                       TRAALKNLYGFSEITSRKRREAQSQSSVRKGTDPKFLNLAPLMAFEKGDTGKARDFITGR  180
gamma-ENaC_Capra_hircus                      TRAALKHLYGFSEITSRKRREAQSRSSVRKGTDPKFLNLAPLMAFEKGDMGKARDFFTGR  180
gamma-ENaC_Ovis_aries                        TRAALKHLYGFSEITSRKRREAQSRSSVRKGTDPRFLNLAPLMALEKGDMGKARDFFTGR  180
gamma-ENaC_Ovis_canadensis                   TRAALKHLYGFSEITSRKRREAQSRSSVRKGTDPRFLNLAPLMAFEKGDMGKARDFFTGR  180
gamma-ENaC_Oreamnos_americanus               TRAALKHLYGFSEITSRKRREAQSRSSVRKGTDPKFLNLAPLMAFEKGDMGKARDFFTGR  180
gamma-ENaC_Hippotragus_niger                 TRAALKTLYGFSEITSRKRREAQSRSSVRKGTDPKFLNLAPLMAFEKGDTGKARDFFTGR  180
gamma-ENaC_Damaliscus_lunatus                TREALKTLYGFSEITSRKRREAQSRSSVRKSTDPKFLNLAPLMAFEQGDTGKARDFFTGR  180
gamma-ENaC_Sus_scrofa                        TRGALKTLYGFSEITSRKRREAESQNSAWEGTGSKFLNLIPLLVFNQGETGKARDFLTGR  180
gamma-ENaC_Vicugna_pacos                     TRAALKTLYGFPEITSRKRRESESWRSASEGSGSKFLNLVPLLAFNQSERNKARDFLTGR  180
gamma-ENaC_Equus_callabus                    TRGALKTLYGFSEIKSRKRRDAESWSSVWEGTRPKFLNLVPLLAFKEGEMGKARDFLTGR  180
                                             **  *. *:** *   ** *:::    . .     * :  **: :. .: .** ** **:

gamma-ENaC_Homo_sapiens                      KRKVGGSIIHKASNVMHIE-SKQVVGFQLCSNDTSDCATYTFSSGINAIQEWYKLHYMNI  237
gamma-ENaC_Globicephala_melas                KRKVSGRIVHTASDVVHVYESKGLVGFQLCSNDTSSCAVYTFTSGVNAIREWYKLHYMNI  240
gamma-ENaC_Lagenorhynchus_obliquidens        KRKVSGRIVHTASDVVHVYESKGLVGFQLCSNDTSSCAVYTFTSGVNAIREWYKLHYMNI  240
gamma-ENaC_Tursiops_truncatus                KRKVSGRIVHTASDVVHVYESKGLVGFQLCSNDTSSCAVYTFTSGVNAIREWYKLHYMNI  240
gamma-ENaC_Orcinus_orca                      KRKVSGRIVHTASDVVHVYESKGLVGFQLCSNDTSSCAVYTFTSGVNAIREWYKLHYMNI  240
gamma-ENaC_Phocoena_sinus                    KRKVSGRIIHTASDVVHVYESKGSVGFQLCSNDTSSCAVYTFTSGVNAIREWYKLHYMNI  240
gamma-ENaC_Neophocaena_asiaeorientalis       KRKVSGRIIHTASDVVHVYESKGSVGFQLCSNDTSSCAVYTFTSGVNAIREWYKLHYMNI  240
gamma-ENaC_Monodon_monocerons                KRKVSGRIIHTASDVVHVYESKGLVGFQLCSNDTSSCAVYTFTSGVNAIREWYKLHYMNI  240
gamma-ENaC_Delphinapterus_leucas             KRKVSGRIIHTASDVVHVYESKGLVGFQLCSNDTSSCAVYTFTSGVNAIREWYKLHYMNI  240
gamma-ENaC_Pontoporia_blainvillei            KRQVSGRIIHKASDVVHVYESKDVVGFQLCLNDTSNCTVYTFSSGVNAIREWYKLHYMNI  240
gamma-ENaC_Inia_geoffrensis                  KRKVSGRIIHKASDVMHVYESKDVVGFQLCSNDTSSCAVYTFSSGVNAIREWYKLHYMNI  240
```

```
gamma-ENaC_Lipotes_vexillifer               KRKVNGRIIHKASDVVHVYESKDMVGFQLCSNDTSSCAVYTFSSGINAIREWYKLHYMNI  240
gamma-ENaC_Hyperoodon_ampullatus            KRKVRGRIIHKASDVMHVHDSKEVVGFQLCPNDTSSCTVYTFNSGVNAIREWYKLHYMNI  240
gamma-ENaC_Mesoplodon_bidens                KRKVRGRIIHKASDVMHVHDSKEVVGFQLCPNDTSSCTVYTFNSGVNAIREWYKLHYMNI  240
gamma-ENaC_Ziphius_cavirostris              KRKVRGKIIHKASDVMHVHDSKEVVGFQLCPNDTSRCTVYTFNSGVNAIREWYKLHYMNI  240
gamma-ENaC_Platanista_gangetica             KRKVSGRIIHKASDVMHVHESKEVVGFQLCSNDTSSCAVYTFSSGVNAIREWYKLHYMNI  240
gamma-ENaC_Platanista_minor                 KRKVSGRIIHKASDVMHVHESKEVVGFQLCSNDTSSCAVYTFSSGVNAIREWYKLHYMNI  240
gamma-ENaC_Kogia_breviceps                  KRKVSGRIIHEASDVMHVHES-EVVGFQLCSNDTSSCAVYTFSSGVNAIQEWYKLHYMNI  239
gamma-ENaC_Physeter_catodon                 KRKVSGRIIHKSSDVMHVHESKEVVGFQLCSNDTSSCAVYTFSSGVNAIQEWYKLHYMNI  240
gamma-ENaC_Balaenoptera_musculus            KRKVSGRIIHKASDVMHVHESKEVVGFQLCSNDTSSCAVYTFSSGVNAIREWYKLHYMNI  240
gamma-ENaC_Eubalaena_japonica               KRKVSGRIIHEASDVMHVHESKEVVGFQLCSNDTSSCAVYTFSSGVNAIREWYKLHYMNI  240
gamma-ENaC_Hippopotamus_amphibius           KRKVNGKIIHKASDVMHVHDSKEVVGFQLCSNDTSSCAVYTFSSGINAIQEWYKLHYMNI  237
gamma-ENaC_Tragulus_javanicus               KRRVRGRIVHKASDVMHIHNSKEVVGFQLCSNDTSDCAVYTFSSGVNAIQEWYKLHYMNI  240
gamma-ENaC_Tragulus_kanchil                 KRRVRGRIVHKASDVMHIHNSKEVVGFQLCSNDTSDCAVYTFSSGVNAIQEWYKLHYMNI  240
gamma-ENaC_Antilocapra_americana            KRKVNARIIHKASDVMHIHSSKEVVGFQLCSNDTSDCAVYTFSSGINAIQEWYKLHYMNI  240
gamma-ENaC_Giraffa_camelopardalis           KRKVNARIIHKASDVMHIHNSKEVVGFQLCSNDTSDCAVYTFSSGVNAIQEWYKLHYMNI  240
gamma-ENaC_Giraffa_tippelskirchi            KRKVNARIIHKASDVMHIHNSKEVVGFQLCSNDTSDCAVYTFSSGVNAIQEWYKLHYMNI  240
gamma-ENaC_Capreolus_pygargus               KRKVNARIIHKASDVMHIHNSKEVVGFQLCSNDTSDCAVYTFSSGVNAIQEWYKLHYMNI  240
gamma-ENaC_Cervus_elaphus                   KRRVNAKIIHKASDVMHIHNSKEVVGFQLCSNDTSDCAVYTFSSGVNAIQEWYKLHYMNI  240
gamma-ENaC_Moschus_moschiferus              KRRVNARIIHKASDVMHIHNSKEVVGFQLCSNDTSDCAVYTFSSGVNAIQEWYKLHYMNI  240
gamma-ENaC_Moschus_berezovskii              KRRVNARIIHKASDVMHIHNSKEVVGFQLCSNDTSDCAVYTFSSGVNAIQEWYKLHYMNI  240
gamma-ENaC_Bos_grunniens                    KRKVNARIIHKASDVMHIHNSKEVVGFQLCSNDTSDCAVYTFSSGVNAIQEWYKLHYMNI  240
gamma-ENaC_Bos_taurus                       KRKVNARIIHKASDVMHIHNSKEVVGFQLCSNDTSDCAVYTFSSGVNAIQEWYKLHYMNI  240
gamma-ENaC_Bubalus_bubalis                  KRKVNARIIHKASDVMHIHNSKEVVGFQLCSNDTSDCAVYTFSSGVNAIQEWYKLHYMNI  240
gamma-ENaC_Nanger_granti                    KRKVNAKIIHKASDVMHIHNSKEVVGFQLCSNDTSDCAVYTFSSGVNAIQEWYKLHYMNI  240
gamma-ENaC_Kobus_leche                      KRKVNARIIHKASDVMHIHNSKEVVGFQLCSNDTSDCAVYTFSSGVNAIQEWYKLHYMNI  240
gamma-ENaC_Capra_hircus                     KRKVNARIIHKASDVMHIHNSKEVVGFQLCSNDTSDCAVYTFSSGVNAIQEWYKLHYMNI  240
gamma-ENaC_Ovis_aries                       KRKVNARIIHKASDVMHIHNSKEVVGFQLCSNDTSDCAVYTFSSGVNAIQEWYKLHYMNI  240
gamma-ENaC_Ovis_canadensis                  KRKVNARIIHKASDVMHIHNSKEVVGFQLCSNDTSDCAVYTFSSGVNAIQEWYKLHYMNI  240
gamma-ENaC_Oreamnos_americanus              KRKVNARIIHKASDVMHIHNSKEVVGFQLCSNDTSDCAVYTFSSGVNAIQEWYKLHYMNI  240
gamma-ENaC_Hippotragus_niger                KRKVNARIIHKASDVMHIHNSKEVVGFQLCSNDTSDCAVYTFSSGVNAIQEWYKLHYMNI  240
gamma-ENaC_Damaliscus_lunatus               KRKVNARIIHKASDVMHIHNSKEVVGFQLCSNDTSDCAVYTFSSGVNAIQEWYKLHYMNI  240
gamma-ENaC_Sus_scrofa                       KRKVSGSIIHKASDVMHVHDSKEVVGFQLCANDTSNCAVYTFSSGVNAIQEWYKLHYMNI  240
gamma-ENaC_Vicugna_pacos                    KRKVSGSIIHKASDVMHGHESKEVVGFQLCSNDTSDCAVYTFSSGVNAIREWYKLHYMNI  240
gamma-ENaC_Equus_callabus                   KRKVSGSIIRKESDVMNVHESKEVVGFQLCSNDTSNCAVYTFSSGVNAIQEWYKLHYMNI  240
                                            **:* . *::  *:*::   *   ****** **** *:.***.**:***:**********

gamma-ENaC_Homo_sapiens                     MAQVPLEKKINMSYSAEELLVTCFFDGVSCDARNFTLFHHPMHGNCYTFNNRENETILST  297
gamma-ENaC_Globicephala_melas               MAQVSLEKKINMSYSAEELLVTCFFDGESCDARNFTRFHHPMYGNCYTFNNRQNETILST  300
gamma-ENaC_Lagenorhynchus_obliquidens       MAQVSLEKKINMSYSAEELLVTCFFDGESCDARNFTRFHHPMYGNCYTFNNRQNETILST  300
gamma-ENaC_Tursiops_truncatus               MAQVSLEKKINMSYSAEELLVTCFFDGESCDARNFTRFHHPMYGNCYTFNNRQNETILST  300
gamma-ENaC_Orcinus_orca                     MAQVSLEKKINMSYSAEELLVTCFFDGESCDARNFTRFHHPMYGNCYTFNNRQNETILST  300
gamma-ENaC_Phocoena_sinus                   MAQVSLEKKINMSYSAEELLVTCFFDGESCDARNFTRFHHPMYGNCYTFNNRQNETILST  300
gamma-ENaC_Neophocaena_asiaeorientalis      MAQVSLEKKINMSYSAEELLVTCFFDGESCDARNFTRFHHPMYGNCYTFNNRQNETILST  300
gamma-ENaC_Monodon_monocerons               MAQVSLEKKINMSYSAEELLVTCFFDGESCDARNFTRFHHPMYGNCYTFNNRQNETILST  300
gamma-ENaC_Delphinapterus_leucas            MAQVSLEKKINMSYSAEELLVTCFFDGESCDARNFTRFHHPMYGNCYTFNNRQNETILST  300
gamma-ENaC_Pontoporia_blainvillei           MAQVSLEKKINMSYSAEELLVTCFFDGESCDARNFTRFHHPMYGNCYTFNNRQNETILST  300
gamma-ENaC_Inia_geoffrensis                 MAQVSLEKKINMSYSAEELLVTCFFDGESCDARNFTRFHHPMYGNCYTFNNRQNETILST  300
gamma-ENaC_Lipotes_vexillifer               MAQVSLEKKINMSYSAEELLVTCFFDGESCDARNFTRFHHPMYGNCYTFNNRQNETVLST  300
gamma-ENaC_Hyperoodon_ampullatus            MARVSPEKKINMSYSAEELLVTCFFDGESCDARNFTRFHHPMYGNCYTFNNGQNETMLST  300
gamma-ENaC_Mesoplodon_bidens                MARVSPEKKINMSYSAEELLVTCFFDGESCDARNFTRFHHPMYGNCYTFNNGQNETMLST  300
gamma-ENaC_Ziphius_cavirostris              MARVSPEKKINMSYSAEELLVTCFFDGESCDARNFTRFHHPMYGNCYTFNNGQNETMLST  300
gamma-ENaC_Platanista_gangetica             MAQVSPEKKINMSYSAEELLVTCFFDGESCDARNFTRFHHPMYGNCYTFNNRQNETILST  300
gamma-ENaC_Platanista_minor                 MAQVSPEKKINMSYSAEELLVTCFFDGESCDARNFTRFHHPMYGNCYTFNNRQNETILST  300
gamma-ENaC_Kogia_breviceps                  MAQVSPEKKINMSYSAEELLVTCFFDGESCDARNFTRFHHPMYGNCYTFNNRENETILST  299
gamma-ENaC_Physeter_catodon                 MAQVSPEKKINMSYSAEELLVTCFFDGESCDARNFTRFHHPMYGNCYTFNKGENETILST  300
gamma-ENaC_Balaenoptera_musculus            MAQVSPEKKINMSYSAEELLVTCFFDGESCDARNFTRFHHPMYGNCYTFNNGQNETILST  300
gamma-ENaC_Eubalaena_japonica               MAQVPPEKKINMSYSAEELLVTCFFDGESCDARNFTRFHHPMYGNCYTFNNGQNETILST  300
gamma-ENaC_Hippopotamus_amphibius           MAQVSLEKKINMSYSAEELLVTCFFDGVSCDARNFTRSHHPMYGNCYTFNNRQNETILST  297
gamma-ENaC_Tragulus_javanicus               MAQVSREKKINMSYSAEELLITCFFDGMSCDARNFTLFHHPMYGNCYTFNNRQNETILST  300
gamma-ENaC_Tragulus_kanchil                 MAQVSREKKINMSYSAEELLITCFFDGMSCDARNFTLFHHPMYGNCYTFNNRQNETILST  300
gamma-ENaC_Antilocapra_americana            MAQVSQEKKINMSYSAEELLVTCFFDGVSCDARNFTLFHHPMYGNCYTFNNRQNETILST  300
gamma-ENaC_Giraffa_camelopardalis           MAQVSQEKKINMSYSADELLVTCFFDGVSCDARNFTLFHHPMYGNCYTFNNRQNETILST  300
gamma-ENaC_Giraffa_tippelskirchi            MAQVSQEKKINMSYSADELLVTCFFDGVSCDARNFTLFHHPMYGNCYTFNNRQNETILST  300
gamma-ENaC_Capreolus_pygargus               MAQVSQEKKINMSYSADELLITCFFDGVSCDARNFTLFHHPMYGNCYTFNNRQNETILST  300
gamma-ENaC_Cervus_elaphus                   MAQVSQEKKINMSYSADELLITCFFDGVSCDARNFTLFHHPMYGNCYTFNNRQNETTLST  300
gamma-ENaC_Moschus_moschiferus              MAQVSQEKKINMSYSADELLVTCFFDGVSCDARNFTLFHHPMYGNCYTFNNRQNETILST  300
gamma-ENaC_Moschus_berezovskii              MAQVSQEKKINMSYSADELLVTCFFDGVSCDARNFTLFHHPMYGNCYTFNNRQNETILST  300
gamma-ENaC_Bos_grunniens                    MAQVSQEKKINMSYSADELLVTCFFDGVSCDARNFTLFHHPMYGNCYTFNNRQNETILST  300
gamma-ENaC_Bos_taurus                       MAQVSQEKKINMSYSADELLVTCFFDGVSCDARNFTLFHHPMYGNCYTFNNRQNETILST  300
gamma-ENaC_Bubalus_bubalis                  MAQVSQEKKINMSYSADELLVTCFFDGVSCDARNFTLFHHPMYGNCYTFNNRQNETILST  300
gamma-ENaC_Nanger_granti                    MAQVSQEKKINMSYSADELLVTCFFDGMSCDARNFTLFHHPMYGNCYTFNNRQNETILST  300
gamma-ENaC_Kobus_leche                      MAQVSQEKKINMSYSADELLVTCFFDGVSCDARNFTLFHHPMYGNCYTFNNRQNETILST  300
gamma-ENaC_Capra_hircus                     MAQVSQEKKINMSYSADELLVTCFFDGVSCDARNFTLFHHPMYGNCYTFNNRQNETILST  300
gamma-ENaC_Ovis_aries                       MAQVSQEKKINMSYSADELLVTCFFDGVSCDARNFTLFHHPMYGNCYTFNNRQNETILST  300
gamma-ENaC_Ovis_canadensis                  MAQVSQEKKINMSYSADELLVTCFFDGVSCDARNFTLFHHPMYGNCYTFNNRQNETILST  300
gamma-ENaC_Oreamnos_americanus              MAQVSQEKKINMSYSADELLVTCFFDGVSCDARNFTLFHHPMYGNCYTFNNRQNETILST  300
gamma-ENaC_Hippotragus_niger                MAQVSQEKKINMSYSADELLVTCFFDGVSCDARNFTLFHHPMYGNCYTFNNRQNETILST  300
gamma-ENaC_Damaliscus_lunatus               MAQVSQEKKINMSYSADELLVTCFFDGVSCDARNFTLFHHPMYGNCYTFNNRQNETILST  300
gamma-ENaC_Sus_scrofa                       MAQVPLEKKINMSYSAEELLVTCFFDGVSCDARNFTLFHHPMYGNCYTFNNRENETTLST  300
gamma-ENaC_Vicugna_pacos                    MAQVSLEKKINMSYSAEELLVNCFFDGVSCDARNFTLFHHPMYGNCYTFNNRQNETVLST  300
gamma-ENaC_Equus_callabus                   MAQVSLEKKINMSYSAEELLVTCFFDGVSCDARNFTLFHHPMYGNCYTFNNRQNETILST  300
                                            **:*  **********:***:.***** ********  ****:*******: :*** ***

gamma-ENaC_Homo_sapiens                     SMGGSEYGLQVILYINEEEYNPFLVSSTGAKVIIHRQDEYPFVEDVGTEIETAMVTSIGM  357
```

```
gamma-ENaC_Globicephala_melas                SMGGSEYGLQVILYLNEEEYNPFLVSSTGAKVIVHRQFEYPFIEDVGTEIETAMATSIGM  360
gamma-ENaC_Lagenorhynchus_obliquidens        SMGGSEYGLQVILYLNEEEYNPFLVSSTGAKVIVHRQFEYPFIEDVGTEIETAMATSIGM  360
gamma-ENaC_Tursiops_truncatus                SMGGSEYGLQVILYLNEEEYNPFLVSSTGAKVIVHRQFEYPFIEDVGTEIETAMATSIGM  360
gamma-ENaC_Orcinus_orca                      SMGGSEYGLQVILYLNEEEYNPFLVSSTGAKVIVHRQFEYPFIEDVGTEIETAMATSIGM  360
gamma-ENaC_Phocoena_sinus                    SMGGSEYGLQVILYLNEEEYNPFLVSSTGAKVIVHRQYEYPFIEDVGTEIETAMATSIGM  360
gamma-ENaC_Neophocaena_asiaeorientalis       SMGGSEYGLQVILYLNEEEYNPFLVSSTGAKVIVHRQYEYPFIEDVGTEIETAMATSIGM  360
gamma-ENaC_Monodon_monocerons                SMGGSEYGLQVILYLNEEEYNPFLVSSTGAKVIVHRQYEYPFIEDVGTEIETAMATSIGM  360
gamma-ENaC_Delphinapterus_leucas             SMGGSEYGLQVILYLNEEEYNPFLVSSTGAKVIVHRQYEYPFIEDVGTEIETAMATSIGM  360
gamma-ENaC_Pontoporia_blainvillei            SMGGSEYGLQVILYINEEEYNPFLVSSTGAKVIVHRQDEYPFIEDVGTEIETAMATSIGM  360
gamma-ENaC_Inia_geoffrensis                  SMGGSEYGLQVILYINEEEYNPFLVSSTGAKVIVHRQDEYPFIEDVGTEIETAMATSIGM  360
gamma-ENaC_Lipotes_vexillifer                SVGGSEYGLQVILYINEEEYNPFLVSSTGAKVVVHRQDEYPFIEDVGTEIETAMATSIGM  360
gamma-ENaC_Hyperoodon_ampullatus             SMGGSDYGLQVILYIDEEEYNPFLVSSTGAKVIVHRQDEYPFIEDVGTEIETAMATSIGM  360
gamma-ENaC_Mesoplodon_bidens                 SMGGSDYGLQVILYIDEEEYNPFLVSSTGAKVIVHRQDEYPFIEDVGTEIETAMATSIGM  360
gamma-ENaC_Ziphius_cavirostris               SMGGSDYGLQVILYIDEEEYNPFLVSSTGAKVIVHRQDEYPFIEDVGTEIETAMATSIGM  360
gamma-ENaC_Platanista_gangetica              SMGGSEYGLQVILYINEEEYNPFLVSSTGAKVIVHQQDEYPFIEDVGTEIETAMATSIGM  360
gamma-ENaC_Platanista_minor                  SMGGSEYGLQVILYINEEEYNPFLVSSTGAKVIVHQQDEYPFIEDVGTEIETAMATSIGM  360
gamma-ENaC_Kogia_breviceps                   SMGGSEYGLQVILYINEEEYNPFLVSSTGAKVIVHRQDEYPFIEDVGTEIETAMATSIGM  359
gamma-ENaC_Physeter_catodon                  SMGGSEYGLQVILYINEEEYNPFLVSSTGAKVIVHRQDEYPFIEDVGTEIETAMATSIGM  360
gamma-ENaC_Balaenoptera_musculus             SMGGSEYGLQVILYINEEEYNPFLVSSTGAKVIVHRQDEYPFIEDVGTEIETAMATSIGM  360
gamma-ENaC_Eubalaena_japonica                SMGGSEYGLQVILYINEEEYNPFLVSSTGAKVIVHRQDEYPFIEDVGTEIETAMATSIGM  360
gamma-ENaC_Hippopotamus_amphibius            SMGGSEYGLQVILYINEEEYNPFLVSSTGAKVIVHRQDQYPFIEDMGTEIETAMATSIGM  357
gamma-ENaC_Tragulus_javanicus                SMGGSEFGLQVILYINEEEYNPFLVSSTGAKVIVHRQDEYPFVEEMGTEIETAMATSIGM  360
gamma-ENaC_Tragulus_kanchil                  SMGGSEFGLQVILYINEEEYNPFLVSSTGAKVIVHRQDEYPFVEEMGTEIETAMATSIGM  360
gamma-ENaC_Antilocapra_americana             SMGGSEFGLQVILYINEEEYNPFLVSSTGAKVIVHRQDEYPFVEDVGTEIETAMATSIGM  360
gamma-ENaC_Giraffa_camelopardalis            SMGGSEFGLQVILYVNEEEYNPFLVSSTGAKVIVHRQDEYPFVEDMGTEIETAMATSIGM  360
gamma-ENaC_Giraffa_tippelskirchi             SMGGSEFGLQVILYVNEEEYNPFLVSSTGAKVIVHRQDEYPFVEDMGTEIETAMATSIGM  360
gamma-ENaC_Capreolus_pygargus                SMGGSEFGLQVILYINEEEYNPFLVSSTGAKVIVHRQDEYPFAEDVGTEIETAMATSIGM  360
gamma-ENaC_Cervus_elaphus                    SMGGSEFGLQVILYINEEEYNPFLVSSTGAKVIVHRQDEYPFVEDVGTEIETAMATSIGM  360
gamma-ENaC_Moschus_moschiferus               SMGGSEFGLQVILYINEEEYNPFLVSSTGAKVIVHRQDEYPFVEDVGTEIETAMATSIGM  360
gamma-ENaC_Moschus_berezovskii               SMGGSEFGLQVILYINEEEYNPFLVSSTGAKVIVHRQDEYPFVEDVGTEIETAMATSIGM  360
gamma-ENaC_Bos_grunniens                     SMGGSEFGLQVILYINEEEYNPFLVSSTGAKVIIHRQDEYPFVEDVGTEIETAMATSIGM  360
gamma-ENaC_Bos_taurus                        SMGGSEFGLQVILYINEEEYNPFLVSSTGAKVIIHRQDEYPFVEDVGTEIETAMATSIGM  360
gamma-ENaC_Bubalus_bubalis                   SMGGSEFGLQVILYINEEEYNPFLVSSTGAKVIVHRQDEYPFIEDVGTEIETAMATSIGM  360
gamma-ENaC_Nanger_granti                     SMGGSEFGLQVILYINEEEYNPFLVSSTGAKVIVHRQDEYPFVEDVGTEIETAMATSIGM  360
gamma-ENaC_Kobus_leche                       SMGGSEFGLQVILYINEEEYNPFLVSSTGAKVIVHRQDEYPFVENMGTEIETAMATSIGM  360
gamma-ENaC_Capra_hircus                      SMGGSEFGLQVILYINEEEYNPFLVSSTGAKVIVHRQDEYPFIEDMGTEIETAMATSIGM  360
gamma-ENaC_Ovis_aries                        SMGGSEFGLQVILYINEEEYNPFLVSSTGAKVIVHRQDEYPFIEDMGTEIETAMATSIGM  360
gamma-ENaC_Ovis_canadensis                   SMGGSEFGLQVILYINEEEYNPFLVSSTGAKVIVHRQDEYPFIEDMGTEIETAMATSIGM  360
gamma-ENaC_Oreamnos_americanus               SMGGSEFGLQVILYINEEEYNPFLVSSTGAKVIVHRQDEYPFVEDVGTEIETAMATSIGM  360
gamma-ENaC_Hippotragus_niger                 SMGGSEFGLQVILYINEEEYNPFLVSSTGAKVIVHRQDEYPFVEDMGTEIETAMATSIGM  360
gamma-ENaC_Damaliscus_lunatus                SMGGSEFGLQVILYINEEEYNPFLVSSTGAKVIVHRQDEYPFVEDAGTEIETAMATSIGM  360
gamma-ENaC_Sus_scrofa                        SMGGSEYGLQVILYVNEEEYNPFLVSSTGAKVIVHRQDEYPFIEDVGTEIETAMATSIGM  360
gamma-ENaC_Vicugna_pacos                     SMGGSEYGLQVILHINEEEYNPFLVSSTGAKVIVHRQDEYPFIEDVGTEIETAMATSIGM  360
gamma-ENaC_Equus_callabus                    SMGGSEYGLQVILHINEEEYNPFLVSSTGAKVIIHRQDEYPFVEDVGTEIETAMATSIGM  360
                                             *:***::******:::****************::*:* :*** *: ********.*****

gamma-ENaC_Homo_sapiens                      HLTESFKLSEPYSQCTEDGSDVPIRNIYNAAYSLQICLHSCFQTKMVEKCGCAQYSQPLP  417
gamma-ENaC_Globicephala_melas                HLTESFKLSEPYSRCTEDWNGVLITNIYNATYSLQICLHSCFQAKMVEKCGCAQFSQPLP  420
gamma-ENaC_Lagenorhynchus_obliquidens        HLTESFKLSEPYSRCTEDWNGVLITNIYNATYSLQICLHSCFQAKMVEKCGCAQFSQPLP  420
gamma-ENaC_Tursiops_truncatus                HLTESFKLSEPYSRCTEDWNGVLITNIYNATYSLQICLHSCFQAKMVEKCGCAQFSQPLP  420
gamma-ENaC_Orcinus_orca                      HLTESFKLSEPYSRCTEDWNGVLITNIYNATYSLQICLHSCFQAKMVEKCGCAQFSQPLP  420
gamma-ENaC_Phocoena_sinus                    HLTESFKLSEPYSRCTEDWNGVLITNIYNATYSLQICLHSCFQAKMVEKCGCAQFSQPLP  420
gamma-ENaC_Neophocaena_asiaeorientalis       HLTESFKLSEPYSRCTEDWNGVLITNIYNATYSLQICLHSCFQAKMVEKCGCAQFSQPLP  420
gamma-ENaC_Monodon_monocerons                HLTESFKLSEPYSRCTEDWNGVLITNIYNATYSLQICLHSCFQAKMVEKCGCAQFSQPLP  420
gamma-ENaC_Delphinapterus_leucas             HLTESFKLSEPYSRCTEDWNGVLITNIYNATYSLQICLHSCFQAKMVEKCGCAQFSQPLP  420
gamma-ENaC_Pontoporia_blainvillei            HLTESFKLSEPYSQCTEDWSDVLITNIYNATYSLQICLHSCFQAKMVEKCGCAQFSQPLP  420
gamma-ENaC_Inia_geoffrensis                  HLTESFKLSEPYSQCTEDWSDVLITNIYNATYSLQICLHSCFQAKMVEKCGCAQFSQPLP  420
gamma-ENaC_Lipotes_vexillifer                HLTESFKLSEPYSQCKEDWSNVLITNIYNAPYSLQICLHSCFQAKMVEKCGCAQFSQPLP  420
gamma-ENaC_Hyperoodon_ampullatus             HLTESFKLSDPYSRCMEDWSDVLITNIYNATYSLRICLHSCFQAKMVEKCGCAQFSQPLP  420
gamma-ENaC_Mesoplodon_bidens                 HLTESFKLSDPYSRCTEDWSDVLITNIYNATYSLRICLHSCFQAKMVEKCGCAQFSQPLP  420
gamma-ENaC_Ziphius_cavirostris               HLTESFKLSDPYSRCMEDWSDVLITNIYNATYSLRICLHSCFQAKMVEKCGCAQFSQPLP  420
gamma-ENaC_Platanista_gangetica              HLTESFKLSEPYSQCTEDWSDVLITNIYNATYSLQICLHSCFQAKMVEKCGCAQFSQPLP  420
gamma-ENaC_Platanista_minor                  HLTESFKLSEPYSQCTEDWSDVLITNIYNATYSLQICLHSCFQAKMVEKCGCAQFSQPLP  420
gamma-ENaC_Kogia_breviceps                   HLTESFKLSEPYSQCTEDWSDVLITNIYNATYSLQICLHSCFQAKMVEKCGCAQFSQPLP  419
gamma-ENaC_Physeter_catodon                  HLTESFKLSEPYSRCTEDWSDVLIKNIYNATYSLQICLHSCFQAKMVEKCGCAQFNQPLP  420
gamma-ENaC_Balaenoptera_musculus             HLTESFKLSEPYSKCTEDWSDVLITNIYNATYSLQICLHSCFQAKMVEKCGCAQFSQPLP  420
gamma-ENaC_Eubalaena_japonica                HLTESFKLSEPYSKCTEDWSDVLITNIYNATYSLQICLHSCFQAKMVEKCGCAQFSQPLP  420
gamma-ENaC_Hippopotamus_amphibius            HLTESFKLSEPYSQCTEDWRDVPVMNIYNATYSLQICLHSCFQAKMVENCGCAQYSQPLP  417
gamma-ENaC_Tragulus_javanicus                HLTESFKLSDPYSQCTEDWGDVQIKNIYNATYSLQICLHSCFQTKMVEKCGCAQYSQPLP  420
gamma-ENaC_Tragulus_kanchil                  HLTESFKLSDPYSQCTEDWGDVQIKNIYNATYSLQICLHSCFQTKMVEKCGCAQYSQPLP  420
gamma-ENaC_Antilocapra_americana             HLTESFKLSDPYSQCTEDWSDVQIRNIYNATYSLQICLHSCFQAKMVENCGCAQYSQPLP  420
gamma-ENaC_Giraffa_camelopardalis            HLTESFKLSDPYSQCTEDWSDVQITNIYNATYSLQICLHSCFQAKMVENCGCAQYSQPLP  420
gamma-ENaC_Giraffa_tippelskirchi             HLTESFKLSDPYSQCTEDWSDVQITNIYNATYSLQICLHSCFQAKMVENCGCAQYSQPLP  420
gamma-ENaC_Capreolus_pygargus                HLTESFKLSDPYSQCTEDWSDVQITNIYNATYSSQICLHSCFQAKMVENCGCAQYSQPLP  420
gamma-ENaC_Cervus_elaphus                    HLTESFKLSDPYSQCTEDWSDVQITNIYNATYSPQICLHSCFQAKMVENCGCAQYSQPLP  420
gamma-ENaC_Moschus_moschiferus               HLTESFKLSDPYSQCTEDWSDVQITNIYNATYSLQICLHSCFQAKMVENCGCAQYSQPLP  420
gamma-ENaC_Moschus_berezovskii               HLTESFKLSDPYSQCTEDWSDVQITNIYNATYSLQICLHSCFQAKMVENCGCAQYSQPLP  420
gamma-ENaC_Bos_grunniens                     HLTESFKLSDPYSQCTEDWSDVQITNIYNATYSLQICLHSCFQAKMVENCGCAQYSQPLP  420
gamma-ENaC_Bos_taurus                        HLTESFKLSDPYSQCTEDWSDVQITNIYNATYSLQICLHSCFQAKMVENCGCAQYSQPLP  420
gamma-ENaC_Bubalus_bubalis                   HLTESFKLSDPYSHCTEDWSDVQITNIYNATYSLQICLHSCFQAKMVENCGCAQYSQPLP  420
gamma-ENaC_Nanger_granti                     HLTESFKLSDPYSHCTEDWSDVQITNIYNATYSLQICLHSCFQAKMVENCGCAQYSQPLP  420
gamma-ENaC_Kobus_leche                       HLTESFKLSDPYSHCTEDWSDVQITNIYNATYSLQICLHSCFQAKMVENCGCAQYSQPLP  420
gamma-ENaC_Capra_hircus                      HLTESFKLSDPYSRCTEDWSDVQITNIYNATYSLQICLHSCFQAKMVENCGCAQYSQPLP  420
gamma-ENaC_Ovis_aries                        HLTESFKLSDPYSRCTEDWSDVQITNIFNATYSLQICLHSCFQAKMVENCGCAQYSQPLP  420
```

```
gamma-ENaC_Ovis_canadensis                  HLTESFKLSDPYSRCTEDWSDVQITNIFNATYSLQICLHSCFQAKMVENCGCAQYSQPLP  420
gamma-ENaC_Oreamnos_americanus              HLTESFKLSDPYSHCTEDWSDVQITNIYNATYSLQICLHSCFQAKMVENCGCAQYSQPLP  420
gamma-ENaC_Hippotragus_niger                HLTESFKLSDPYSHCTEDWSDVQITNIYNATYSLQICLHSCFQAKMVENCGCAQYSQPLP  420
gamma-ENaC_Damaliscus_lunatus               HLTESFKLSDPYSHCTEDWSDVQITNIYNATYSLQICLHSCFQAKMVENCGCAQYSQPLP  420
gamma-ENaC_Sus_scrofa                       HLTESFKLGEPYSQCTEDGSDVPVENIYGAAYSLQICLNSCFQAKMVEKCGCAQYSKPLP  420
gamma-ENaC_Vicugna_pacos                    HLTESFKLSEPYSQCTEDGSEVPIQNIYNASYSLQICLHSCFQAKMVEKCGCAQYSKPLP  420
gamma-ENaC_Equus_callabus                   HLTESFKLSEPYSQCTEDGSDVPVENIYKAAYSLKICLHSCFQTKMVEKCGCAQYSQPLP  420
                                            ********.:***:* **   * : **: * ** :***:****:****:*****:.:***

gamma-ENaC_Homo_sapiens                     PAANYCNYQQHPNWMYCYYQLHRAFVQEELGCQSVCKEACSFKEWTLTTSLAQWPSVVSE  477
gamma-ENaC_Globicephala_melas               QGANYCNYRQHPNWMYCYYELHQAFVWGELGCQAMCKEACSFKEWTLTTSLAQWPSEVSE  480
gamma-ENaC_Lagenorhynchus_obliquidens       QGANYCNYRQHPNWMYCYYELHQAFVWGELGCQSMCKEACSFKEWTLTTSLAQWPSEVSE  480
gamma-ENaC_Tursiops_truncatus               QGANYCNYRQHPNWMYCYYELHQAFVWGELGCQSMCKEACSFKEWTLTTSLAQWPSEVSE  480
gamma-ENaC_Orcinus_orca                     QGANYCNYRQHPNWMYCYYELHQAFVWGELGCQSMCKEACSFKEWTLTTSLAQWPSEVSE  480
gamma-ENaC_Phocoena_sinus                   QGANYCNYRQHPNWMYCYYELHQAFVWGELGCQSVCKEACSFKEWTLTTSLAQWPSEVSE  480
gamma-ENaC_Neophocaena_asiaeorientalis      QGANYCNYRQHPNWMYCYYELHQAFVWGELGCQSVCKEACSFKEWTLTTSLAQWPSEVSE  480
gamma-ENaC_Monodon_monocerons               QGANYCNYRQHPNWMYCYYELHQAFVWGELGCQSVCKEACSFKEWTLTTSLAQWPSEVSE  480
gamma-ENaC_Delphinapterus_leucas            QGANYCNYRQHPNWMYCYYELHQAFVWGELGCQSVCKEACSFKEWTLTTSLAQWPSEVSE  480
gamma-ENaC_Pontoporia_blainvillei           RGASYCNYRQHPNWMYCYYRLHQAFVWGELGCQSVCKEACSFKEWTLTTSLAQWPSEVSE  480
gamma-ENaC_Inia_geoffrensis                 RGASYCNYRQHPNWMYCYYELHQAFVWGELGCQSVCKEACSFKEWTLTTSLAQWPSEVSE  480
gamma-ENaC_Lipotes_vexillifer               QGANYCNYRQHPNWMYCYYELHQAFVWGELGCQSVCKEACSFKEWTLTTSLAQWPSEVSE  480
gamma-ENaC_Hyperoodon_ampullatus            QGANYCNYQQHPNWMYCYYELHKDFVRGELGCQSVCKEACSFKEWTLTTSLAQWPSEVSE  480
gamma-ENaC_Mesoplodon_bidens                QGANYCNYQQHPNWMYCYYELHKDFVRGELGCQSVCKEACSFKEWTLTTSLAQWPSEVSE  480
gamma-ENaC_Ziphius_cavirostris              QGANYCNYQQHPNWMYCYYELHEDFVRGELGCQSVCKEACSFKEWTLTTSLAQWPSEVSE  480
gamma-ENaC_Platanista_gangetica             RGANYCNYQQHPNWMYCYYELHQAFVREELGCQSLCKEACSFKEWTLTTSLAQWPSEVSE  480
gamma-ENaC_Platanista_minor                 RGANYCNYQQHPNWMYCYYELHQAFVREELGCQSLCKEACSFKEWTLTTSLAQWPSEVSE  480
gamma-ENaC_Kogia_breviceps                  QGVNYCNYQQHPNWMYCYYELHQDFVREELGCQSLCKEACSFKEWTLTTSLAQWPSEVSE  479
gamma-ENaC_Physeter_catodon                 QGVNYCNYQQHPNWMYCYYELHQDFVREELGCQSMCKEACSFKEWTLTTSLAQWPSEVSE  480
gamma-ENaC_Balaenoptera_musculus            RGANYCNYQQHPNWMYCYYELHQDFVREELGCQALCKEACSFKEWTLTTSLAQWPSEVSE  480
gamma-ENaC_Eubalaena_japonica               RGANYCNYQQHPNWMYCYYELHQDFVREELGCQVLCKEACSFKEWTLTTSLAQWPSEVSE  480
gamma-ENaC_Hippopotamus_amphibius           QGANYCNYQQHPNWMYCYYQLHQAFVREELGCQSVCKEACSFKEWTLTTSLAQWPSEVSE  477
gamma-ENaC_Tragulus_javanicus               RGADYCNYQQHPNWMYCYYQLHQAFVREELGCQSVCKEACSFKEWTLTTSLAQWPSEVSE  480
gamma-ENaC_Tragulus_kanchil                 RGADYCNYQQHPNWMYCYYQLHQAFVREELGCQSVCKEACSFKEWTLTTSLAQWPSEVSE  480
gamma-ENaC_Antilocapra_americana            QGADYCNYQQHPNWMYCYYQLHQAFVREELGCQSVCKEACSFKEWTLTTSLAQWPSEVSE  480
gamma-ENaC_Giraffa_camelopardalis           QGANYCNYQQHPNWMYCYYQLHQAFVREELGCQSVCKEACSFKEWTLTTSLAQWPSEVSE  480
gamma-ENaC_Giraffa_tippelskirchi            QGANYCNYQQHPNWMYCYYQLHQAFVREELGCQSVCKEACSFKEWTLTTSLAQWPSEVSE  480
gamma-ENaC_Capreolus_pygargus               QGADYCNYQQHPNWMYCYYKLHQAFVREELGCQSVCKEACSFKEWTLTTSLAQWPSEVSE  480
gamma-ENaC_Cervus_elaphus                   QGADYCNYQQHPNWMYCYYQLHQAFVREELGCQSVCKEACSFKEWTLTTSLAQWPSEVSE  480
gamma-ENaC_Moschus_moschiferus              QGADYCNYQQHPNWMYCYYQLHQAFVREELGCQSVCKEACSFKEWTLTTSLAQWPSEVSE  480
gamma-ENaC_Moschus_berezovskii              QGADYCNYQQHPNWMYCYYQLHQAFVREELGCQSVCKEACSFKEWTLTTSLAQWPSEVSE  480
gamma-ENaC_Bos_grunniens                    RGADYCNYQQHPNWMYCYYQLHQAFVREELGCQSVCKEACSFKEWTLTTSLAQWPSEVSE  480
gamma-ENaC_Bos_taurus                       RGADYCNYQQHPNWMYCYYQLHQAFVREELGCQSVCKEACSFKEWTLTTSLAQWPSEVSE  480
gamma-ENaC_Bubalus_bubalis                  RGADYCNYQQHPNWMYCYYQLHQAFVREELGCQSVCKEACSFKEWTLTTSLAQWPSEVSE  480
gamma-ENaC_Nanger_granti                    QGADYCNYQQHPNWMYCYYQLHQAFVREELGCQSVCKEACSFKEWTLTTSLAQWPSEVSE  480
gamma-ENaC_Kobus_leche                      QGADYCNYQQHPNWMYCYYQLHQAFVREELGCQSVCKEACSFKEWTLTTSLAQWPSEVSE  480
gamma-ENaC_Capra_hircus                     QGADYCNYQQHPNWMYCYYQLHQAFVREELGCQSVCKEACSFKEWTLTTSLAQWPSEVSE  480
gamma-ENaC_Ovis_aries                       QGANYCNYQQHPNWMYCYYQLHQAFVREELGCQSVCKEACSFKEWTLTTSLAQWPSEVSE  480
gamma-ENaC_Ovis_canadensis                  QGANYCNYQQHPNWMYCYYQLHQAFVREELGCQSVCKEACSFKEWTLTTSLAQWPSEVSE  480
gamma-ENaC_Oreamnos_americanus              QGADYCNYQQHPNWMYCYYQLHQAFVREELGCQSVCKEACSFKEWTLTTSLAQWPSEVSE  480
gamma-ENaC_Hippotragus_niger                QGADYCNYQQHPNWMYCYYQLHQAFVREELGCQSVCKEACSFKEWTLTTSLAQWPSEVSE  480
gamma-ENaC_Damaliscus_lunatus               QGADYCNYQQHPNWMYCYYQLHQAFVREELGCQSVCKEACSFKEWTLTTSLAQWPSEVSE  480
gamma-ENaC_Sus_scrofa                       PPANYCNYQQHPNWMYCYYQLSQAFVREELGCQSVCKEACSFKEWTLTTSLAQWPSEVSE  480
gamma-ENaC_Vicugna_pacos                    PKVNYCNYQQHPNWMYCFYQLHQAFVREELGCQSVCREACSFKEWSLTTSLAQWPSEVSE  480
gamma-ENaC_Equus_callabus                   PAANYCNYQQHPNWMYCYYQLHQAFVREELGCQSVCREACSFKEWTLTTSLAQWPSEVSE  480
                                              ..****:********:*.* . **  ***** :*:********:********** ***

                                                                                     TM2
gamma-ENaC_Homo_sapiens                     KWLLPVLTWDQGRQVNKKLNKTDLAKLLIFYKDLNQRSIMESPANSIEMLLSNFGGQLGL  537
gamma-ENaC_Globicephala_melas               KWLLSVLTWDQG-QIKKKLNKTDLAKLLIFYKDLNQRSIVESPANSIEMLLSNIGGQLGL  539
gamma-ENaC_Lagenorhynchus_obliquidens       KWLLSVLTWDQG-QIKKKLNKTDLAKLLIFYKDLNQRSIVESPANSIEMLLSNIGGQLGL  539
gamma-ENaC_Tursiops_truncatus               KWLLSVLTWDQG-QIKKKLNKTDLAKLLIFYKDLNQRSIVESPANSIEMLLSNIGGQLGL  539
gamma-ENaC_Orcinus_orca                     KWLLSVLTWDQG-QIKKKLNKTDLAKLLIFYKDLNQRSIVESPANSIEMLLSNIGGQLGL  539
gamma-ENaC_Phocoena_sinus                   KWLLSVLTWDQG-QIKKKLNKTDLAKLLIFYKDLNQRSIVESPANSIEMLLSNIGGQLGL  539
gamma-ENaC_Neophocaena_asiaeorientalis      KWLLSVLTWDQG-QIKKKLNKTDLAKLLIFYKDLNQRSIVESPANSIEMLLSNIGGQLGL  539
gamma-ENaC_Monodon_monocerons               KWLLSVLTWDQG-QIKKKLNKTDLAKLLIFYKDLNQRSIVESPANSIEMLLSNIGGQLGL  539
gamma-ENaC_Delphinapterus_leucas            KWLLSVLTWDQG-QIKKKLNKTDLAKLLIFYKDLNQRSIVESPANSIEMLLSNIGGQLGL  539
gamma-ENaC_Pontoporia_blainvillei           KWLLSVLTWDQG-QIKKKLNKTDLAKLLIFYKDLNQRSIVESPANSIEILLSNIGGQLGL  539
gamma-ENaC_Inia_geoffrensis                 KWLLSVLTWDQG-QIKRKLNKTDLAKLLIFYKDLNQRSIVESPANSIEMLLSNIGGQLGL  539
gamma-ENaC_Lipotes_vexillifer               KWLLSVLTWDQG-QIKKKLNKTDLAKLLIFYKDLSHRSIVESPANSIEMLLSNIGGQLGL  539
gamma-ENaC_Hyperoodon_ampullatus            KWLLSVLTWDQG-QIKKKLNKTDLAKLLIFYKDLNQRSIVESPANSIEMLLSNIGGQLGL  539
gamma-ENaC_Mesoplodon_bidens                KWLLSVLTWDQG-QIKKKLNKTDLAKLLIFYKDLNQRSIVESPANSIEMLLSNIGGQLGL  539
gamma-ENaC_Ziphius_cavirostris              KWLLSVLTWDQG-QIKKKLNKTDLAKLLIFYKDLNQRSIVESPANSIEMLLSNIGGQLGL  539
gamma-ENaC_Platanista_gangetica             KWLLSLLTWDQG-QIKKKLNKTDLAKLLIFYKDLNQRSIVESPANSIEMLLSNIGGQLGL  539
gamma-ENaC_Platanista_minor                 KWLLSLLTWDQG-QIKKKLNKTDLAKLLIFYKDLNQRSIVESPANSIEMLLSNIGGQLGL  539
gamma-ENaC_Kogia_breviceps                  KWLLSVLTWDQG-KIKKKLNKTDLAKLLIFYKDLNQRSIVESPANSIEMLLSNIGGQLGL  538
gamma-ENaC_Physeter_catodon                 KWLLSVLTWDQG-QIKKKLNKTDLAKLLIFYKDLNQRSIVESPANSIEMLLSNIGGQLGL  539
gamma-ENaC_Balaenoptera_musculus            KWLLSVLTWDQG-QIKKKLNKTDLAKLLIFYKDLNQRSIVESPANSIEMLLSNIGGQLGL  539
gamma-ENaC_Eubalaena_japonica               KWLLSVLTWDQG-QIKKKLNKTDLAKLLIFYKDLNQRSIVESPANSIEMLLSNIGGQLGL  539
gamma-ENaC_Hippopotamus_amphibius           KWLLSVLTWDQRQLINRKLNKTDLAKLLIYYKDLNQRSIMESPANSIEMLLSNIGGQLGL  537
gamma-ENaC_Tragulus_javanicus               KWLLSLLTWDQSQQIKKKLNKTDLAQLLIFYKDLNQRSIMENPANSIEQLLSNIGGQLGL  540
gamma-ENaC_Tragulus_kanchil                 KWLLSLLTWDQSQQIKKKLNKTDLAQLLIFYKDLNQRSIMENPANSIEQLLSNIGGQLGL  540
gamma-ENaC_Antilocapra_americana            KWLLSVLTWDQSQQIKKKLNKTDLAKLLIFYKDLNQRSIMENPANSIEQLLSNIGGQLGL  540
gamma-ENaC_Giraffa_camelopardalis           KWLLSVLTWDQSQQIKKKLNKTDLAKLLIFYKDLNQRSIMENPANSIEQLLSNIGGQLGL  540
gamma-ENaC_Giraffa_tippelskirchi            KWLLSVLTWDQSQQIKKKLNKTDLAKLLIFYKDLNQRSIMENPANSIEQLLSNIGGQLGL  540
```

```
gamma-ENaC_Capreolus_pygargus               KWLLSVLTWDQSQQIKKKLNKTDLAKLLIFYKDLNQRSIMENPANSIEQLLSNIGGQLGL  540
gamma-ENaC_Cervus_elaphus                   KWLLSVLTWDQSQQIKKKLNKTDLAKLLIFYKDLNQRSIVENPANSIEQLLSNIGGQLGL  540
gamma-ENaC_Moschus_moschiferus              KWLLSVLTWDQSQQIKKKLNKTDLAKLLIFYKDLNQRSIMENPANSIEQLLSNIGGQLGL  540
gamma-ENaC_Moschus_berezovskii              KWLLSVLTWDQSQQIKKKLNKTDLAKLLIFYKDLNQRSIMENPANSIEQLLSNIGGQLGL  540
gamma-ENaC_Bos_grunniens                    KWLLSILTWDQSQQIKKKLNKTDLAKLLIFYKDLNQRSIMENPANSIEQLLSNIGGQLGL  540
gamma-ENaC_Bos_taurus                       KWLLSILTWDQSQQIKKKLNKTDLAKLLIFYKDLNQRSIMENPANSIEQLLSNIGGQLGL  540
gamma-ENaC_Bubalus_bubalis                  KWLLSILTWDQSQQIKKKLNKTDLAKLLIFYKDLNQRSIMENPANSIEQLLSNIGGQLGL  540
gamma-ENaC_Nanger_granti                    KWLLSVLTWDQSQQIKKKLNKTDLAKLLIFYKDLNQRSIMENPANSIEQLLSNIGGQLGL  540
gamma-ENaC_Kobus_leche                      KWLLSVLTWDQSQQIKKKLNKTDLAKLLIFYKDLNQRSIMENPANSIEQLLSNIGGQLGL  540
gamma-ENaC_Capra_hircus                     KWLLSVLTWDQSQQIKKKLNKTDLAKLLIFYKDLNQRSIMENPANSIEQLLSNIGGQLGL  540
gamma-ENaC_Ovis_aries                       KWLLSVLTWDQSQQIKKKLNKTDLAKLLIFYKDLNQRSIMENPANSIEQLLSNIGGQLGL  540
gamma-ENaC_Ovis_canadensis                  KWLLSVLTWDQSQQIKKKLNKTDLAKLLIFYKDLNQRSIMENPANSIEQLLSNIGGQLGL  540
gamma-ENaC_Oreamnos_americanus              KWLLSVLTWDQSQQIKKKLNKTDLAKLLIFYKDLNQRSIMENPANSIEQLLSNIGGQLGL  540
gamma-ENaC_Hippotragus_niger                KWLLSVLTWDQSQQIKKKLNKTDLAKLLIFYKDLNQRSIMENPANSIEQLLSNIGGQLGL  540
gamma-ENaC_Damaliscus_lunatus               KWLLSVLTWDQSQQIKKKLNKTDLAKLLIFYKDLNQRSIMENPANSIEQLLSNIGGQLGL  540
gamma-ENaC_Sus_scrofa                       KWLLSVLTWDQGQQIKKKLNKTDLAKLLIFYKDLNQRSIMESPANSIEMLLSNIGGQLGL  540
gamma-ENaC_Vicugna_pacos                    KWLLSVLTWDQSQQIKKKLNKTDLAKLLIFYKDLNQRSIMESPANSIEMLLSNIGGQLGL  540
gamma-ENaC_Equus_callabus                   KWLLSVLTWDQGLQIKKKLNKTDLAKLLIFYKDLNQRSIMENPANSIEMLLSNFGGQLGL  540
                                            **** :*****   :::********:***:****.:***:*.****** ****:******

gamma-ENaC_Homo_sapiens                     WMSCSVVCVIEIIEVFFIDFFSIIARRQWQKAKEWWAWKQAPPCPEAP-RSPQGQDNPAL  596
gamma-ENaC_Globicephala_melas               WMSCSIVCIIEIIEVFFIDSLSIVARHQWHKAKEWWARRQASPCPEAPP-SPQGQDNLGL  598
gamma-ENaC_Lagenorhynchus_obliquidens       WMSCSIVCIIEIIEVFFIDSLSIVARHQWHKAKEWWARRQASPCPEAPP-SPQGQDNLGL  598
gamma-ENaC_Tursiops_truncatus               WMSCSIVCIIEIIEVFFIDSLSIVARHQWHKAKQWWARRQASPCPEAPP-SPQGQDNLGL  598
gamma-ENaC_Orcinus_orca                     WMSCSIVCIIEIIEVFFIDSLSIVARHQWHKAKEWWARRQASPCPEAPP-SPQGQDNLGL  598
gamma-ENaC_Phocoena_sinus                   WMSCSIVCIIEIIEVFFIDSLSIVARHQWHKAKEWWARRQASPCPEAPP-SPQGQDNLGL  598
gamma-ENaC_Neophocaena_asiaeorientalis      WMSCSIVCIIEIIEVFFIDSLSIVARHQWHKAKEWWARRQASPCPEAPP-SPQGQDNLGL  598
gamma-ENaC_Monodon_monocerons               WMSCSIVCIIEIIEVFFIDSLSIVARHQWHKAKEWWARRQASPCPEAPP-SPQGQDNLGL  598
gamma-ENaC_Delphinapterus_leucas            WMSCSIVCIIEIIEVFFIDSLSIVARHQWHKAKEWWARRQASPCPEAPP-SPQGQDNLGL  598
gamma-ENaC_Pontoporia_blainvillei           WMSCSIVCIIEIIEVFFIDSLSIVARHQWHKAKEWWARRQASPCPEAPH-SPQGQDNLGL  598
gamma-ENaC_Inia_geoffrensis                 WMSCSIVCIIEIIEVFFIDSLSIVARHQWHKAKEWWARRQATPCPEAPP-SPQGQDNLGL  598
gamma-ENaC_Lipotes_vexillifer               WMSCSIICIIEIIEVFFIDSLSIVARHQWHKAKEWWARRQASPCPEAPP-SPQGQDNLGL  598
gamma-ENaC_Hyperoodon_ampullatus            WLSCSIVCVIEIIEVFFIDSLSIVARHQWHKAKEWWARRQASPRPEAPP-SPQGQDNLGL  598
gamma-ENaC_Mesoplodon_bidens                WLSCSIVCVIEIIEVFFIDSLSIVARHQWHKAKEWWARRQASPHPEAPP-SPQGQDNLGL  598
gamma-ENaC_Ziphius_cavirostris              WLSCSIVCVIEIIEVFFIDSLSIVARHQWHKAKEWWARRQASPRPEAPP-SPQGQDNLGL  598
gamma-ENaC_Platanista_gangetica             WMSCSIVCVIEIIEVFFIDSLSIVARHQWHKAKEWWARRQASPCPEAPP-SPQGQDNLGL  598
gamma-ENaC_Platanista_minor                 WMSCSIVCVIEIIEVFFIDSLSIVARHQWHKAKEWWARRQASPCPEAPP-SPQGQDNLGL  598
gamma-ENaC_Kogia_breviceps                  WMSCSIVCIIEIIEVFFIDSLSIVARHQWHKAKEWWARRQATTCPEAPP-SPQGRDNLGL  597
gamma-ENaC_Physeter_catodon                 WMSCSIVCVIEIIEVFFIDSLSIVARHQWHKAKEWWARRQASTCPEAPP-SPQGQDNLGL  598
gamma-ENaC_Balaenoptera_musculus            WMSCSVVCVIEIVEVFLVDSLSIVARHQWRKAKEWWARRQAPPWPEAPP-SPQGRDNLGL  598
gamma-ENaC_Eubalaena_japonica               WMSCSVVCVIEIVEVFLVDSLSIVARHQWRKAKEWWARRQAPPWPEATH-SPQGRDNLGL  598
gamma-ENaC_Hippopotamus_amphibius           WMSCSVVCVIEIIEVFFIDSLSIIARHQWHKAKGWWARRQAPPCPEAP-PSPQGQDNPGL  596
gamma-ENaC_Tragulus_javanicus               WMSCSVVCVIEIIEVFFIDSLSIIARHQWHKARDWWARRRAPPCPEAP-RSPQGQDNPGL  599
gamma-ENaC_Tragulus_kanchil                 WMSCSVVCVIEIIEVFFIDSLSIIARHQWHKARDWWARRRAPPCPEAP-RSPQGQDNPGL  599
gamma-ENaC_Antilocapra_americana            WMSCSVVCVIEIIEVFFIDSLSIIARHQWHKAKGWWARRQAPTCPEAP-RRPQGQDNAGL  599
gamma-ENaC_Giraffa_camelopardalis           WMSCSVVCVIEIIEVFFIDSLSIIARHQWHKAKGWWARRRAPTCPEAP-RSPQGQDNPGL  599
gamma-ENaC_Giraffa_tippelskirchi            WMSCSVVCVIEIIEVFFIDSLSIIARHQWHKAKGWWARRRAPTCPEAP-RSPQGQDNPGL  599
gamma-ENaC_Capreolus_pygargus               WMSCSVVCVIEIIEVFFIDSLSIIARQQWHKAKGWWARRRAPACPEAP-RTRQGRVNPGL  599
gamma-ENaC_Cervus_elaphus                   WMSCSIVCVIEIIEVFFIDSLSIIARQQWHKAKGWWARRRAPACPEAP-RTRQGRVNPGL  599
gamma-ENaC_Moschus_moschiferus              WMSCSVVCVIEIIEVFFIDSLSIIARHQWHKAKGWWAQRRAPACPEAP-HTPQGRDNPGL  599
gamma-ENaC_Moschus_berezovskii              WMSCSVVCVIEIIEVFFIDSLSIIARHQWHKAKGWWAQRRAPACPEAP-HTPQGRDNPGL  599
gamma-ENaC_Bos_grunniens                    WMSCSVVCVIEIIEVFFIDSLSIIARHQWHKAKGWWARRRAPACPEAP-RAPQGRDNPSL  599
gamma-ENaC_Bos_taurus                       WMSCSVVCVIEIIEVFFIDSLSIIARHQWHKAKGWWARRRAPACPEAP-RAPQGRDNPSL  599
gamma-ENaC_Bubalus_bubalis                  WMSCSVVCIIEIIEVFFIDSLSIIARHQWHKAKRWWARRRAPACPEAP-RAPQGRDNPSL  599
gamma-ENaC_Nanger_granti                    WMSCSVVCVIEIIEVFFIDSLSIIARHQWHKAKGWWARRRAPTCPEAP-RIPQGRDNPGL  599
gamma-ENaC_Kobus_leche                      WMSCSVVCVIEIIEVFFIDSLSIIARHQWHKAKGWWARRRAPACPEVP-RIPQGRDNPGL  599
gamma-ENaC_Capra_hircus                     WMSCSVVCVIEIIEVFFIDSLSIIARHQWHKAKGWWARRRAPACPEAP-HIPQGRDNPGL  599
gamma-ENaC_Ovis_aries                       WMSCSVVCVIEIIEVFFIDSLSIIARHQWHKAKGWWARRRAPACPEAP-HIPQGRDNPGL  599
gamma-ENaC_Ovis_canadensis                  WMSCSVVCVIEIIEVFFIDSLSIIARHQWHKAKGWWARRRAPACPEAP-HIPQGRDNPGL  599
gamma-ENaC_Oreamnos_americanus              WMSCSIVCVIEIIEVFFIDSLSIIARHQWHKAKGWWARRRAPACPEAP-HIPHGRDNPGL  599
gamma-ENaC_Hippotragus_niger                WMSCSVVCVIEIIEVFFIDSLSIIARHQWHKAKGWWARRRAPTCHEAP-HIPQGRDNPGL  599
gamma-ENaC_Damaliscus_lunatus               WMSCSVVCVIEIIEVFFIDSLSIIARHQWHKAKGWWARRRAPTCCEAP-RIPQGRDNPGL  599
gamma-ENaC_Sus_scrofa                       WMSCSVVCVIEIIEVFFIDSLSIIARHQWHKAKGWWARRQAPPCPEAP-PSPQGRDNPAL  599
gamma-ENaC_Vicugna_pacos                    WMSCSVVCVIEIIEVFFIDSLSIIARHQWHKAKEWWARRQAPACPEAPPRSPQGQVNPVL  600
gamma-ENaC_Equus_callabus                   WMSCSIVCIIEIVEVFFIDSLSIIARQQWQKAKQWWAQRQAPPCPEAPP-SPQGQDNPVL  599
                                            *:***::*:***:***:* :**:**:**: *** ::*    *.      :*: *  *

gamma-ENaC_Homo_sapiens                     DIDDDLPTFNSALHLPPALGTQVPGTPPPKYNTLRLERAFSNQLTDTQMLDEL---     649
gamma-ENaC_Globicephala_melas               NIDDDLPTFTSALCLPPAPAAQVPGTPPPRYNTLRLERAFSRQLTGTEVSAES---     651
gamma-ENaC_Lagenorhynchus_obliquidens       NIDDDLPTFTSALCLPPAPAAQVPGTPPPRYNTLRLERAFSRQLTGTEVSAES---     651
gamma-ENaC_Tursiops_truncatus               NIDDDLPTFTSALCLPPAPAAQVPGTPPPRYNTLRLERAFSRQLTGTEVSAES---     651
gamma-ENaC_Orcinus_orca                     HIDDDLPTFTSALCLPPAPAAQVPGTPPPRYNTLRLERAFSRQLTGTEVSAES---     651
gamma-ENaC_Phocoena_sinus                   NIDDDLPTFTSALCLPPAPAAQVPGTPPPRYNTLSLERAFSRQLTGTEVPAES---     651
gamma-ENaC_Neophocaena_asiaeorientalis      NIDDDLPTFTSALCLPPAPAAQVPGTPPPRYNTLSLERAFSRQLTGTEVPAES---     651
gamma-ENaC_Monodon_monocerons               NIDDDLPTFTSALCLPPAPAAQVPGTPPPRYNTLRLERTFSRQLTGTEVPAES---     651
gamma-ENaC_Delphinapterus_leucas            NMDDDLPTFTSALCLPPAPAAQVPGTPPPRYNTLRLERTFSRQLTGTEVPAES---     651
gamma-ENaC_Pontoporia_blainvillei           NIDDDLPTFTSALGLPPAPAAQVPGTPPPRYNTLRLERAFSRQLTDTEVPAES---     651
gamma-ENaC_Inia_geoffrensis                 NIDDDLPTFTSALCLPPAPAAQVPGTPPPRYNTLRLERAFSRQLTDTEVPAES---     651
gamma-ENaC_Lipotes_vexillifer               NIDDDLPTFTSALCLPPAPAAQVPGTPPPRYNTLRLERAFSRQLTGTEVPAES---     651
gamma-ENaC_Hyperoodon_ampullatus            HIDDDLPTFTSALCLPPAPAAQVPGTPPPRYNTLRLERAFSRQLTGTEVPAES---     651
gamma-ENaC_Mesoplodon_bidens                HIDDDLPTFTSALCLPPAPAAQVPGTPPPRYNTLRLERAFSRQLTGTEVPAES---     651
gamma-ENaC_Ziphius_cavirostris              HIDDDLPTFTSALCLPPAPAAQVPGTPPPRYNTLSLERAFSRQLTGTEVPAES---     651
gamma-ENaC_Platanista_gangetica             NIDDDLPTFTSALCLPPAPAAQVPGTPPPRYNTLRLERAFSRQLTDTEVPAES---     651
gamma-ENaC_Platanista_minor                 NIDDDLPTFTSALCLPPAPAAQVPGTPPPRYNTLRLERAFSRQLTDTEVPAES---     651
```

```
gamma-ENaC_Kogia_breviceps                  DVDDDLPTFTSALCLPPAPAAQVPGTPPPRYNTLRLDRAFSRQLTDTEVPAES---     650
gamma-ENaC_Physeter_catodon                 DVDDDLPTFTSALCLPPAPGGQVPGTPPPRYNTLRLDRTFSRQLTDTEVPAES---     651
gamma-ENaC_Balaenoptera_musculus            DIDDDLPTFTSALCLPPAPAAQVPGTPPPRYNTLRLERDFPSQLADTEVPAES---     651
gamma-ENaC_Eubalaena_japonica               DIDDDLPTFTSALCLPPAPAAQVPGTPPPRYNTLRLERDFPSQLADTEVPAES---     651
gamma-ENaC_Hippopotamus_amphibius           VIDDDLPTFTSALRLPPAPAAQVPGTPPPRYNTLRLERAFASQLTDTQVPAES---     649
gamma-ENaC_Tragulus_javanicus               EIDNDLPTFTSALSLPPAPGAQVPGTPPPRYNTLRLERAFSSQLTDTQAPAES---     652
gamma-ENaC_Tragulus_kanchil                 EIDNDLPTFTSALSLPPAPGAQVPGTPPPRYNTLRLERAFSSQLTDTQAPAES---     652
gamma-ENaC_Antilocapra_americana            DIDDDLPTFTSALSLPPAPGAQVPGTPPPRYNTLRLERAFSSQLTDTQVPAES---     652
gamma-ENaC_Giraffa_camelopardalis           YIDDDLPTFTSALSLPPAPGAQVPGTPPPRYNTLRLERAFSSQLTDTQVPTES---     652
gamma-ENaC_Giraffa_tippelskirchi            YIDDDLPTFTSALSLPPAPGAQVPGTPPPRYNTLRLERAFSSQLTDTQVPTES---     652
gamma-ENaC_Capreolus_pygargus               NIDDDLPTFTSALSLPPAPGAQVPGTPPPRYNTLRLERAFSSQLTDTQEAAES---     652
gamma-ENaC_Cervus_elaphus                   DIDDDLPTFTSALSLPPAPGAQVPGTPPPRYNTLRLERAFSSQLTDTQEAAES---     652
gamma-ENaC_Moschus_moschiferus              NIDDDLPTFTSALSLPPAPGAQVPGTPPPRYNTLRLERAFSSQLTDTQVPAES---     652
gamma-ENaC_Moschus_berezovskii              NIDDDLPTFTSALSLPPAPGAQVPGTPPPRYNTLRLERAFSSQLTDTQVPAES---     652
gamma-ENaC_Bos_grunniens                    DIDDDLPTFTSALSLPPAPGSQVPGTPPPRYNTLRLERAFSSQLTDTQVPAES---     652
gamma-ENaC_Bos_taurus                       DIDDDLPTFTSALSLPPAPGSQVPGTPPPRYNTLRLERAFSSQLTDTQVPAES---     652
gamma-ENaC_Bubalus_bubalis                  DIDDDLPTFTSALSLPPAPGSQVPGTPPPRYNTLRLERAFSSQLSDTQVPAES---     652
gamma-ENaC_Nanger_granti                    NIDDDLPTFTSALSLPPAPGAQVPGTPPPRYNTLRLERAFSSQLTDTQVPAES---     652
gamma-ENaC_Kobus_leche                      NIDDDLPTFTSALSLPPAPGAQVPGTPPPRYNTLRLERAFSSQLTDTQVPAES---     652
gamma-ENaC_Capra_hircus                     NIDDDLPTFTSALSLPPAPGPQVPGTPPPRYNTLRLERAFSSQLTDTQVPAES---     652
gamma-ENaC_Ovis_aries                       NIDDDLPTFTSALSLPPAPGPQVPGTPPPRYNTLRLERAFSSQLTDTQVPAES---     652
gamma-ENaC_Ovis_canadensis                  NIDDDLPTFTSALSLPPAPGPQVPGTPPPRYNTLRLERAFSSQLTDTQVPAES---     652
gamma-ENaC_Oreamnos_americanus              NIDDDLPTFTSALSLPPAPGPQVPGTPPPRYNTLRLERAFSSQLTDTQVPAES---     652
gamma-ENaC_Hippotragus_niger                NIDDDLPTFTSALSLPPAPGPQVPGTPPPRYNTLRLERAFSSQLTDTQVPAES---     652
gamma-ENaC_Damaliscus_lunatus               NIDDDLPTFTSALSLPPAPGPQVPGTPPPRYNTLRLERAFSSQLTDTQVPAES---     652
gamma-ENaC_Sus_scrofa                       NIDDDLPTFTSALRLPPAPAAQVPGTPPPRYNTLRLERAFSSQLSDTQVPAES---     652
gamma-ENaC_Vicugna_pacos                    ELDDDLPTFTSALCLPPAPGAQVPGTPPPKYNTLRLERTFSSQLTDTQVPPESPES     656
gamma-ENaC_Equus_callabus                   DIDDDLPTFTSALHLPPAPGTQVPGTPPPKYNTLRLERAFSNQLTDTQVPAEF---     652
                                             :*:*****.*** **** . ********:**** *:* *  **:.*:   *
```

**δ-ENaC (*SCNN1D*) amino acid sequence alignment** created with Clustal Omega (v.1.2.4, accessed 24.03.2024). Putative structural motifs are highlighted based on the Alphafold2 prediction of the human ENaC subunit (https://alphafold.ebi.ac.uk/entry/A0A024R0D5; accessed 24.03.2024). Transmembrane domains (TM1/TM2) are highlighted in yellow. An N-Terminal HG-motif affecting ENaC open probability is marked in magenta. Residues putatively forming the selectivity filter within TM2 are shown in red font. The incoporated amino acids due to exon 11/12 fusion in the Antilopinae is highlighted in blue.

```
delta-ENaC_Homo_sapiens                MAEHRSMDGRMEAATRGGSHLQAA----------------AQTPPRP---------GPP      34
delta-ENaC_Hippopotamus_amphibius      ----------MEMPTQGGSSLE--AGALAQAAGRGPGTCTCLQLPPPPPP----------      38
delta-ENaC_Tragulus_javanicus          ----------MEAPARGGSSLK--AEGAAQAAGGGSGTCMCPQPPPRPPR----------      38
delta-ENaC_Tragulus_kanchil            ----------MEAPAR-----------------GGSSLKMCPQPPPRPPR----------      23
delta-ENaC_Antilocapra_americana       ----------MEAPAQGGCSLE--AEGTGQTVGGGPGTWTCPQAP---PL----------      35
delta-ENaC_Giraffa_camelopardalis      ----------MEAHAQGGCSFEMQAEGTGQTVGGGPGTWTCPQAP---PP----------      37
delta-ENaC_Giraffa_tippelskirchi       ----------MEAHAQGGCSFE--AEGTGQTVGGGPGTWTCPQAP---PP----------      35
delta-ENaC_Capreolus_pygargus          ----------MDARAQGGCSLEMQTEGTGQTVGGGPGTWTCSQAP---PP----------      37
delta-ENaC_Cervus_elaphus              ----------MDAHAQGGCSLEMQAEGTGQTVGGGPGTWMCSQAP---PP----------      37
delta-ENaC_Moschus_moschiferus         ----------MDAHAQGGCSLEM-----GQTVGGGPGTWTCPQAP---PP----------      32
delta-ENaC_Moschus_berezovskii         ----------MDAHAQGGCSLEM-----GQTVGGGPGTWTCPQAP---PP----------      32
delta-ENaC_Bos_grunniens               ----------MDAHAQGGCSLEMQAEGTGQTVGGGPGTWTCPQAS---PP----------      37
delta-ENaC_Bos_taurus                  ----------MDAHAQGGCSLEMQAEGTGQTVGGGPGTWTCPQAS---PP----------      37
delta-ENaC_Bubalus_bubalis             ----------MDAHAQGGCSLKMQAEGTGQTVGGGPGTWTCPQAS---PP----------      37
delta-ENaC_Nanger_dama                 ----------MDAHPQGGCSLEMQAEGTGQTVGGGPGTWTCPQAS---PP----------      37
delta-ENaC_Kobus_leche                 ----------MDAHAQGGCSLEMQAEGTGQTVGGGPGTWTCPQAS---PP----------      37
delta-ENaC_Capra_hircus                ----------MDADAQGGCSLEMQAEGTGQTVGEGPGTWMCPQAS---PP----------      37
delta-ENaC_Ovis_aries                  ----------MDAHAQGGCSLEMQAEGTGQTVGDGPGTWTCPQAS---PP----------      37
delta-ENaC_Ovis_canadensis             ----------MDAHAQGGCSLEMQAEGTGQTVGDGPGTWTCPQAS---PP----------      37
delta-ENaC_Oreamnos_americanus         ----------MDAHVQGGCSLEMQAEGTGQ-VGEGPGTWMCPQAS---PP----------      36
delta-ENaC_Hippotragus_niger           ----------MDAHAQGGCSLEMQAEGTGQTVGGGAGTWTCPQAS---PP----------      37
delta-ENaC_Damaliscus_lunatus          ----------MDAHAQGGCSLEMQAEGTGQTVGGGPGTWMCPQAS---PP----------      37
delta-ENaC_Sus_scrofa                  ----------MEEPAQGGSSLK--AGASAQAAGGGPGAGRCPQPL---------------      33
delta-ENaC_Vicugna_pacos               ----------MEAPAQGGCRLE--AGARAPEAGGGPGTRKL-------------------      29
delta-ENaC_Equus_callabus              ----------MEASAHRGSCFK--AGTVVQTTSEGLGIWKYPQGQPPLPPLPEDTPPRPE      48
                                                 *:   :

                                                                                         TM1?
delta-ENaC_Homo_sapiens                SAPPPPPKEGHQEGLVELPASFRELLTFFCTNATIHGAIRLVCSRGNRLKTTSWGLLSLG      94
delta-ENaC_Hippopotamus_amphibius      ----PAPEEERRERLVELHTSFREMITFFCTNATIHGTIRLVCSSQNRLKTASWGLLLVG      94
delta-ENaC_Tragulus_javanicus          ----PLPEEEPGERLVEPHASFRELVTFFCTNSTIHGTIRLVCSSQNRLKTASWGLLLAG      94
delta-ENaC_Tragulus_kanchil            ----PLPEEEPGERLVEPHASFRELVTFFCTNSTIHGTIRLVCSSQNRLKTASWGLLLAG      79
delta-ENaC_Antilocapra_americana       ----TLPEEEHGERLVELHASFRELITFFCTNSTIHGTIRLVCSSQNRLKTASWGLLLGG      91
delta-ENaC_Giraffa_camelopardalis      ----MLPKEERGERLVELHASYRELVTFFCTNSTIHGTIRLVCSSQNRLKTASWGLLLAG      93
delta-ENaC_Giraffa_tippelskirchi       ----TLPKEERGERLVELHASYRELVTFFCTNSTIHGTIRLVCSSQNRLKTASWGLLLAG      91
delta-ENaC_Capreolus_pygargus          ----TLPEEERGERLVELHTSFRELVTFFCTNSTIHGTIRLVCSSQNRLKTASWGLLLAG      93
delta-ENaC_Cervus_elaphus              ----TLPEEERGERLIELHASFRELVTFFCTNSTIHGTIRLVCSSQNRLKTASWGLLLAG      93
delta-ENaC_Moschus_moschiferus         ----TLPEEEHGERLVELHTSFRELVTFFCTNSTIHGTIRLVCSSQNHLKTASWGLLLAG      88
delta-ENaC_Moschus_berezovskii         ----TLPEEEHGERLVELHTSFRELVTFFCTNSTIHGTIRLVCSSQNHLKTASWGLLLAG      88
delta-ENaC_Bos_grunniens               ----TLPEEEHGERLVELHASFRELVTFFCTNSTIHGTIRLVCSSQNRLKTASWGLLLAG      93
delta-ENaC_Bos_taurus                  ----TLPEEEHGERLVELHASFRELVTFFCTNSTIHGTIRLVCSSQNRLKTASWGLLLAG      93
delta-ENaC_Bubalus_bubalis             ----TLPEEEHGERLVELHASFRELVTFFCTNSTIHGTIRLVCSSQNRLKTASWGLLLAG      93
delta-ENaC_Nanger_dama                 ----TLPEEEHGERLVELHASFRELVTFFCTNSTIHGTIRLVCSSQNRLKTASWGLLLAG      93
delta-ENaC_Kobus_leche                 ----TLSEEERGERLVELHSSFRELVTFFCTNSTIHGTIRLVCSSQNRLKTASWGLLLAG      93
delta-ENaC_Capra_hircus                ----TLPEEEHGERLVELHTSFRELVTFFCTNSTIHGTIRLVCSSQNRLKTASWGLLLAG      93
delta-ENaC_Ovis_aries                  ----TLPEEEHGERLVELHTSFRELVTFFCTNSTIHGTIRLVCSSQNRLKTASWGLLLAG      93
delta-ENaC_Ovis_canadensis             ----TLPEEEHGERLVELHTSFRELVTFFCTNSTIHGTIRLVCSSQNRLKTASWGLLLAG      93
delta-ENaC_Oreamnos_americanus         ----TLPEEEHGERLVELHTSFRELVTFFCTNSTIHGTIRLVCSSQNRLKTAFWGLLLAG      92
delta-ENaC_Hippotragus_niger           ----TLPEEEHGERLVELHASFRELVTFFCTNSTIHGTIRLVCSSQNRLKTASWGLLLAG      93
delta-ENaC_Damaliscus_lunatus          ----MLPEEEHRERLVELHASFRELVTFFCTNSTIHGTIRLVCSSQNRLKTASWGLLLAG      93
delta-ENaC_Sus_scrofa                  --PLPPSEEERREGLVELHTSFRELVTFFCTNATIHGTIRLVCSSQNRLKTATWALLLAG      91
delta-ENaC_Vicugna_pacos               -------EEECGERLVEFHASFRELVTFFCTNTTIHGSIRLICSSQNRLKTVSWGLLLVG      82
delta-ENaC_Equus_callabus              DRPL-PPEDRPGERLVELHTSFRELFTFFCTNATIHGTIRLVCSRQNRLKTASWGLLLVG      107
                                              ::   * *:*  :*:**:.******:****:***:**  *:***. *.**  *

delta-ENaC_Homo_sapiens                ALVALCWQLGLLFERHWHRPVLMAVSVHSERKLLPLVTLCDGNPRRPSPVLRHLELLDEF      154
delta-ENaC_Hippopotamus_amphibius      SLGMLYWQFGLLFEQYWRYPVSMTVSVRSERKLFPSVTLCDMNPHRPHAARHHLWALDDF      154
delta-ENaC_Tragulus_javanicus          ALGVLYWQFGLLFEQYWRYPVIMTVSVHSERKLLPSVTLCDMNPRWPRSAGRHLRALDDF      154
delta-ENaC_Tragulus_kanchil            ALGVLYWQFGLLFEQYWRYPVIMTVSVHSERKLLPSVTLCDMNPRWPRSAGRHLRALDDF      139
delta-ENaC_Antilocapra_americana       ALGVLYWQFALLLEQYWRYPVIMTVSVHSERKLFPSVTLCDMNPHRPHLARYHLRVLDDF      151
delta-ENaC_Giraffa_camelopardalis      ALGVLYWQFALLFEQYWRYPVIMTVSVHSERKLFPSVTLCDMNPHRPHLARHHLRVLDDF      153
delta-ENaC_Giraffa_tippelskirchi       ALGVLYWQFALLFEQYWRYPVIMTVSVHSERKLFPSVTLCDMNPHRPHLARHHLRVLDDF      151
delta-ENaC_Capreolus_pygargus          ALGVLYWQFALLFEQYWRYPVIMTVSVHSERKLFPSVTLCDMNPHRPHLARHHLRVLDDF      153
delta-ENaC_Cervus_elaphus              ALGVLYWQFALLFEQYWRYPVIMTVSVHSERKLFPSVTLCDMNPHRPHLARHHLRVLDDF      153
delta-ENaC_Moschus_moschiferus         ALGVLYWQFALLFEQYWRYPVIMTVSVHSERKVFPSVTLCDMNPHRPHLARHHLRALDDF      148
delta-ENaC_Moschus_berezovskii         ALGVLYWQFALLFEQYWRYPVIMTVSVHSERKVFPSVTLCDMNPHRPHLARHHLRALDDF      148
delta-ENaC_Bos_grunniens               ALGVLYWQFALLFEQYWRYPVIMTVSVHSERKLFPSVTLCDMNPHRPHLARHHLRVLDDF      153
delta-ENaC_Bos_taurus                  ALGVLYWQFALLFEQYWRYPVIMTVSVHSERKLFPSVTLCDMNPHRPHLARHHLRVLDDF      153
delta-ENaC_Bubalus_bubalis             ALGVLYWQFALLFEQYWRYPVIMTVSVHSERKLFPSVTLCDMNPHRPHLARHHLRVLDDF      153
delta-ENaC_Nanger_dama                 ALSVLYWQFALLFEQYWRYPVIMTVSVHSERKLFPSVTLCDMNPHRPHLARHHLRVLDDF      153
delta-ENaC_Kobus_leche                 ALGVLYWQFALLFEQYWRYPVIMTVSVHSERKLFPSVTLCDMNPHRPHLARHHLRVLDDF      153
delta-ENaC_Capra_hircus                ALGVLYWQFALLFEQYWHYPVIMTVSVHSERKLFPSVTLCDMNPHRPHLARHHLRILDDF      153
delta-ENaC_Ovis_aries                  ALGVLYWQFALLFEQYWRYPVIMTVSVHSERKLFPSVTLCDMNPHRPHLARHHLRILDDF      153
```

```
delta-ENaC_Ovis_canadensis            ALGVLYWQFALLFEQYWRYPVIMTVSVHSERKLFPSVTLCDMNPHRPHLARHHLRILDDF    153
delta-ENaC_Oreamnos_americanus        ALGVLYWQFALLFEQYWRYPVIMTVSVHSERKIFPSVTLCDMNPHRPHLARHHLRILDDF    152
delta-ENaC_Hippotragus_niger          ALGVLYWQFALLFEQYWRYPVIMTVSVHSERKLFPSVTLCDMNPHRPHLARHHLRVLDDF    153
delta-ENaC_Damaliscus_lunatus         ALGVLYWQFALLFEQYWRYPVIMTVSVHSERKLFPSVTLCDMNPHRPHLARHHLRVLDDF    153
delta-ENaC_Sus_scrofa                 ALSALYWQFGLLFQQYWRYPVVMAVSVHSERKLFPSVTLCDMNPHRPRPARRHLRALDDF    151
delta-ENaC_Vicugna_pacos              TLGMLYWQFGLLFEQYWRYPVIMTVSLHSERKLFPSVTLCDMNPHRPRPARRHLQALDDF    142
delta-ENaC_Equus_callabus             ALGALYWQFGLLFEQYWRYPVIMTVSVHSERKLFPSVTLCDMNPHRPSPARRHLQALDEF    167
                                      :*  * **:.**::::*: ** *:**::****::* ***** **: *  .  **  **:*

delta-ENaC_Homo_sapiens               ARENIDSLYNVNLSKGRAALSATVPRHEPPFHLDREIRLQRLSHSGSRVRVGFRLCNSTG    214
delta-ENaC_Hippopotamus_amphibius     ARENIYSLYRFNFSESRDAPGAEAPGPEPIFQLDRRVCLQRLSPPGGQNRVGFKLCNSAG    214
delta-ENaC_Tragulus_javanicus         AQENIYSLYGFNFSDSRDVPGAQAPGPEPTFQLDRGIRLQRLSPLDGQNRVGFRLCNSSG    214
delta-ENaC_Tragulus_kanchil           AQENIYSLYGFNFSDSRDVPGAQAPGPEPTFQLDRGIRLQRLSPLDGQNRVGFRLCNSSG    199
delta-ENaC_Antilocapra_americana      AQENIYSLYRFNFSDSRDTLGAEVPGPETAFQLDRRIRLQRLRPLNGQNRVGFKLCNSTG    211
delta-ENaC_Giraffa_camelopardalis     ARENIYSLYRFNFSDSRDALGAEVPGPEPAFQLDRRIRLQRLRPLDGQNRVGFKLCNSTS    213
delta-ENaC_Giraffa_tippelskirchi      ARENIYSLYRFNFSDSRDALGAEVPGPEPAFQLDRRIRLQRLRPLDGQNRVGFKLCNSTS    211
delta-ENaC_Capreolus_pygargus         ARENIYSLYQFNFSDSREALGAEVPGPEPAFQLDRRIRLQRLRPLEGQNRVGFKLCNSTG    213
delta-ENaC_Cervus_elaphus             ARENIYSLYQFNFSDSRDALGAEVPGPEPAFQLDRRIRLQRLRPLDGQNRVGFKLCNSTG    213
delta-ENaC_Moschus_moschiferus        ARESIYSLYRFNFTDSRDALGAEVPSPEPAFQLDRRIRLQRLRPLDGQNRVGFKLCNSTG    208
delta-ENaC_Moschus_berezovskii        ARESIYSLYRFNFSDSRNALGAEVPGPEPVFQLDRRIRLQRLRPLDGQNRVGFKLCNSTG    208
delta-ENaC_Bos_grunniens              ARESIYSLYRFNFSDSMDALGAEVPGPEPAFHLDRRIRLQRLRPLDGQNRVGFKLCNSTG    213
delta-ENaC_Bos_taurus                 ARESIYSLYRFNFSDSMDALGAEVPGPEPAFHLDRRIRLQRLRPLDGQNRVGFKLCNSTG    213
delta-ENaC_Bubalus_bubalis            ARESIYSLYRFNFSNSMDALGAEVPGPEPAFHLDRRIRLQRLRPLHGQNRVGFKLCNSSG    213
delta-ENaC_Nanger_dama                ARESIYSLYRFNFSDSRDALGAKVPGPEPAFQLDRRIHLQRLRPLDGQNRVGFKLCNSTG    213
delta-ENaC_Kobus_leche                ARENIYSLYRFNFSDSRDALGAEVPGPEPAFQLDRRIYLQRLRPLDGQNRVGFKLCNSTG    213
delta-ENaC_Capra_hircus               ARENIYSLYRFNFSDSKDALGAEVPGPEPAFQLDRRIHLQRLRPLDGQNRVGFKLCNSTG    213
delta-ENaC_Ovis_aries                 ARENIYSLYRFNFSDSRDALGAEVPGPEPAFQLDRRIHLQRLRPLDGQNRVGFKLCNSTG    213
delta-ENaC_Ovis_canadensis            ARENIYSLYRFNFSDSRDALGAEVPGPEPAFQLDRRIHLQRLRPLDGQNRVGFKLCNRTG    213
delta-ENaC_Oreamnos_americanus        ARENIYSLYRFNFSDSRDALGAEVLGPEPAFQLDRRIHLQRLRPLDGQNRVGFKLCNSTG    212
delta-ENaC_Hippotragus_niger          ARENIYSLYRFNFRDSRDALGAEAPGPEPAFQLDRSIHLQRLRPLDGQNRVGFKLCNSTG    213
delta-ENaC_Damaliscus_lunatus         ARENIYSLYRFNFSDSRDALGAEVPGPEPAFQLDRRIHLQRLRPLDGRNRVGFKLCNSTG    213
delta-ENaC_Sus_scrofa                 ARENIYSLYGFNFTESEEDPGAEALGSEPAFQLDRGIRLQRLSPLGGQNRVGFRLCNRTG    211
delta-ENaC_Vicugna_pacos              AQENIYSLYGVSVSKTQDPLAAEDLGPEPHFQLDRGVRLQKLSPLGGQNRVGFRLCNSSG    202
delta-ENaC_Equus_callabus             AQENIYSLYKFNFSESWEGPFAEDLGPEPSFQLDRGIRLQRLNHPGRQSKVGFRLCNSTG    227
                                      *:*.* *** ... .      *     *  *:*** : **:*     : :***:*** :.

delta-ENaC_Homo_sapiens               GDCFYRGYTSGVAAVQDWYHFHYVDILALLPAAWEDSHGSQDGHFVLSCSYDGLDCQARQ    274
delta-ENaC_Hippopotamus_amphibius     SDCIQRAYSSGVAATREWYRFHYVNILASLPATHKDSRHSHGSRFILSCRYDGQDCQARH    274
delta-ENaC_Tragulus_javanicus         GDCVQRAYSSGVLAAREWYRFHYINILALLPATHKD---SPSSRFILSCLYDGQDCYVQH    271
delta-ENaC_Tragulus_kanchil           GDCVQRAYSSGVLAAREWYRFHYINILALLPATHKD---SPSSRFILSCLYDGQDCYVQH    256
delta-ENaC_Antilocapra_americana      GDCVQRAYSSGVVATREWYHFHYINILALLPAAHED---SHSSHFVFSCRYDDRDCHAQH    268
delta-ENaC_Giraffa_camelopardalis     GDCVQQAYSSGVVAAREWYRFHYINILALLPAAHED---SHGSHFVFSCRYDDRDCHAQH    270
delta-ENaC_Giraffa_tippelskirchi      SDCVQQAYSSGVVAAREWYRFHYINILALLPAAHED---SHGSHFVFSCRYDDRDCHAQH    268
delta-ENaC_Capreolus_pygargus         GDCVQRAYSSGMVAAQEWYRFHYINILALLPAAHED---SHSSHFVFSCRYDDRDCHAQH    270
delta-ENaC_Cervus_elaphus             GDCVQQAYSSGIIAAQEWYRFHYINILALLPTAHED---SHGSHFVFSCRYDDRDCHAQH    270
delta-ENaC_Moschus_moschiferus        GDCVQRAYSSGVVAAREWYRFHYMNILALLPAAHEE---SHGSHFVFSCRYNGRDCHAQH    265
delta-ENaC_Moschus_berezovskii        GDCVQRAYSSGVVAAREWYRFHYMNILALLPAAHEE---SHGSHFVFSCRYDGRDCHAQH    265
delta-ENaC_Bos_grunniens              GDCVERAYSSGVVAAREWYRFHYINILALLPAAHED---SHGSHFVFSCRYDDRDCHAQH    270
delta-ENaC_Bos_taurus                 GDCVERAYSSGVVAAREWYRFHYINILALLPAAHED---SHGSHFVFSCRYDDRDCHAQH    270
delta-ENaC_Bubalus_bubalis            GDCVERAYSSGVVAAREWYRFHYINILALLPAAHED---SHGSHFVFSCRYDDRDCHAQH    270
delta-ENaC_Nanger_dama                GDCVQRAYSSGVVAAREWYRFHYINILALLPADHED---SHDSHFVFSCRYDDQDCHAQH    270
delta-ENaC_Kobus_leche                GDCVQRAYSSGVVAAREWYRFHYINILALLPAAHED---SHGSHFVLSCRYDDRDCHARH    270
delta-ENaC_Capra_hircus               GDCVQRAYSSGVVAAREWYRFHYINILALLPAAHED---SHGSHFVFSCRYDDRDCHARH    270
delta-ENaC_Ovis_aries                 GDCVQRAYSSGVVAAREWYRFHYINILALLPAAHED---SHGSHFVFSCRYDDRDCHARH    270
delta-ENaC_Ovis_canadensis            GDCVQRAYSSGVVAAREWYRFHYINILALLPAAHED---SHGSHFVFSCRYDDRDCHARH    270
delta-ENaC_Oreamnos_americanus        GDCVQRAYSSGVVAAREWYRFHYINILALLPAAHED---SHGSHFVFSCRYDDRDCHARH    269
delta-ENaC_Hippotragus_niger          GDCVQRAYSSGVVAAREWYHFHYINILALLPAAHED---SHGSHFVFSCRYDDQDCHARH    270
delta-ENaC_Damaliscus_lunatus         GDCVQRAYSSGVVAAREWYRFHYINILALLPAAHED---SHGSHFVFSCRYDDRDCHARH    270
delta-ENaC_Sus_scrofa                 GDCVHRAYSSGVTAAQEWYRFHYVNILALLPNAPEDSHRSHGGHLVSSCRYDGQDCQAGH    271
delta-ENaC_Vicugna_pacos              GDCIHRAYSSGVTAAREWYRFHYVNILALLPPAHEDHHRSLGSRFIFSCLYDGQDCQARH    262
delta-ENaC_Equus_callabus             GDCFYRVYSSGVTAAREWYRFHYMNILALLPPTREDSHRSHGGHFVFSCRYNGRDCQARH    287
                                      .**. : *:**: *.::**:***::*** **   ::   * ..::: ** *:. ** . :

delta-ENaC_Homo_sapiens               FRTFHHPTYGSCYTVDGVWTAQRPGITHGVGLVLRVEQQPHLPLLSTLAGIRVMVHGRNH    334
delta-ENaC_Hippopotamus_amphibius     FQTSQHPTYGSCYTFNGVWAAQRPGVTHGISLVLSTERQDHLPLLSTEAGVKVMIHRRGH    334
delta-ENaC_Tragulus_javanicus         LQASHHPTYGSCYTFNGVWAAQRPGITHGISLVLRAEQQEHLPLLSMKASVQLIIHPRGH    331
delta-ENaC_Tragulus_kanchil           LQASHHPTYGSCYTFNGVWAAQRPGITHGISLVLRAEQQEHLPLLSMKASVQLIIHPRGH    316
delta-ENaC_Antilocapra_americana      FQTSHHPTYGSCYTFNGVWAAQRPGVTHRISLVLRAEQQDHLPLLSTRAGIKVMIHPQDH    328
delta-ENaC_Giraffa_camelopardalis     FQTSHHPTYGSCYTFNGVWAAQRPGVTHRISLVLRAEQQDHLPLLSTKAGIKVMIHPQDH    330
delta-ENaC_Giraffa_tippelskirchi      FQTSHHPTYGSCYTFKGVWAAQRPGVTHRISLVLRAEQQDHLPLLSTKAGIKVMIHPQDH    328
delta-ENaC_Capreolus_pygargus         FQTSHHPTYGSCYTFNSVWATQRPGVTHRISLVLRAEQQDHLPLLSTKAGIKVMIHPQDH    330
delta-ENaC_Cervus_elaphus             FQTSHHPTYGSCYTFNSVWATHRPGVTHRISLVLRAEQQDHLPLLSTKAGIKVMIHPQDH    330
delta-ENaC_Moschus_moschiferus        FQMSHHPTYGSCYTFNGVWAAPQPGVTHRISLVLRAEQQDHLPLLSTKAGIKVMIHPQDH    325
delta-ENaC_Moschus_berezovskii        FQTSHHPTYGSCYTFNGVWAAPQPGVTHRISLVLRAEQQDHLPLLSTKAGIKVMIHPQDH    325
delta-ENaC_Bos_grunniens              FQTSHHPTYGSCYTFNGVWAAQRPGVTHRISLVLRAEQLDHLPLLSTKAGIKVMIHPQDH    330
delta-ENaC_Bos_taurus                 FQTSHHPTYGSCYTFNGVWAAQRPGVTHRISLVLRAEQLDHLPLLSTKAGIKVMIHPQDH    330
delta-ENaC_Bubalus_bubalis            FQTSHHPTYGSCYTFNSVWAAQRPGVTHRISLVLRAEQLDHLPLLSTKAGIKVMIHPQDH    330
delta-ENaC_Nanger_dama                FQTSHHPTYGSCYTFNGVWATQQPGVTHRISLVLKAEQQDHLPLLSTKAGIKVMIHPQDH    330
delta-ENaC_Kobus_leche                FQTSHHPTYGSCYTFNGVWAAQRPGVTHRISLVLRAEQQDHLPLLSTKAGIKVMIHPQDH    330
delta-ENaC_Capra_hircus               FQTSHHPTYGSCYTFNGVWAAQRPGVTHRISLVLRAEQQDHLPLLSTKAGIKVMIHPQDH    330
delta-ENaC_Ovis_aries                 FQTSHHPTYGSCYTFNGVWAAQRPGVTHRISLVLRAEQQDHLPLLSTKAGIKVMIHPQDH    330
delta-ENaC_Ovis_canadensis            FQTSHHPTYGSCYTFNGVWATQRPGVTHRISLVLRAEQQDHLPLLSTKAGIKVMIHPQDH    330
delta-ENaC_Oreamnos_americanus        FQTSHHPTYGSCYTFNGVWAAPRPGVTHRISLVLRAEQQDHLPLLSTKAGIKVMIHPQDH    329
delta-ENaC_Hippotragus_niger          FQTSHHPTYGSCYTFNGVWAAQRPGVTHRISLVLKAEQQDHLPLLSTKAGIKVMIHPQDH    330
```

```
delta-ENaC_Damaliscus_lunatus        FQTSHHPTYGSCYTFNGVWAAQRPGVTHRISLVLKAEQQDHLPLLSTKAGIKVMIHPQDH       330
delta-ENaC_Sus_scrofa                FQTSHHPTYGSCYTFHGIWAAQHPGITHGISLVLRAELQDHLPLLFTEAGIKVLIHRHDQ       331
delta-ENaC_Vicugna_pacos             FQTSHHPTYGSCYTFDGIWATQCPGITHGISLVLRTEQQDHLPLLSTEAGIKIMIHKRDH       322
delta-ENaC_Equus_callabus            FQTFHHPTYGSCYTFNGAWAAQHPGITHGISLVLRTEQQDHLPLLSTENGIKVMIHRRNQ       347
                                     ::  :**********... *::  **:** :.*** .*   *****    .:::::* :.:

delta-ENaC_Homo_sapiens              TPFLGHHSFSVRPGTEATISIREDEVHRLGSPYGHCTAGGEGVEVELLHNTSYTRQACLV       394
delta-ENaC_Hippopotamus_amphibius    APFLEHQGFSVRPGTETTIGIREDEVHRLGSPYGHCTDGTG-VDVRLLYNTSYTRQACLA       393
delta-ENaC_Tragulus_javanicus        TPFLEHQGFSVRPGTETTISIREDEVHRLGSPYGQCVDGAGGTDGRPLYNTSYTRQACLV       391
delta-ENaC_Tragulus_kanchil          TPFLEHQGFSVRPGTETTISIREDEVHRLGSPYGQCVDGAGGTDGRPLYNTSYTRQACLV       376
delta-ENaC_Antilocapra_americana     TPFLEHQGFSIRPGTETTIDIREDEVHRLGSPYGQCMDSIGSVDVQLLYNTSYTRQACLV       388
delta-ENaC_Giraffa_camelopardalis    TPFLEHQGFSIRPGTETTIDIREDEVHRLGSPYGQCMDSTGTVDVQLLYNTSYTRQACLV       390
delta-ENaC_Giraffa_tippelskirchi     TPFLEHQGFSIRPGTETTIDIREDEVHRLGSPYGQCMDSTGTVDVQLLYNTSYTRQACLV       388
delta-ENaC_Capreolus_pygargus        TPFLEHQGFSIRPGTETTIDIREDEVHRLGSPYGQCMDSTGSVDVQLLYNTSYTRQACLV       390
delta-ENaC_Cervus_elaphus            TPFLEHQGFSIRPGTETTIDIREDEVHRLGSPYGQCMDSTGSVDVQLLYNTSYTRQACLV       390
delta-ENaC_Moschus_moschiferus       TPFLEHQGFSIRPGTETTIDIREDEVHRLGSPYGQCMDSTGSVDMQLLYNTSYTRQACLV       385
delta-ENaC_Moschus_berezovskii       TPFLEHQGFSIRPGTETTIDIREDEVHRLGSPYGQCMDSTGSVDVQLLYNTSYTRQACLV       385
delta-ENaC_Bos_grunniens             TPFLEHQGFSIRPGTETTIDIREDEVHRLGSPYGQCMDSTGSVDVQLLYNTSYTRQACLV       390
delta-ENaC_Bos_taurus                TPFLEHQGFSIRPGTETTIDIREDEVHRLGSPYGQCMDSTGSVDVQLLYNTSYTRQACLV       390
delta-ENaC_Bubalus_bubalis           TPFLEHQGFSIRPGTETTIDIREDEVHRLGSPYGQCMVSTGSVDVQLLYNTSYTRQACLV       390
delta-ENaC_Nanger_dama               TPFLEHQGFSIRPGTETTIDIREDEVHRLGSPYGQCMDSTGSVDVQLLYNTSYTRQACLA       390
delta-ENaC_Kobus_leche               TPFLEHQGFSIRPGTETTIDIREDEVHRLGSPYGQCMDSTGSVDVQLLYNTSYTRQACLA       390
delta-ENaC_Capra_hircus              TPFLEHQGFSIRPGTETTIDIREDEVHRLGSPYGQCMDSTGSMDVQLLYNTSYTRQACLA       390
delta-ENaC_Ovis_aries                TPFLEHQGFSIRPGTETTIDIREDEVHRLGSPYGQCMDSTGNMDVQLLYNTSYTRQACLA       390
delta-ENaC_Ovis_canadensis           TPFLEHQGFSIRPGTETTIDIREDEVHRLGSPYGQCMDSTGSMDVQLLYNTSYTRQACLA       390
delta-ENaC_Oreamnos_americanus       TPFLEHQGFSIRPGTETTIDIREDEVHRLGSPHGQCMDSTGSMDVQLLYNTSYTRQACLA       389
delta-ENaC_Hippotragus_niger         TPFLEHQGFSIRPGTETTIDIREDEVHRLGSPYGQCMDSTGSMDVQLLYNTSYTRQACLA       390
delta-ENaC_Damaliscus_lunatus        TPFLEHQGFSIRPGTETTIDIREDEVHRLGSPYGQCMDSTSSMDVQLLYNTSYTRQACLA       390
delta-ENaC_Sus_scrofa                TPFLEHQGFSIRPGTETTIGIREDEVHRLGSPYGHCTHSAGSVDVRLLYKASYTKQACLV       391
delta-ENaC_Vicugna_pacos             TPFLEHRGFSVRPGTETTIGIREDEVHWLGSPYGQCTDSTEGVDVPLLYNASYTMQACLV       382
delta-ENaC_Equus_callabus            TPFLEHRGFSIRPGTETTIGIREDEVHRLGSPYGHCSDGMEAVDVPLLYNATYTMQACLV       407
                                     :*** *:.**:*****:**.******* ****:*:*  .    :   *::::** ****.

delta-ENaC_Homo_sapiens              SCFQQLMVETCSCGYYLHPLPAGAEYCSSARHPAWGHCFYRLYQDLETHRLPCTSRCPRP       454
delta-ENaC_Hippopotamus_amphibius    SCFQQLMVETCSCGYFFYPLPAGAQYCSYTRHPAWGHCFHRLYKDLQTHRLPCASRCPQL       453
delta-ENaC_Tragulus_javanicus        SCFQQLMVETCSCGYFFYPLPAGAVYCSYGRHPAWGHCFHRLYQKLEAQQLPCATRCPRP       451
delta-ENaC_Tragulus_kanchil          SCFQQLMVETCSCGYFFYPLPAGAVYCSYGRHPAWGHCFHRLYQKLEAQQLPCATRCPRP       436
delta-ENaC_Antilocapra_americana     SCFQQLMVETCSCGYFFYPLPAEAEYCSYTRHPAWGHCFHHLYQKLKTHQLSCTTRCPRP       448
delta-ENaC_Giraffa_camelopardalis    SCFQQLMVETCSCGYFFYPLPAGAEYCSYMRHPAWGHCFHRLYQKLKTHQLPCATRCPRP       450
delta-ENaC_Giraffa_tippelskirchi     SCFQQLMVETCSCGYFFYPLPAGAEYCSYMRHPAWGHCFHRLYQKLKTHQLPCATRCPRP       448
delta-ENaC_Capreolus_pygargus        SCFQQLMVQTCSCGYFFYPLPAGAEYCSYMRHPAWGHCFHHLYQKLKTHQLPCTTRCPQP       450
delta-ENaC_Cervus_elaphus            SCFQQLMVQTCSCGYFFYPLPAGAEYCSYMRHPAWGHCFHHLYQKLKTHQLPCTTRCPQP       450
delta-ENaC_Moschus_moschiferus       SCFQQLMVETCSCGYFFYPLPAGAEYCSYMRHPAWGHCFHRLYQKLKTHQLPCTTRCPRP       445
delta-ENaC_Moschus_berezovskii       SCFQQLMVETCSCGYFFYPLPAGAEYCSYMRHPAWGHCFHRLYQKLKTHQLPCTTRCPRP       445
delta-ENaC_Bos_grunniens             SCFQHLMVETCSCGYFFYPLPAGAEYCSYMRHPAWGHCFHHLYQKLKTHQLPCTTRCPRP       450
delta-ENaC_Bos_taurus                SCFQHLMVETCSCGYFFYPLPAGAEYCSYMRHPAWGHCFHHLYQKLKTHQLPCTTRCPRP       450
delta-ENaC_Bubalus_bubalis           SCFQHLMVETCSCGYFFYPLPAGAEYCSYMRHPAWGHCFHHLYQKLKTHQLPCTTRCPRP       450
delta-ENaC_Nanger_dama               SCFQHLMVETCSCGYFFYPLPAGAEYCSYMRHPAWGHCFHRLYQKLKTHQLPCTTRCPRP       450
delta-ENaC_Kobus_leche               SCFQHLMVETCSCGYFFYPLPAGAEYCSYMRHPAWGHCFHRLYQKLKTHQLPCNTRCPRP       450
delta-ENaC_Capra_hircus              SCFQHLMVETCSCGYFFYPLPVGAEYCSYMRHPAWGHCFHRLYQKLKTHQLPCTTRCPRP       450
delta-ENaC_Ovis_aries                SCFQHLMVETCSCGYFFYPLPVGAEYCSYMRHPAWGHCFHRLYQKLKTHQLPCTTRCPRP       450
delta-ENaC_Ovis_canadensis           SCFQHLMVETCSCGYFFYPLPVGAEYCSYMRHPAWGHCFHRLYQKLKTHQLPCTTRCPRP       450
delta-ENaC_Oreamnos_americanus       SCFQHLMVETCSCGYFFYPLPVGAEYCSYMRHPAWGHCFHHLYQKLKTHQLPCTTRCPRP       449
delta-ENaC_Hippotragus_niger         SCFQHLMVETCSCGYFFYPLPAGAEYCSYMRHPAWGHCFHRLYQKLKTHQLPCTTRCPRP       450
delta-ENaC_Damaliscus_lunatus        SCFQHLMVETCSCGYFFYPLPVGAEYCSYMRHPAWGHCFHRLYQKLKTHQLPCTIRCPRP       450
delta-ENaC_Sus_scrofa                SCFQQLMVDTCSCGYYLYPLPAGAEYCSYTRHPAWGHCFYRLYQDLETHRLPCTSRCPRP       451
delta-ENaC_Vicugna_pacos             SCFQQLMVQTCSCGYYFYPLPAGAEYCSYVRHPAWGHCFYRLYRDLGTHRLPCASRCPRP       442
delta-ENaC_Equus_callabus            SCFQQLMVETCSCGYYLYPLPVGAEYCSYTRHPAWGHCFYRLYRDLETHRLPCASRCPRP       467
                                     ****:***:******:::***. * ***  *********::**:.* :::* *  ***:

delta-ENaC_Homo_sapiens              CRESAFKLSTGTSRWPSAKSAGWTLATLGEQGLPHQSHR---------------------       493
delta-ENaC_Hippopotamus_amphibius    CRESSYKLSAGTSRWPSSKSAGWILALLDTPGHRSPRPGRAQL-----------------       496
delta-ENaC_Tragulus_javanicus        CRESSYKLSAGTSRWPSSKSADWVLAVLGEPGPRSPSPS------WVQP-----------       494
delta-ENaC_Tragulus_kanchil          CRESSYKLSAGTSRWPSSKSADWVLAVLGEPGPRSPSPS------WVQP-----------       479
delta-ENaC_Antilocapra_americana     CRESSYKLSAGTSRWPSSTSSDWVLAVLGEPSRRNPWPPSASI-----------------       491
delta-ENaC_Giraffa_camelopardalis    CRESSYKLSAGTSRWPSSTSADWVLAVLGEPSGRNPWLPSASV-----------------       493
delta-ENaC_Giraffa_tippelskirchi     CRESSYKLSAGTSRWPSSTSADWVLAVLGEPSGRNPWPPSASV-----------------       491
delta-ENaC_Capreolus_pygargus        CRESSYKLSAGTSRWPSSTSADWVLAVLGEPSRRNPWPPSASI-----------------       493
delta-ENaC_Cervus_elaphus            CRESSYRLSAGTSRWPSSTSADWVLAVLGETSRRNPWPPSASI-----------------       493
delta-ENaC_Moschus_moschiferus       CRESSYKLSAGTSRWPSSTSADWVLAVLGEPSRRNPWVPSASV-----------------       488
delta-ENaC_Moschus_berezovskii       CRESSYKLSAGTSRWPSSTSADWVLAVLGEPSRRNPWVPSASV-----------------       488
delta-ENaC_Bos_grunniens             CRESSYKLSAGTSRWPSSTSADWVLAVLGEPSRRNPWPSSASI-----------------       493
delta-ENaC_Bos_taurus                CRESSYKLSAGTSRWPSSTSADWVLAVLGEPSRRNPWPSSASI-----------------       493
delta-ENaC_Bubalus_bubalis           CRESSYKLSAGTSRWPSSTSADWVLAVLGEPSRRNPWPPSASI-----------------       493
delta-ENaC_Nanger_dama               CRESSYKLSAGTSRWPSSTSADWVLAVLSEPSRRNPWPPSASIKSWPLPLPSPSRACTEG       510
delta-ENaC_Kobus_leche               CRESSYKLSAGTSRWPSSTSADWVLAVLGEPSRRNPWPPSASIKSWPLPLPSPSRACTEG       510
delta-ENaC_Capra_hircus              CRESSYKLSAGTSRWPSSTSADWVLAVLGEPSRWNPWPPSASIKSWPLSLPSPSRACTEG       510
delta-ENaC_Ovis_aries                CRESSYKLSAGTSRWPSSTSADWVLAVLGEPSRWNPWPPSASIKSWPLSLPSPSRACTEG       510
delta-ENaC_Ovis_canadensis           CRESSYKLSAGTSRWPSSTSADWVLAVLGEPSRWNPWPPSASIKSWPLSLPSPSRACTEG       510
delta-ENaC_Oreamnos_americanus       CRESSYKLSAGTSRWPSSTSADWVLAVLGEPSRWNPWPPSASIKSWPLSLPSPSRACTEG       509
delta-ENaC_Hippotragus_niger         CRESSYKLSAGTSRWPSSTSADWVLAVLGEPSRRNPWPPSASIKSWPLPLPSPSRACTEG       510
delta-ENaC_Damaliscus_lunatus        CRESSYKLSAGTSRWPSSTSADWVLAVLGEPSRRNPWPPSASIKSWPLPLPSPSRACTEG       510
delta-ENaC_Sus_scrofa                CRESTYKLTAGVSRWPSSKSADWIPAVLDRPGGRQ-SPS---------------------       489
delta-ENaC_Vicugna_pacos             CRESSFRLSAGTSRWPSSKSADWILSVLGKPGH-R-SPS---------------------       479
```

```
delta-ENaC_Equus_callabus              CRESSYKLSAGTSRWPSSKSADWILAVLGKRGHQKPSQS--------------------      506
                                       ****::*:*.*****:.*:.*  : *.   .

                                                                                   TM2?
delta-ENaC_Homo_sapiens                ---------------QRSSLAKINIVYQELNYRSVEEAPVYSVPQLLSAMGSLCSLWFGA      538
delta-ENaC_Hippopotamus_amphibius      -----------PPAKATGTLAKVNIFYQELNYRTVDETPVYSAPQLLSAMGSLWSLWFGS      545
delta-ENaC_Tragulus_javanicus          -----------PSAAVPSSLAKVNIFYQELNYRAVNETPVYSVPQLLSAMGSLWSLWFGS      543
delta-ENaC_Tragulus_kanchil            -----------PSAAVPSSLAKVNIFYQELNYRAVNETPVYSVPQLLSAMGSLWSLWFGS      528
delta-ENaC_Antilocapra_americana       ----------------KISLAKVNIFYQELNYRTVDETPVYSVPQLLSAMGSLWSLWFGS      535
delta-ENaC_Giraffa_camelopardalis      ----------------KISLAKVNIFYQELNYRTVDETPVYSVPQLLSAMGSLWSLWFGS      537
delta-ENaC_Giraffa_tippelskirchi       ----------------KISLAKVNIFYQELNYRTVDETPVYSVPQLLSAMGSLWSLWFGS      535
delta-ENaC_Capreolus_pygargus          ----------------KISLAKVNIFYQELNYRMVNETPVYSVSQLLSAMGSLWSLWLGS      537
delta-ENaC_Cervus_elaphus              ----------------KISLAKVNIFYQELNYRMVNETPVYSVSQLLSAMGSLWSLWLGS      537
delta-ENaC_Moschus_moschiferus         ----------------KISLAKVNIFYQELNYRTVDEMPVYSVPQLLSAMGSLWSLWFGS      532
delta-ENaC_Moschus_berezovskii         ----------------KISLAKVNIFYQELNYRTVDEMPVYSVPQLLSAMGSLWSLWFGS      532
delta-ENaC_Bos_grunniens               ----------------NISLAKVNIFYQELNYRTVDETPVYSVPQLLSAMGSLWSLWFGS      537
delta-ENaC_Bos_taurus                  ----------------NISLAKVNIFYQELNYRTVDETPVYSVPQLLSAMGSLWSLWFGS      537
delta-ENaC_Bubalus_bubalis             ----------------NISLAKVNIFYQELNYRTVDETPVYSVPQLLSAMGSLWSLWFGS      537
delta-ENaC_Nanger_dama                 LTSRSGAQPLSPAPCPRISLAKVNIFYQELNYRTVDETPVYSVPQLLSAMGSLWSLWFGS      570
delta-ENaC_Kobus_leche                 PASRSRAQTLSPAPCPRISLAKVNIFYQELNYRTVDETPVYSVPQLLSAMGSLWSLWFGS      570
delta-ENaC_Capra_hircus                PTGRSGAQPLSPAPCPRISLAKVNIFYQELNYRTVDETPVYSVPQLLSAMGSLWSLWFGS      570
delta-ENaC_Ovis_aries                  PTGRSGAQPLSPAPCPRISLAKVNIFYQELNYRTVDETPVYSVPQLLSAMGSLWSLWFGS      570
delta-ENaC_Ovis_canadensis             PTGRSGAQPLSPAPCPRISLAKVNIFYQELNYRTVDETPVYSVPQLLSAMGSLWSLWFGS      570
delta-ENaC_Oreamnos_americanus         PTGRSGAQPLSPAPCPRISLAKLNIFYQELNYRTVDETPVYSVPQLLSAMGSLWSLWFGS      569
delta-ENaC_Hippotragus_niger           PTSRSGAQPLSPAPCPRISLAKVNIFYQELNYRTVDETPVYSVPQLLSAMGSLWSLWFGS      570
delta-ENaC_Damaliscus_lunatus          PTNRSGAQPLSPAPCPRISLAKVNIFYQELNYRTVDETPVYSVPQLLSAMGSLWSLWFGS      570
delta-ENaC_Sus_scrofa                  ---------------PRSRLAKVNIFYRELNYRTVDEMPVYSVPQLLSAMGSLWSLWFGS      534
delta-ENaC_Vicugna_pacos               ---------------PRSNLAKVNILYQELNYRTVDETPIYSVPRLLSAMGSLWSLWFGS      524
delta-ENaC_Equus_callabus              ---------------PRSNIAKMNIFYQELNYRTVDETPVYSVPQLLSAMGSLWSLWFGS      551
                                                          :**:**.*:***** *:* *:**. :******** ***:*:

delta-ENaC_Homo_sapiens                SVLSLLELLELLLDASALTLVLGGRRLRRAWFSWPRASPASGASSIKPEASQMPPPAGGT      598
delta-ENaC_Hippopotamus_amphibius      SVLSVVEVLELLLDAAALALLLCCRRLRGA-RGQPGTATGAPAPSQRP--AGCPAAAGMT      602
delta-ENaC_Tragulus_javanicus          SVLSVVEVLELLLDATALALLLGHHWLCGA-GGRPRAAPGVPAPSQRL--AGGPVAASPR      600
delta-ENaC_Tragulus_kanchil            SVLSVVEVLELLLDATALALLLGHHWLCGA-GGRPRAAPGVPAPSQRL--AGGPVAASPR      585
delta-ENaC_Antilocapra_americana       SVLSVVEVLELLLDAMALTLLLCCRWLRGS-RGHPRVATRVPSSSQRP--ASGPVAARTM      592
delta-ENaC_Giraffa_camelopardalis      SVLSVVEVLELLLDAMALTLLLCCRWLRGS-RRQPGAATRVPSPSQRP--ASGPVAAGTT      594
delta-ENaC_Giraffa_tippelskirchi       SVLSVVEVLELLLDAMALTLLLCCRWLRGS-RRQPGAATRVPSPSQRP--ASGPVAAGTT      592
delta-ENaC_Capreolus_pygargus          SVLSVVEVLELLLDAMALTLLLCCRRLRGS-QGQPRAAARVPPPSQRP--ASGPVAAGTT      594
delta-ENaC_Cervus_elaphus              SVLSVVEVLELLLDAMALTLLLCCRWLRRS-QGQPRAAMRVPPPSQRP--ASGPVAAGTT      594
delta-ENaC_Moschus_moschiferus         SVLSVVEVLELLLDAMALTLLLCCRWLRGS-QRQPRAATRVPPPSQRP--ASGPVAAGTT      589
delta-ENaC_Moschus_berezovskii         SVLSVVEVLELLLDAMALTLLLCCRWLRGS-QRQPRAATRVPPPSQRP--ASGPVAAGTT      589
delta-ENaC_Bos_grunniens               SVLSVVEVLELLLDAIALTLLLCCRWLCGS-RGQPRAATRVHPPSQRP--ASGPVAADTT      594
delta-ENaC_Bos_taurus                  SVLSVVEVLELLLDAIALTLLLCCRWLCGS-RGQPRAATRVHPPSQRP--ASGPVAADTT      594
delta-ENaC_Bubalus_bubalis             SVLSVVEVLELLLDAMALTLLLCCRWLRGS-RGQPRAATRVHPPSQRP--ASGPVAAGTT      594
delta-ENaC_Nanger_dama                 SVLSVVEVLELLLDAMALTLLLCCRWLRGS-WGQPRAATSVPPPSQRP--ASGPVPAGTT      627
delta-ENaC_Kobus_leche                 SVLSVVEVLELLLDAMALTLLLCCRWLRGS-RGQPRAATRVPPPSQRP--ASGPVAAGTT      627
delta-ENaC_Capra_hircus                SVLSVVEVLELLLDTMALALLLCCRWLRGS-QGQPRAATRVPPPSQRP--ASRPVAAGT-      626
delta-ENaC_Ovis_aries                  SVLSVVEVLELLLDTMALALLLCCRWLRGS-QGQPRAATRVPPPSQRP--ASGPVAAGTT      627
delta-ENaC_Ovis_canadensis             SVLSVVEVLELLLDTMALALLLCFRWLRGS-QGQPRAATRVPPPSQRP--ASGPVAAGTT      627
delta-ENaC_Oreamnos_americanus         SVLSVVEVLELLLDTMALALLLCCRWLRRS-QGQPRVATRVPPPSQRP--ASGPVATGTT      626
delta-ENaC_Hippotragus_niger           SVLSVVEVLELLLDAMALALLLCCRWLRGS-RGQPRAATRVPPPSQRP--ASGPVAVGTT      627
delta-ENaC_Damaliscus_lunatus          SVLSVVEVLELLLDAMALALLLCCRWLRGS-RGQPRAATRVPPPSQRP--ASGPVAAGTT      627
delta-ENaC_Sus_scrofa                  SVLSVLEVLELLLDAMVLTLLLCCRQLHRA-RGQLGAAPGEPTPSQRP--TCCPAAAGMT      591
delta-ENaC_Vicugna_pacos               SVLSVLEVLELLLDATALTFLLCCRRLHAA-RGQPRAATRAPTPSQRP--ASCPVAGGTS      581
delta-ENaC_Equus_callabus              SVLSVLELLELLLDATALALLLCCRRLRRAWRSQPGAATRASGPRPEA--TPLPSDCGPD      609
                                       ****::*:******: .*:::*  : *  :      .:         .   :  *

delta-ENaC_Homo_sapiens                SDDPEPSGPHLPRVMLPGVL--A-GVSAE------------   624
delta-ENaC_Hippopotamus_amphibius      SHAPGPSWLHLRRGCQGSSRSLG------------------   625
delta-ENaC_Tragulus_javanicus          SHAPGPAAFASHNAAGAPAGVLAAELSQVWAPETPNT----   637
delta-ENaC_Tragulus_kanchil            SHAPGPAAFASHNAAGAPAGVLAAELSQVWAPETPNT----   622
delta-ENaC_Antilocapra_americana       SNTPGPGCLHLPRCCWGFSR----SLS--------------   615
delta-ENaC_Giraffa_camelopardalis      SNTPGPGCLHLPGLQQESRL----RNSAKCGPQRPLTPEPV   631
delta-ENaC_Giraffa_tippelskirchi       SNTPGPGCLHLPGLQQESRL----RNSAKCGPQRPLTPEPV   629
delta-ENaC_Capreolus_pygargus          SNAPGPGCLRLPRCCRDFSR----SLG--------------   617
delta-ENaC_Cervus_elaphus              SNAPGPGCLRLPRCCRDFSR----SLG--------------   617
delta-ENaC_Moschus_moschiferus         SNAPGPGCLHLPRCCRDFSR----SLG--------------   612
delta-ENaC_Moschus_berezovskii         SNAPGPGCLHLPRCCRDFSR----SLG--------------   612
delta-ENaC_Bos_grunniens               SNAPGPGCLHLPRCCRDFSR----SLG--------------   617
delta-ENaC_Bos_taurus                  SNAPGPGCLHLPRCCRDFSR----SLG--------------   617
delta-ENaC_Bubalus_bubalis             SNAPGPGCLHLPRCCRDFGR----SLD--------------   617
delta-ENaC_Nanger_dama                 SNAPGPSCLHLPRCCRDFSR----SLG--------------   650
delta-ENaC_Kobus_leche                 SNALGPSCLHLPQCCQDFSR----SLG--------------   650
delta-ENaC_Capra_hircus                SNALGPSCLHLPRCCRDFSR----SLS--------------   649
delta-ENaC_Ovis_aries                  SNALGPSCLHLPRCCRDFSR----SLG--------------   650
delta-ENaC_Ovis_canadensis             SNALGPSCLHLPRCCRDFSR----SLG--------------   650
delta-ENaC_Oreamnos_americanus         SNALGPSCLHLPRCCRDFSR----SLG--------------   649
delta-ENaC_Hippotragus_niger           SNAPGPGCLHLPRCCRDFSR----SLS--------------   650
delta-ENaC_Damaliscus_lunatus          SNAPGPGCLHLPRCCRDFSR----SLG--------------   650
delta-ENaC_Sus_scrofa                  SNARGPSWPRFP-----------------------------   603
delta-ENaC_Vicugna_pacos               -DVQGPG----------------------------------   587
delta-ENaC_Equus_callabus              --LKCSGTPAGASAEESRL----------------------   626
                                              .
```
