## Supplemental Data 3 for "The evolutionary path of the epithelial sodium channel δ-subunit in Cetartiodactyla points to a role in sodium sensing"

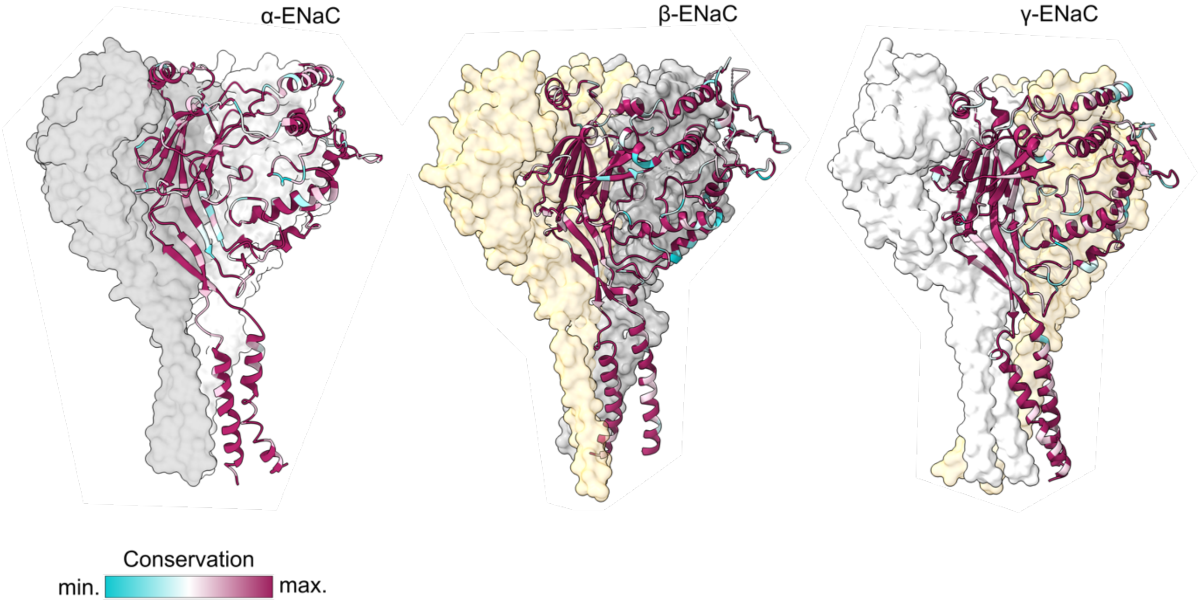


**Conservation of ENaC subunit sequences**. The Cryo-EM derived structure of human ENaC (PDB: 6BQN) (Noreng et al. 2018) is displayed and each ENaC subunit is coloured by the conservation in the multiple sequence alignments provided in Supplemental Data 2. Molecular graphics and analyses were performed with UCSF ChimeraX (Meng et al. 2023), developed by the Resource for Biocomputing, Visualization, and Informatics at the University of California, San Francisco, with support from National Institutes of Health R01-GM129325 and the Office of Cyber Infrastructure and Computational Biology, National Institute of Allergy and Infectious Diseases.
